# WDR82-binding long non-coding RNA *lncEry* controls mouse erythroid differentiation and maturation

**DOI:** 10.1101/2021.07.13.452142

**Authors:** Shangda Yang, Guohuan Sun, Peng Wu, Cong Chen, Yijin Kuang, Zhaofeng Zheng, Yicheng He, Quan Gu, Ting Lu, Caiying Zhu, Fengjiao Wang, Fanglin Gou, Zining Yang, Xiangnan Zhao, Shiru Yuan, Liu Yang, Shihong Lu, Yapu Li, Xue Lv, Fang Dong, Yanni Ma, Jia Yu, Lai Guan Ng, Lihong Shi, Jing Liu, Hui Cheng, Tao Cheng

**Affiliations:** State Key Laboratory of Experimental Hematology, National Clinical Research Center for Blood Diseases, Institute of Hematology & Blood Diseases Hospital, Chinese Academy of Medical Sciences & Peking Union Medical College, Tianjin, China; Center for Stem Cell Medicine, Department of Stem Cell & Regenerative Medicine, Chinese Academy of Medical Sciences, Tianjin, China; Department of Cell Biology, Tianjin Medical University, Tianjin, China; Molecular Biology Research Center, Center for Medical Genetics, Hunan Province Key Laboratory of Basic and Applied Hematology, School of Life Sciences, Central South University, Changsha, China; State Key Laboratory of Medical Molecular Biology, Key Laboratory of RNA Regulation and Hematopoiesis, Department of Biochemistry and Molecular Biology, Institute of Basic Medical Sciences, Chinese Academy of Medical Sciences, School of Basic Medicine Peking Union Medical College, Beijing, China; Singapore Immunology Network, Agency for Science, Technology and Research, Biopolis, 138648 Singapore

## Abstract

Hematopoietic differentiation is controlled by both genetic and epigenetic regulators. Long non-coding RNAs (lncRNAs) have been demonstrated to be important for normal hematopoiesis, but their function in erythropoiesis needs to be further explored. We profiled the transcriptomes of 16 murine hematopoietic cell populations by deep RNA-sequencing and identified a novel lncRNA, *Gm15915*, that was highly expressed in erythroid-related progenitors and erythrocytes. For this reason, we named it *lncEry*. We also identified a novel *lncEry* isoform, which was also the principal transcript that has not been reported before. *LncEry* depletion impaired erythropoiesis, indicating the important role of the lncRNA in regulating erythroid differentiation and maturation. Mechanistically, we found that *lncEry* interacted with WD repeat-containing protein 82 (WDR82) to promote the transcription of *Klf1* and globin genes and thus control the early and late stages of erythropoiesis, respectively. These findings identified *lncEry* as an important player in the transcriptional regulation of erythropoiesis.

## INTRODUCTION

Hematopoietic stem cells (HSCs) are multipotent precursors with the capacity to self-renew and differentiate into all mature blood cell types^1–4^. During hematopoietic differentiation, long-term HSCs (LT-HSCs) differentiate into multiple blood cellular components^5^, including short-term HSCs (ST-HSCs), multipotent progenitor cells (MPPs), committed progenitor cells, and mature blood cells^6,7^. Hematopoiesis is tightly regulated by various regulatory elements, including non-coding RNAs (ncRNAs), to maintain normal biological processes^8,9^.

In mammals, approximately two-thirds of genomic DNA is pervasively transcribed^10^, while most genomic DNA is transcribed into ncRNAs, suggesting that RNA-based regulatory mechanisms might be involved in the complex developmental processes of eukaryotes^11,12^. These ncRNAs can be divided into two main types: small ncRNAs and long non-coding RNAs (lncRNAs). LncRNAs are defined as transcripts of >200 nucleotides with no apparent open reading frames^9,13,14^. Numerous functional lncRNAs have been discovered, including those that have a vital role in mediating hematopoiesis. For example, the oncofetal lncRNA gene *H19* controls the balance between HSC quiescence and activation by regulating Igf2-Igf1r pathway activation^15^, promoting pre-HSC and HSC specification via the demethylation of hematopoietic transcription factors such as Runx1 and Spi1^16^, or by participating in tumorigenesis^17^. *LncHSC-1* and *lncHSC-2* regulate the differentiation of myeloid and T cells, respectively^4^, while *lnc-DC* regulates monocyte-derived DC differentiation through STAT3 binding^18^. Erythropoiesis is also regulated by lncRNAs^19–24^, for example, lncRNA-*ERS*^25^ regulates the terminal differentiation of erythroid cells by promoting erythroid progenitor survival; lncRNA *UCA1* controls erythropoiesis at the proerythroblast stage through the regulation of heme metabolism^26^; and *lnc-EC1* and *lnc-EC6* regulate erythroblast enucleation. Yet, despite such advances in our knowledge lncRNA identification, the function of most lncRNAs in erythropoiesis regulation remains largely unknown.

Erythroid Krüppel-like factor (EKLF; KLF1)^29^ is a zinc-finger hematopoietic transcription factor that plays a global role in regulating the activation of genes in different stages of erythropoiesis^30,31^. The selective expression of *Klf1* promotes erythropoiesis and represses megakaryopoiesis^32,33^. During maturation, nucleated erythrocytes (NuEs) shed their nucleus and progressively gain erythroid characteristics, changing from nucleated erythrocytes to reticulocytes and ultimately mature blood cells. These mature blood cells are regulated by hemoglobins, which, if dysfunctional, can induce hemoglobinopathies such as β-thalassemia or sickle cell anemia^34^. In differentiated erythroid cells, the remote regulatory sequences of the α-globin gene recruit polymerase II and the pre-initiation complex, then bind to transcription factors located in the promoter region to activate α-globin transcription^35^. Despite the depth of our understanding, we still need to ascertain the complexities of the transcriptional mechanisms regulating *Klf1* and globin expression.

While many functional lncRNAs are recognized hematopoiesis mediators, the lncRNAs that regulate erythroid differentiation need clarification. Thus, we aimed to search for previously unidentified functional lncRNA(s) that play a role in erythroid differentiation by employing high-throughput RNA sequencing (RNA-seq) approaches. We further investigated the mechanisms of IncRNA in the regulation of erythropoiesis by promoting the transcriptional activation of KLF1 and globin genes at different stages of erythropoiesis.

## RESULTS

### *Gm15915* is highly expressed in an erythroid lineage

Previous studies have confirmed the involvement of lncRNAs in many biological processes, including lineage differentiation^4,36–38^. To identify novel lncRNAs with biological relevance, we isolated 16 hematopoietic cell subsets from the bone marrow (BM) of C57BL/6 mice by fluorescence-activated cell sorting (FACS). These cell types included LT-HSCs, ST-HSCs, MPPs, common lymphoid progenitors (CLPs), common myeloid progenitors (CMPs), granulocyte-macrophage progenitors (GMPs), and megakaryocyte-erythroid progenitors (MEPs). We also isolated 10 mature lineage cell subsets: NK cells, B cells, CD4-T cells, CD8-T cells, monocytes, macrophages, granulocytes, megakaryocytes, and NuEs. Through RNA-seq, we identified 2,250 lncRNAs and 13,168 protein-coding genes. Thus, just a handful of highly expressed lncRNAs constituted the hematopoietic cell landscape (Figure 1A, Supplementary Table 1). Consistent with previous studies^15^, we found that the classical lncRNA *H19* was highly expressed in LT-HSCs (Figure 1A), illustrating the validity of our approach.

**Figure 1.**
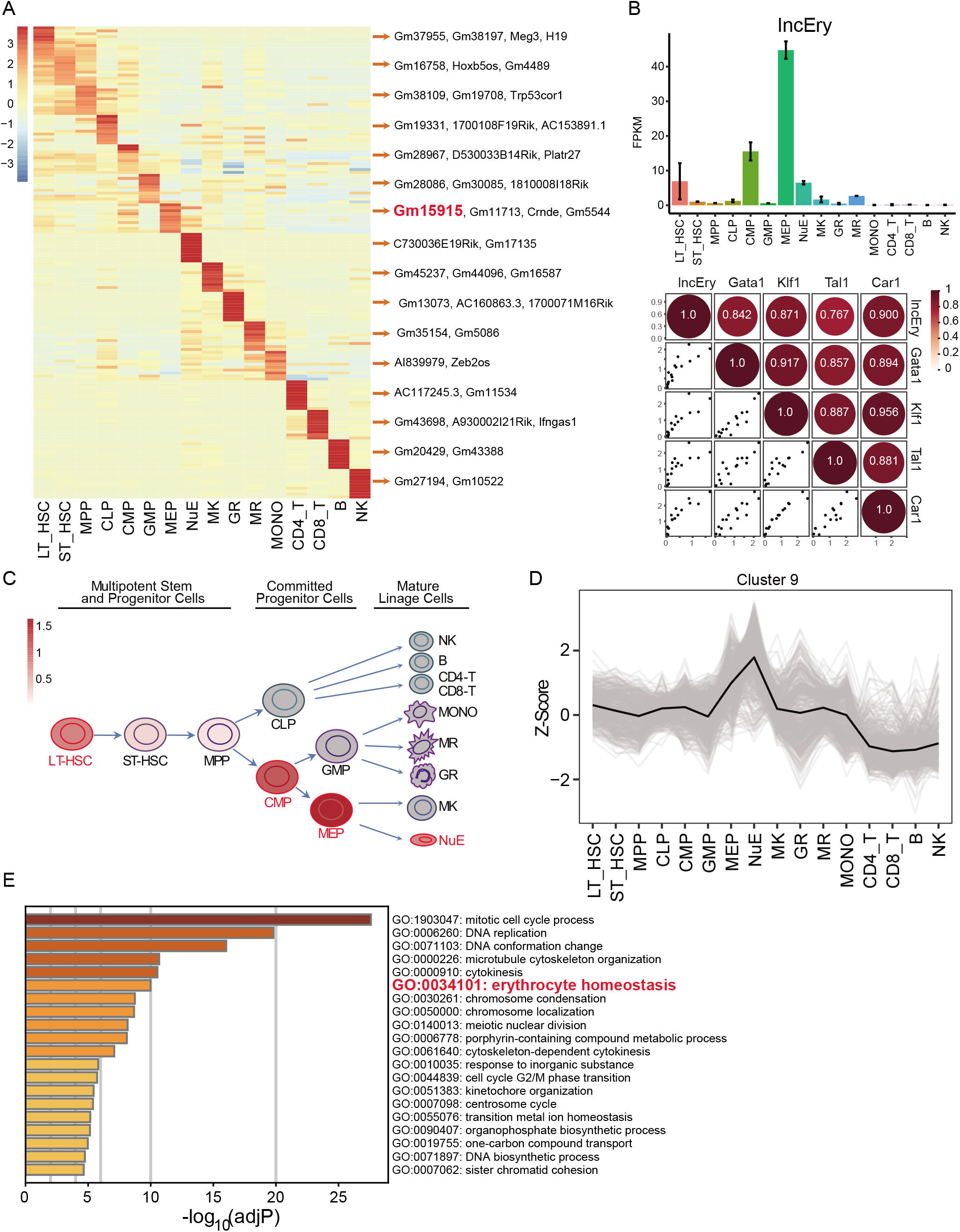
Gm15915 is highly expressed in erythroid lineage. (A) Heat map of lncRNA expression across 16 hematopoietic cell populations. Representative lncRNAs in different cell types are shown on the right. The top 10 lncRNAs are shown in Supplementary Table 1. (B) RNA-seq analysis of *lncEry* expression in the 16 hematopoietic cell populations (upper panel). Correlation analysis of *lncEry* and erythroid lineage related gene expression in the 16 hematopoietic cell populations (lower panel). Each bar represents normalized duplicate samples. (C) Schematic representation of the hematopoietic hierarchy showing the cell types used in this study; the intensity of the red color indicates the level of *lncEry* expression. (D) Line chart showing Cluster 9 gene expression in the 17 hematopoietic cell populations. (E) Gene enrichment analysis showing −log 10 of the uncorrected P-values on the x-axis; darker shading corresponds to a greater amount of enriched genes in each term.

To gain insights into the expression of lncRNAs in erythropoiesis, we decided to focus on the lncRNAs specifically expressed in MEPs, of which *Gm15915* was the most highly expressed (Figure 1A). *Gm15915* was also highly expressed in LT-HSCs, CMPs, and NuEs (Figure 1B, upper panel). *Gm15915* levels positively correlated with the expression of erythroid lineage-development-associated genes, such as *Gata1, Klf1, Tal1*, and *Car1* (Figure 1B, lower panel), suggesting *Gm15915* is an erythroid-lineage-specific lncRNA (Figure 1C). Hence, *Gm15915* was named “*lncEry*”. To investigate *lncEry* further, we reanalyzed its expression in hematopoietic cells using a dataset from a previous study^39^. *lncEry* was highly expressed in LT-HSCs, ST-HSCs, CMPs, MEPs, and NuEs (Figure S1A), confirming our findings. In addition, by analyzing single-cell RNA-seq data from a previous study, we found *lncEry* to be highly expressed in the MEP population (Figure S1B). Next, the expressions of *lncEry* in CMPs, GMPs, and MEPs were compared to determine any correlation with cell fate and found that *lncEry* expression increased as CMPs differentiated into MEPs but not GMPs (Figure S1C).

To further investigate *lncEry* function, we performed unsupervised hierarchical clustering on the expression levels of protein-coding and lncRNA genes. We defined 10 clusters and hypothesized that the genes expressed within the same cluster might have similar functions (Figure S1D). The genes in Cluster 9 were highly expressed in MEPs and NuEs (Figure 1D). We found known erythropoiesis-associated genes, such as *Gata1, Klf1, Tal1*, and *Car1*, and *lncEry* in Cluster 3. Finally, Gene Ontology (GO) enrichment analysis showed that the coding genes in Cluster 3 were significantly enriched for erythrocyte differentiation, supporting the functional role of *lncEry* in erythroid differentiation (Figure 1E). Taken together, our data indicate that *lncEry* is potentially an important regulator of erythroid differentiation.

### LncEry is a *bona fide* long non-coding RNA

To gain insights into the molecular characteristics of *lncEry*, we performed 5’ and 3’ rapid amplification of cDNA ends (RACE) PCR of MEP clones, followed by Sanger sequencing, to identifying *lncEry* transcript isoforms (Figure 2A). Two isoforms (isoform-1: NONMMUG004428.1 and isoform-2: NONMMUG0048.2) are annotated in the NONCODE database, and we discovered a third isoform, isoform-3, which shares four incomplete exons (exons 2-4, and 6 of isoform-1) with the other two (Figure 2B).

**Figure 2.**
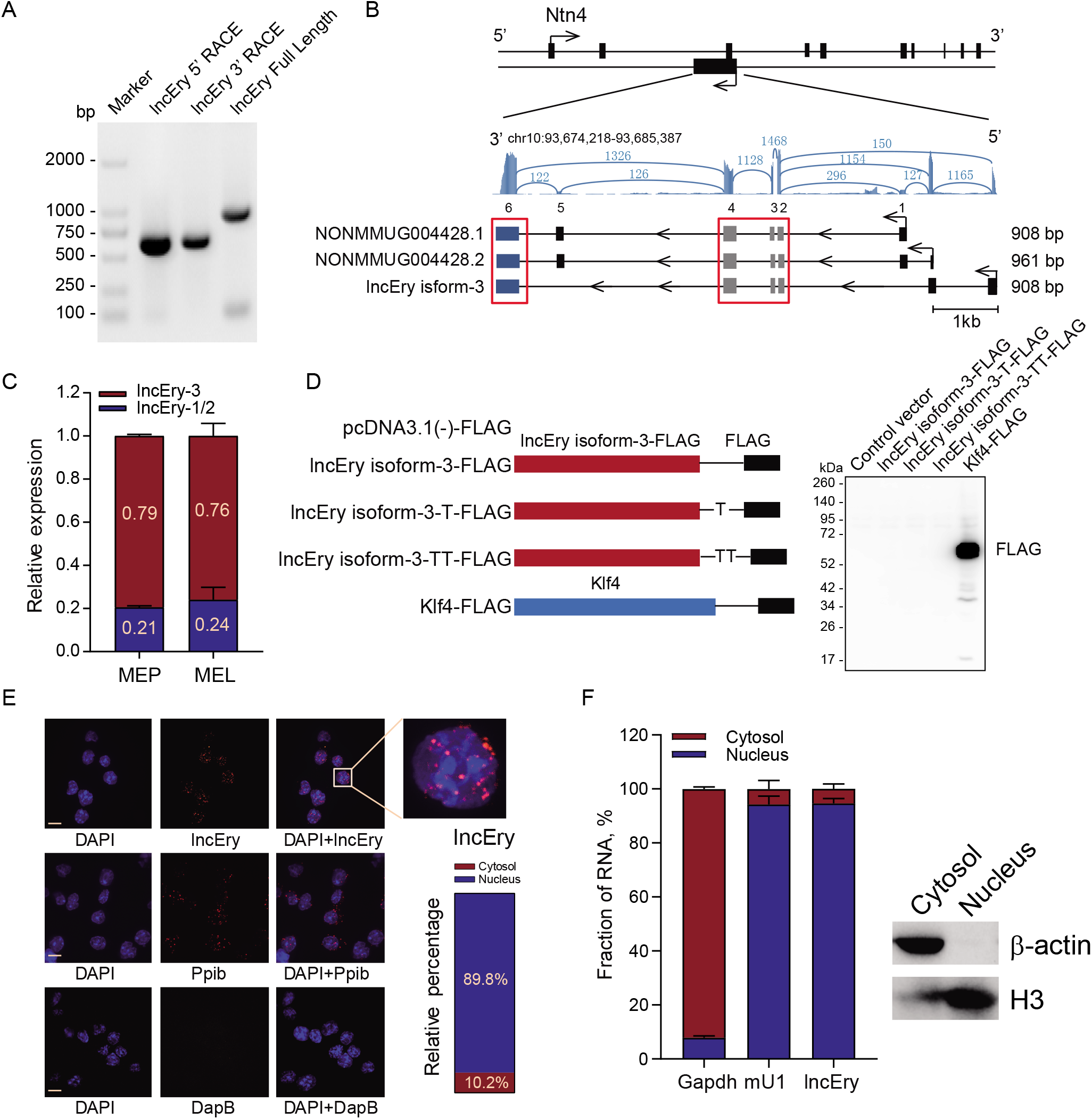
*LncEry* is a *bona fide* long non-coding RNA. (A) 5’ and 3’ RACE assays and gel electrophoresis analysis to detect *lncEry* transcripts in of MEP cells. (B) RNA-seq tracks at *lncEry* loci with different read numbers linked to the different exons of the *lncEry* isoforms. The shared exons of the different isoforms are marked by the red frame. *LncEry* isoform-1: NONMMOUG004428.1; *lncEry* isoform-2: NONMMUG004428.2. (C) Expression of different *lncEry* isoforms in MEP and MEL cells analyzed by qPCR. Error bars represent the mean ± S.D. for biological triplicates. (D) Western blot showing that there was no expression of Flag-tagged lncEry isoform-3 in all three reading frames in MEL cells. Flag-tagged KLF4 was used as a positive control. Full-length *lncEry* isoform-3 was cloned into the eukaryotic expression vector pcDNA3.1(-) with an N-terminal start codon ATG and a C-terminal Flag tag in all three reading frames; these plasmids were transfected into MEL cells separately and analyzed by western blotting.. (E) RNAscope and confocal microscopy analysis of *lncEry* subcellular localization in MEL cells. Ppib and DapB probes were used as positive and negative controls, respectively. The percentage of different *lncEry* subcellular localization points in >50 cells was calculated. Scale bar, 10 μm. (F) The *lncEry* fraction in the cytosol and nucleus. Gapdh and mU1 were used as markers of the cytosolic and nuclear RNA fractions, respectively. Each error bar represents the mean ± S.D. of triplicate experiments. The separation efficiency of each cell component was determined by western blotting.

After analyzing the detailed sequence information (Figure S2A) and the RNA-seq read coverage of the *lncEry* gene in MEPs, we found *lncEry* isoform-3 to be the principal transcript (Figure 2B). Specifically, ~79.5% and ~20.5% of *lncEry* was expressed as isoform-3 and isoform-1/2, respectively. Similarly, 76% of *lncEry* in mouse erythroleukemia cell line (MEL) was expressed as isoform-3 (Figure 2C and Figure S2B, C). Next, we examined the coding capacity of isoform-3. Full-length isoform-3 was inserted into the eukaryotic expression vector pcDNA3.1 with N-terminal start codon ATG and C-terminal Flag tag in all three reading frames, and the results confirmed the non-protein-coding characteristics of isoform-3 (Figure 2D).

The functions of most lncRNAs are restricted to their subcellular localization^40–42^. We thus performed RNAscope assays to identify the location of *lncEry* in MEL cells. Nearly 90% of *lncEry* molecules localized to the nucleus (Figure 2E), which was confirmed by subcellular fractionation assay followed by quantitative PCR (qPCR) (Figure 2F). We presumed that the nuclear location of this lncRNA indicated its involvement in transcriptional regulation^43^.

To prove this hypothesis, we transfected MEL cells with small interfering (si)RNAs targeting *lncEry* and analyzed the expression of the genes found within 1 Mbp upstream and downstream of the *lncEry* locus on chromosome 10. *lncEry* downregulation not only affected the expression levels of adjacent genes, such as *Ntn4* and *Ccdc38*, but also influenced (to some extent) the expression of genes located >100 Kbps distant, e.g., *Hal* and *Lta4h* (Figure S2D). We thus speculated that *lncEry* has transcription regulatory capacity.

### Erythroid differentiation is impaired upon *lncEry* depletion in HSPCs

To determine the functional impact of *lncEry* on HSPCs, we transduced cKit^+^ cells with a lentivirus carrying GFP-fused *lncEry* short hairpin (sh)RNA, then performed *in vitro* colony forming unit (CFU) and *in vivo* transplantation assays (Figure S3A). First, we confirmed that the two shRNA constructs exhibited high-knockdown efficiencies at the mRNA level (Figure S3B). The CFU assays showed that *lncEry*-knockdown decreased the number of CFU-GM colonies by 32% on average and more potently decreased the number of BFU-E and CFU-GEMM colonies (74% and 66% on average, respectively) (Figure S3C). In addition, colony sizes were significantly decreased upon *lncEry* knockdown (Figure S3D). These results are consistent with our hypothesis that *lncEry* is involved in erythroid differentiation.

Next, we infected donor cells with *lncEry*-shRNA-carrying lentiviruses, and after 48 h of culture, achieved transduction efficiencies of approximately 94%, 84%, and 40% for control, shRNA-1, and shRNA-2, respectively (Figure S3E). We then transplanted the transduced cells (without sorting) into lethally irradiated mouse recipients (Figure S3A); 21 days after transplantation, two recipients from the *lncEry* shRNA-1 group died, and the remaining three mice in the *lncEry* shRNA-1 group showed pale paws and were moribund, indicative of severe anemia. The recipients of the *lncEry* shRNA-1 and *lncEry* shRNA-2 had decreased numbers of white blood cells (WBCs) and red blood cells (RBCs) (Figure S3F), as well as very low hemoglobin levels compared with the controls (Figure S3F). Finally, the *lncEry* knockdown animals showed a decrease in the percentage of GFP^+^ cells in the peripheral blood (PB) and BM (Figure S3G and S3H) and in the percentage of reticulocytes and RBCs in the GFP^+^ cells within the BM (Figure S3I). We thus considered that *lncEry* is involved in erythroid differentiation from HSPCs.

### Erythroid differentiation is impaired in *lncEry* Δ/Δ mice

To study the function of *lncEry* in erythroid differentiation, we generated *lncEry^fl/fl^* (*flox/flox*) mice (Figure S4A). Then, we generated Mx1-Cre; *lncEry^fl/fl^* (Δ/Δ) mice, in which *lncEry* deletion could be induced by poly I:C administration^44,45^. We analyzed the expression of *lncEry* isoforms in the BM cells of both lncEryfl/fl and Δ/Δ mouse strains: excision of exons 1/2 of *lncEry* isoform-3 strongly decreased the expression of isoform-3 as well as the other two *lncEry* isoforms (Figure 3A).

**Figure 3.**
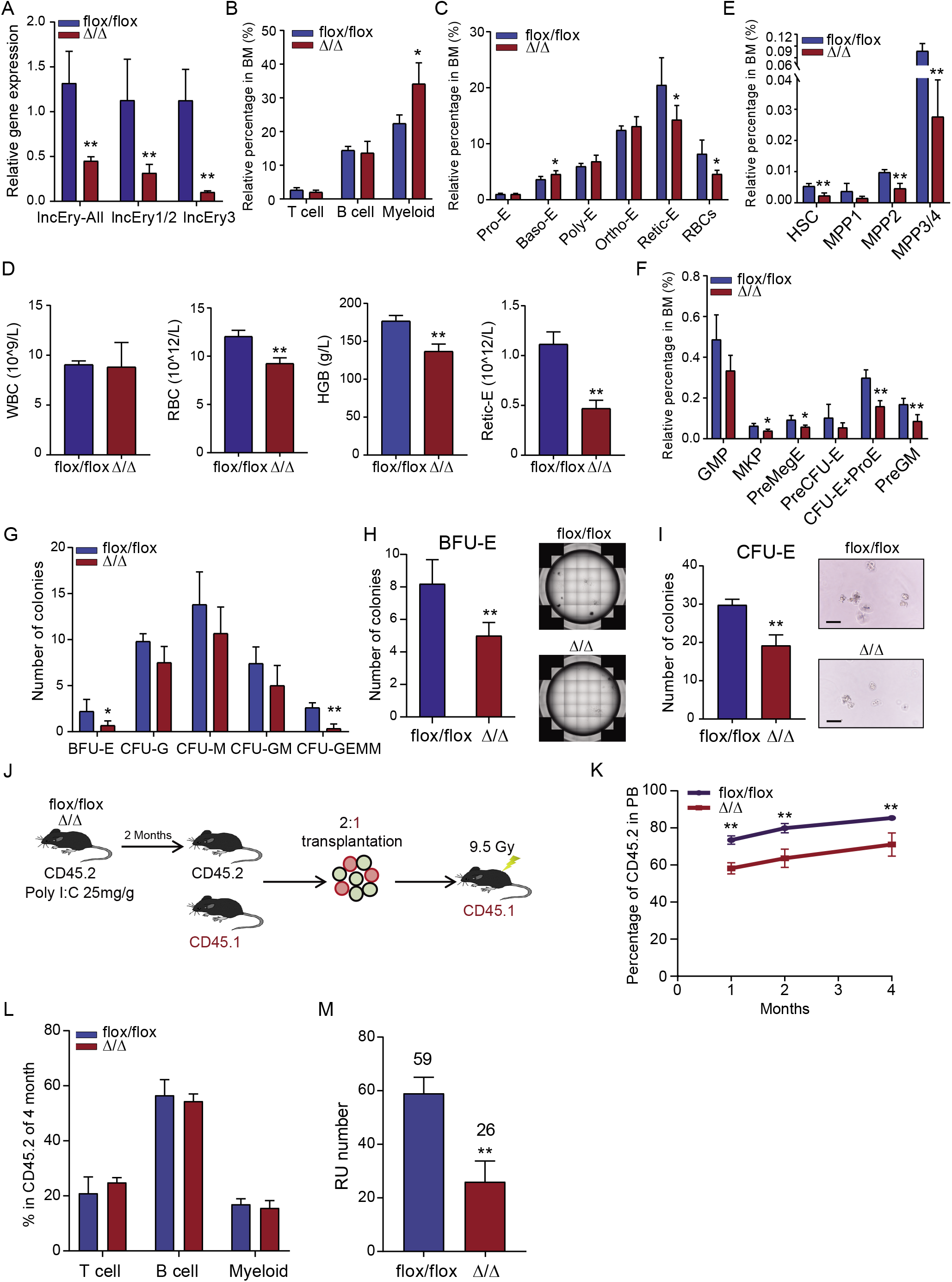
Erythroid differentiation is impaired in *lncEry* Δ/Δ mouse. (A) qPCR analysis of *lncEry* isoform expression in *flox/flox* or Δ/Δ bone marrow (BM) cells. (B-C) Percentage of indicated cell populations in *flox/flox* or Δ/Δ mice BM. Each bar represents the mean ± S.D. of triplicate experiments. *P < 0.05, **P < 0.01, one-way ANOVA. (D) Absolute numbers or concentrations of the indicated items in the peripheral blood of *flox/flox* or Δ/Δ mice. (E-F) Percentage of indicated cell populations in the BM of *flox/flox* or Δ/Δ mice. (G) 1 × 10^4^ BM cells from *flox/flox* or Δ/Δ mice were cultured for 10-14 days in colony forming unit (CFU) assays in complete methylcellulose-based medium, and colony numbers were counted. (H) BFU-E CFU assays of 1 × 10^4^ *flox/flox* or Δ/Δ mice BM cells cultured in methylcellulose-based medium with EPO cytokine supplementation for 10-14 days. Representative images from triplicate experiments are shown. (I) CFU-E colony assays of 500 *flox/flox* or Δ/Δ mice MEP cells cultured in methylcellulose-based medium with EPO cytokine for 48 h. Representative images from triplicate experiments are shown. Scale bar, 50 μm. (J) Experimental flow-chart of competitive transplantation; mice were treated with poly I:C 25 mg/g three times every other day before transplantation. (K) Percentage of CD45.2^+^ cells in the peripheral blood (PB) of recipient (CD45.1^+^) mice. (L) Percentage of CD45.2^+^ cells in indicated populations 4 months after transplant. (M) Repopulating units (RU) of donor cells calculated after 4 months reconstitution. Each bar represents the mean ± S.D. for triplicate experiments. *P < 0.05, **P < 0.01.

To monitor the effects of *lncEry* on the hematopoietic system, we analyzed BM cells from *flox/flox* and Δ/Δ mice by flow cytometry. We found that the Δ/Δ mice had normal BM cellularity but an increased percentage of myeloid cells (Figure 3B and Figure S4B) and a decreased percentage of reticulocytes (Retic-E) and RBCs in BM and Ter119^+^ cells^46,47^ (Figure 3C and Figure S4C-E). These findings are consistent with those of our knockdown assay (Figure S3I). In this case, however, it seemed that complete loss of *lncEry* might have impaired the terminal differentiation of erythropoiesis. To investigate how *lncEry* is involved in terminal erythroid differentiation, we analyzed several parameters in the PB of *flox/flox* and Δ/Δ mice (Figure 3C). We found that knocking out *lncEry* reduced the number of RBCs and Retic-Es as well as the concentration of hemoglobin (HGB) (Figure 3D) but had no effect on WBCs in Δ/Δ mice.

Next, we examined the effects of knocking out *lncEry* on HSPCs. The percentage of most HSPC subsets (including LT-HSCs, ST-HSCs, MPP2, MPP3/4, CMPs, MEPs, and MKPs) decreased in Δ/Δ mice compared with their littermate controls (Figures 3E, 3F and Figure S4F). Furthermore, the percentages of preMegE, CFU-E/Pro-E, and PreGM were significantly reduced in the knockout animals (Figure 3F), whereas CLPs and GMPs were minimally affected (Figure 3F and Figure S4G). Loss of *lncEry* apparently impairs the differentiation of erythroid lineage cells.

To examine the function of *lncEry* in erythropoiesis, we compared the colony-forming ability of BM cells from *fIox/fIox* and Δ/Δ mice during *in vitro* culture in complete methylcellulose based medium. We found that the colony numbers of BFU-E, CFU-G, and CFU-GEMM from Δ/Δ mice were lower than those from their littermate controls (Figure 3G and Figure S4H). We then cultured BM or MEP cells in methylcellulose-based medium with erythropoietin (EPO) and established a BFU-E colony assay; the number and size of BFU-E colonies again decreased in Δ/Δ compared to *flox/flox* mice (Figure 3H and Figure S4I). Finally, we cultured MEP cells sorted from *flox/flox* or Δ/Δ mice in methylcellulose-based medium with EPO to support the optimal growth of CFU-E colonies. The colony number decreased for Δ/Δ MEP cells (Figure 3I), indicating that loss of *lncEry* isoform-3 not only affects terminal differentiation during erythropoiesis but also reduces the growth of erythroid progenitor cells.

To directly assess the effect of *lncEry* on the regenerative function of HSCs *in vivo*, we transplanted BM cells from *flox/flox* and Δ/Δ mice (CD45.2) accompanied with competitor cells into irradiated recipients (CD45.1) (Figure 3J). The BM cells from Δ/Δ mice had a lower reconstitution capacity than cells from control mice (71.1% vs. 85.4%) (Figure 3K), but donor cells from the two groups gave rise to the same level of myeloid (Mac-1^+^) and lymphoid (CD3^+^ and B200^+^) lineages (Figure 3L). The frequency of LT-HSCs in Δ/Δ mice was approximately 2-to 3-fold lower than in the control mice (Figure 3E and S4G), and BM cells from Δ/Δ mice showed a 2.3-fold reduction in donor-cell engraftment 16 weeks after transplantation (Figure 3M). These data might explain the decreased level of engraftment observed in recipients when unseparated BM cells were transplanted. Together, these results demonstrated that loss of *lncEry* decreased the growth of MEPs, which ultimately led to suppressed erythroid differentiation and decreased RBC production. Interestingly, although *lncEry* deletion decreased the number of HSCs, the functions of the individual HSCs were unaffected.

We generated an additional *lncEry^fl/fl^* mouse *(flox/flox-2)* model by inserting *loxP* sites around exons 2-6 of *lncEry* isoform-1 using CRISPR/Cas9 technology (Figure S5A). Then, we generated Mx1-Cre; lncEry^fl/fl^ mice (Δ/Δ-2). After poly I:C induction, the knockout efficiency of *LncEry* was confirmed by qPCR (Figure S5B). The deficits of erythroid differentiation and maturation were also observed in the new Δ/Δ-2 mice (Figure S5C-D). These results further confirmed the function of *lncEry* in regulating erythropoiesis.

### *LncEry* deletion decreases *Klf1* expression in MEP cells

To gain mechanistic insights into the function of *lncEry* in erythropoiesis, we isolated MEP cells from *flox/flox* and Δ/Δ mice and performed RNA-seq analysis (Figure 4A). Compared with the *flox/flox* group, 3,256 genes were differentially expressed genes (DEGs), 1,054 of which were downregulated. To explore the changes in chromatin accessibility, we performed transposase-accessible chromatin (ATAC)-seq and observed 7,926 differential peaks (Figure 4B), and 4,599 peaks of accessibility were decreased upon deletion of *lncEry* in MEP cells. Notably, an integrative analysis with RNA-seq and ATAC-seq data revealed a significant correlation between downregulated genes and decreased accessibility, and we identified 421 overlapping genes (Figure S6A). In addition, GO enrichment analysis showed that the overlapping downregulated genes were enriched for erythrocyte differentiation and some metabolism-related terms. We then examined the ROS levels, mitochondrial membrane potential, and glucose uptake of MEP cells (Figure S6C), and the results indicated that the function of *lncEry* in erythropoiesis regulation does not depend on metabolic changes.

**Figure 4.**
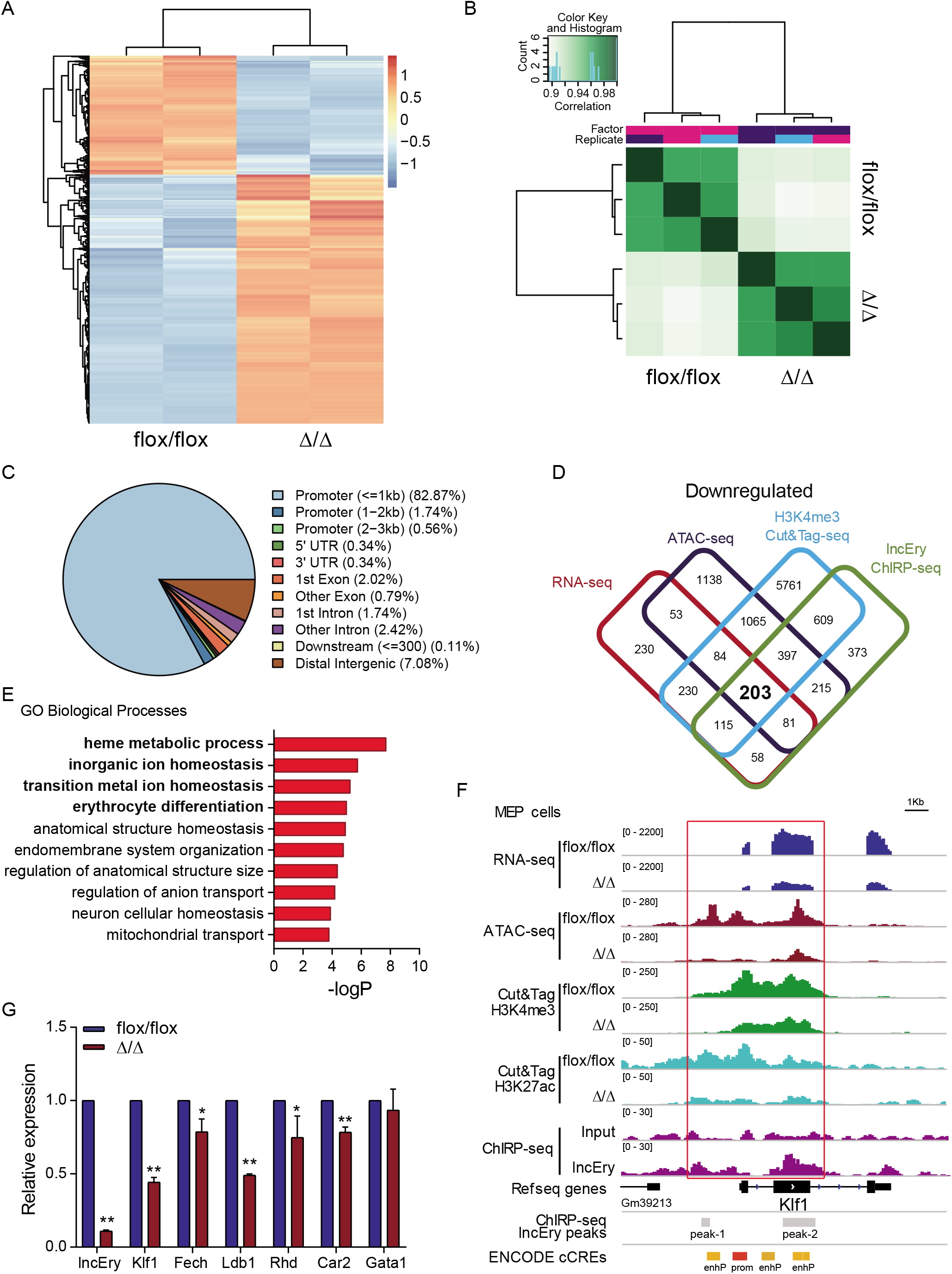
*LncEry* deletion decreases *Klf1* expression in MEP cells. (A) Heat map of differentially expressed genes from RNA-seq of MEP cells sorted from BM of *flox/flox* or Δ/Δ mice. (B) Heat map showing replication of samples from ATAC-seq of MEP cells sorted from BM of *flox/flox* or Δ/Δ mice. (C) Distribution of *lncEry*-binding sites across the indicated intergenic or intragenic regions in MEL cells, as shown by chromatin isolation by ChIRP-seq. (D) Cut&Tag sequence analysis of H3K4me3 in MEP cells sorted from BM of *flox/flox* or Δ/Δ mice; the decreased peak genes under *lncEry* deletion overlapped with the downregulated genes detected by RNA-seq, downregulated peak genes from ATAC-seq and *lncEry*-binding peak genes from ChIRP-seq. (E) GO term analysis of the downregulated genes in (D), −log 10 of the uncorrected P-value on the x axis. (F) Visualization of RNA-seq, ATAC-seq, Cut&Tag sequence of H3K4me3 and H3K27ac and ChIRP-seq peaks and predict cis-regulate elements in *Klf1* regions with IGV software. (G) qPCR analysis of the indicated genes in MEP cells sorted from BM of flox/flox or Δ/Δ mice. *P < 0.05, **P < 0.01.

The nuclear location of a lncRNA might be suggestive of its function in transcriptional regulation^48,54^. To understand the mechanisms of *lncEry* transcriptional regulation, we sought to determine the binding sites for *lncEry* in the genome through chromatin isolation by RNA purification sequencing (ChIRP-seq)^4,48,49^. We performed the ChIRP assay on MEL cells and confirmed the identities of the isolated RNAs by qPCR, and ~17% of *lncEry* RNA was pulled down (Figure S6D). In the sequencing analysis, we identified ~1,786 *lncEry* binding sites when compared with input. When we analyzed the locations of these binding sites in the genome; ~85% were located in promoter regions (Figure 4C) and were mainly concentrated within a region 1 Kbp from the transcriptional start sites (TSS) (Figure S6E). Some of these results support our theory of a role of *lncEry* in transcriptional regulation.

To further explore the transcriptional regulatory function of *lncEry* in MEP cells, we performed cleavage under targets and tagmentation (Cut&Tag) assays^50^ on *lncEry*-deficient MEP cells using an antibody against the histone H3K4me3 (which is associated with transcriptional activation^51^), followed by sequencing (Figure S6F). Then, we performed integrative analysis of the downregulated genes from RNA-seq, ATAC-seq, ChIRP-seq, and H3K4me3 Cut&Tag data to ascertain the directly regulated target genes, and 203 overlapping genes were identified (Figure 4D and Supplementary Table 2). GO enrichment analysis showed that these target genes were enriched in erythrocyte homeostasis- and differentiation-related terms (Figure 4E). In addition, the peaks of H3K4me3 and H3K27ac at the cis-regulate region (CRR) of the *Klf1* gene were decreased upon *lncEry* deletion (Figure 4F, S6G-H). Importantly, *lncEry* could bind to the CRR of *Klf1*, and deletion of *lncEry* decreased *Klf1* expression (Figure 4G and S6I), chromatin accessibility, and the transcriptional active stage of *Klf1* CRR (Figure 4F), suggesting that *lncEry* participates in the transcriptional regulation of *Klf1* in MEP cells. *LncEry* deletion also significantly decreased the expression of other erythrocyte-differentiation-related genes: *Fech, Ldb1, Rhd*, and *Car2* (Figure 4G). In addition, gene set enrichment analysis (GSEA) revealed that deletion of *LncEry* in MEP cells reduced the enrichment of KLF1-target genes (Figure S6H). Together, these results indicate that *lncEry* regulates the transcription of *Klf1* to affect the early stage of erythropoiesis.

### LncEry regulates late-stage erythropoiesis by promoting globin gene expression

To further explore the function of *lncEry* in the late stage of erythropoiesis, we transfected MEL cells with siRNAs targeting *lncEry* and performed RNA-seq analysis. Compared with the control group, 117 and 134 DEGs after *lncEry* knockdown with siRNA-1 and siRNA-2, respectively (Figure 5A). Of these DEGs, 75 overlapping genes were downregulated (Figure 5B and Supplementary Table 3). When we conducted enrichment analysis of these 75 genes with the Metascape online tool, strikingly, the top most enriched term was erythrocyte homeostasis (Figure 5C).

**Figure 5.**
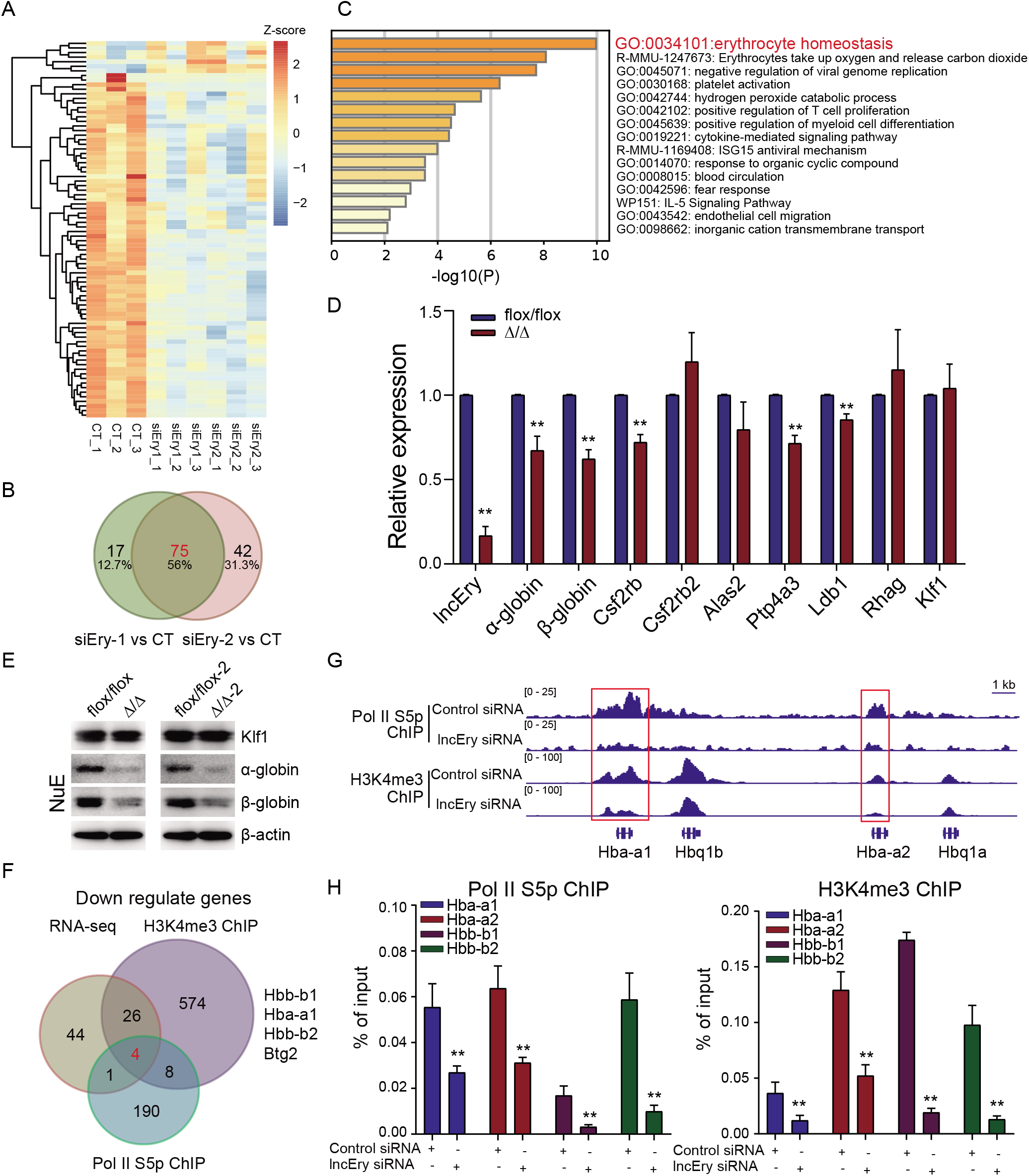
*LncEry* regulates late stage of erythropoiesis by promoting globin gene expression. (A) Heat map of differentially expressed genes in *lncEry*-siRNA-treated and control (CT) MEL cells according to RNA-seq. (B) The numbers of overlapping, downregulated, differentially expressed genes in *lncEry*-depleted and CT MEL cells using two different targeting siRNAs. (C) Gene enrichment analysis of downregulated genes with −log 10 plot of the uncorrected P-value on the x-axis; darker shading corresponds to a greater level of enriched genes in each term. (D) qPCR of indicated genes in NuE cells sorted from BM of *flox/flox* or Δ/Δ mice. (E) Cellular extracts of NuE cells sorted from BM of *flox/flox*, Δ/Δ, *flox/flox-2*, or Δ/Δ-2 mice were prepared and analyzed by western blotting. (F) MEL cells were transfected with control siRNA or *lncEry* siRNA. Soluble chromatin was collected for ChIP-seq analysis using antibodies against Pol II–S5p or H3K4me3; the decreased peak genes under *lncEry* knockdown overlapped with the downregulated genes detected by RNA-seq. (G) ChIP-seq trace showing Pol II–S5p and H3K4me3 binding of control or lncEry knockdown cells in relation to the indicated gene CRRs, visualized with IGV software. (H) qChIP of Pol II–S5p (left panel) or H3K4me3 (right panel) with primers covering the promoters of the indicated genes. Each bar represents the mean ± S.D. for triplicate experiments. *P < 0.05, **P < 0.01.

We then performed qPCR to analyze the expression of several DEGs: *lncEry* knockdown significantly decreased the expression of erythrocyte-homeostasis- and differentiation-related genes, such as *Hba-a1, Hba-a2* (downregulated in siEry2 parts) (two variants of α-globin), *Hbb-b1, Hbb-b2* (two variants of β-globin), *Alas2*, and *Rhag* in MEL cells (Figure S7A). Furthermore, *lncEry* knockdown significantly reduced the protein levels of α- and β-globin (Figure S7B). We also examined DEGs in NuEs sorted from the BM of *flox/flox* and Δ/Δ mice, and *lncEry* knockout decreased mRNA expression and protein levels of α- and β-globin (Figure 5D-E). Unlike in MEPs, *lncEry* knockout did not reduce the expression of *Klf1* in NuE cells (Figure 5D-E), and the DEGs in MEL cells were also not enriched in *Klf1*-target genes (Figure S7C). These results indicate that *lncEry* participates in erythropoiesis regulation at different stages and using different mechanisms.

To further explore the role of *lncEry* in regulating DEGs at the transcriptional level, we performed chromatin immunoprecipitation (ChIP) assays in *lncEry*-deficient cells using antibodies against Ser5-phosphorylated RNA polymerase II (Pol II–S5p) and histone H3K4me3, followed by sequencing. We then compared the peaks enriched by Pol II–S5p and histone H3K4me3 of the gene promoters in control and *lncEry*-depleted cells. Interestingly, we found four downregulated DEGs in our RNA-seq dataset (*Hbb-b1, Hba-a1, Hbb-b2*, and *Btg2*) that overlapped in the Pol II–S5p and H3K4me3 ChIP-sequencing datasets (Figure 5F and Figure S7D). Consistently, the Pol II–S5p and H3K4me3 enrichment peaks of globin gene cis-regulated regions, such as *Hba-a1*, *Hba-a2, Hbb-b1*, and *Hbb-b2*, sharply declined upon *lncEry* knockdown (Figure 5G and Figure S7E), and the results of the qChIP assays verified these findings (Figure 5H). We thus concluded that *lncEry* depletion affects the transcription of globin genes in the later stage of erythropoiesis.

### *LncEry* is physically associated with Wdr82

*LncRNAs* are usually associated with numerous cellular functions, most of which require interactions with one or more RNA-binding proteins^52,53^. To determine whether *lncEry* acts alone or in concert with other proteins in the different stages of erythropoiesis, we performed RNA-pulldown assays (Figure S8A) followed by silver staining and mass spectrometry to identify *lncEry* isoform-3 interaction partners in MEPs sorted from the BM of wildtype mice and MEL cells. Then, we performed integrative analysis and found 11 common interacting partners in the two types of cells (Figure 6A-B, S8B, and Supplementary Tables 4 and 5). Interestingly, we identified two transcription regulators, WD repeat-containing protein 82 (WDR82) and DEAD-box helicase 5 (DDX5), as likely *lncEry* binding partners, which may explain some of the mechanisms involved in regulating the DEGs of *lncEry*-depleted MEP and NuE cells at the transcriptional level. Western blot analyses confirmed the interactions between *lncEry* and its binding partners (Figure 6C).

**Figure 6.**
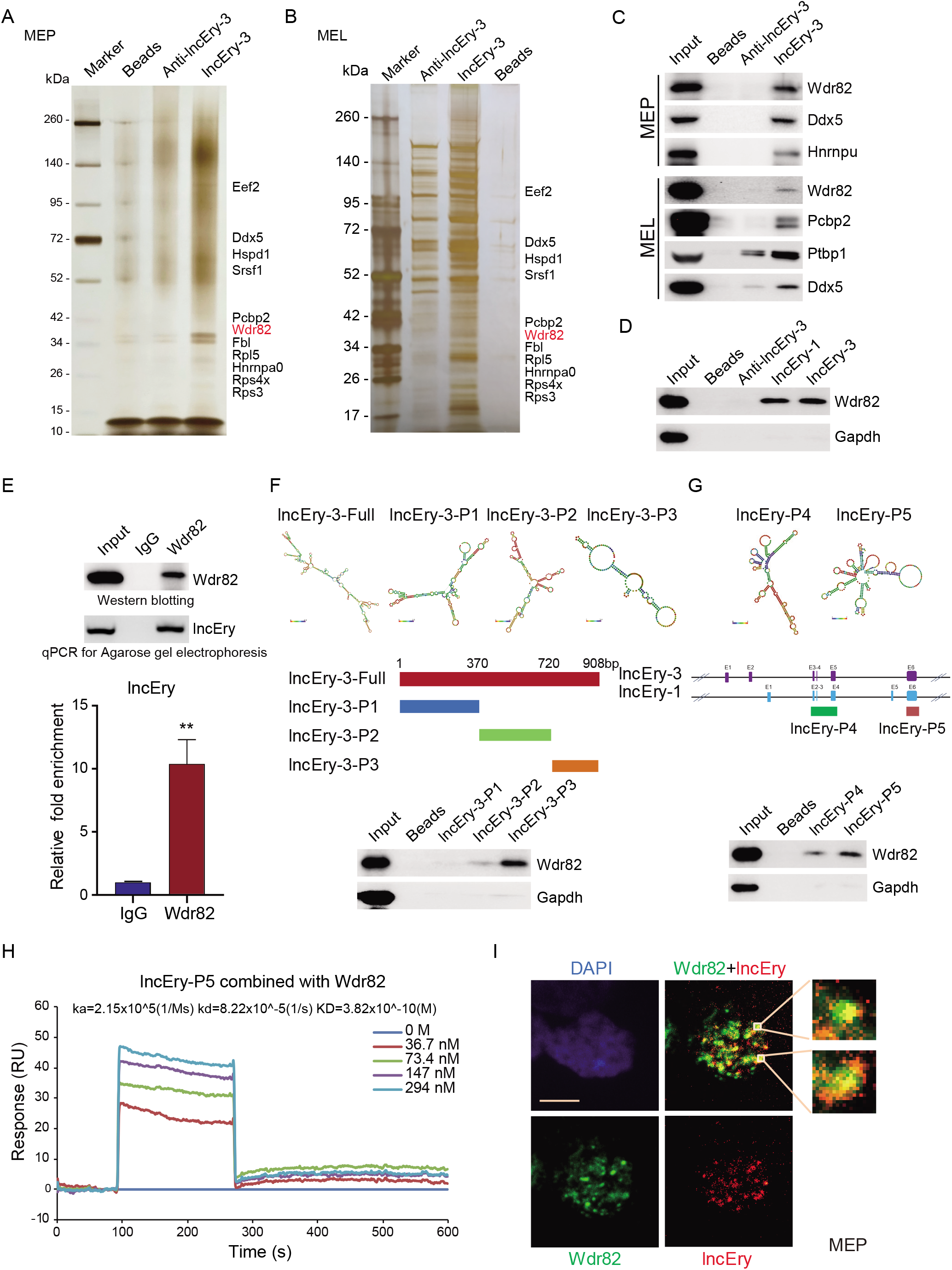
*LncEry* is physically associated with Wdr82. (A-B) RNA pulldown and mass spectrometry analyses of *lncEry*-interacting proteins. Whole-cell extracts from MEP (A) or MEL (B) were prepared and subjected to RNA pulldown using *in vitro* transcribed *lncEry*-3 or anti-*lncEry*-3 as the bait. After extensive washing, the bound proteins were eluted and visualized by silver staining on SDS-PAGE. The protein bands on the gel were recovered by trypsinization and analyzed by mass spectrometry. Detailed results from the mass spectrometric analysis are provided in Supplementary Tables 4 and 5. (C-D) RNA-pulldown-purified proteins retrieved using the indicated baits were analyzed by western blotting with antibodies against the indicated proteins. (E) Whole-cell lysates from MEL cells were immunoprecipitated with Wdr82 antibodies; purified RNA was analyzed by qPCR and agarose gel electrophoresis. (F) Schematic of minimum free energy (MFE) structure and the *lncEry* isoform-3 truncation mutant. RNA-pulldown-purified proteins retrieved by *lncEry* truncation mutant baits were analyzed by western blotting. (G) Schematic of the MFE structure and the lncEry truncation mutants. RNA pull-down purified proteins by lncEry truncation mutants baits were analyzed by western blotting with antibodies against the indicated proteins. (H) Sensorgrams of *lncEry-P5* binding to Wdr82, as measured by SPR technology on a Biacore 3000 instrument. Representative sensorgrams were obtained by injecting various concentrations (0, 36.7, 73.4, 147, and 294 nM) of *lncEry-P5* over Wdr82 immobilized on a CM5 sensor chip. (I) RNAscope and immunofluorescence assays using *lncEry* probes and Wdr82 antibodies, respectively, of MEP cells and analysis by confocal microscopy. Scale bar, 10 μm. Each bar represents the mean ± S.D. for triplicate experiments. **P < 0.01.

To explore the functional impact of Wdr82 and Ddx5 on HSPCs, we transduced cKit^+^ cells with a lentivirus carrying GFP-fused *Wdr82* or *Ddx5* shRNAs and performed *in vitro* colony assays (Figure S8C and S8D). The results showed that *Wdr82* knockdown decreased the numbers of CFU-G, CFU-M, and BFU-E, whereas Ddx5 knockdown did not decrease BFU-E numbers (Figure S8C and S8D). The knockdown efficiency was confirmed by western blotting (Figure S8E). Then, we cultured *Wdr82* knockdown cKit^+^ cells in M3436 methylcellulose-based medium with EPO and established a BFU-E colony assay. The colony number and size of the BFU-E colonies decreased in *Wdr82*-knockdown cells (Figure S8F). However, when we cultured the cKit^+^ cells *in vitro* for about 1 week after they were transduced with *Wdr82* shRNA, *Wdr82* knockdown was seen to decrease the percentage of late-stage erythropoiesis cells (Ter119^+^CD44^-^) and arrest the progress of erythropoiesis (Figure S8G). These results suggest that Wdr82, with similar phenotypes to *lncEry*, was more likely to participate in erythropoiesis with *lncEry* than Ddx5. Following this, we were interested in exploring the function of *lncEry* interaction with Wdr82 in regulating the early and late stages of erythropoiesis. When we examined the interaction between *lncEry* isoform-1 or isoform-3 and Wdr82, we found no evidence of transcriptional specificity (Figure 6D). Consistent with these findings, RNA immunoprecipitation (RIP) assays further confirmed this interaction between all *lncEry* isoforms and Wdr82 (Figure 6E).

We were then intrigued to identify the binding sites underlying the *lncEry–Wdr82* interaction. To do so, we generated *lncEry* isoform-3 truncation mutants according to the isoform’s main minimum free energy (MFE) stem-loop regions, which we predicted using RNAfold WebServer tools (http://rna.tbi.univie.ac.at) (Figure 6F, upper panel). *In vitro* RNA-pulldown assays showed that the interaction was primarily dependent on the *lncEry* C-terminal loop regions (Figure 6F). We also generated two *lncEry* fragments that are shared by all three isoforms: exons 2-4 and 6 of *lncEry* isoform-1. The results of the binding assay revealed that *lncEry* mainly interacted with Wdr82 through the last exon transcript (exon 6) (Figure 6G). Next, we conducted surface plasmon resonance assays using the GE Healthcare Biacore 3000 platform to examine the binding kinetics of *lncEry* and Wdr82. Indeed, the last exon region *lncEry-P5* (exon 6) shared by each of the three lncEry isoforms directly interacted with Wdr82 that was purified from MEL cell lines (Figure S8H) with a KD value of 38.2 nM (Figure 6H). Finally, we performed co-localization assays combining RNAscope with immunofluorescent staining followed by fluorescent confocal microscopy. We observed that *lncEry* mainly co-localized with Wdr82 in the nuclei of MEP and MEL cells (Figure 6I and Figure S8I). All *lncEry* isoforms physically associated with Wdr82 in the nuclei of MEP and MEL cells. We thus proposed a hypothesis that the lncEry–Wdr82 complex serves to regulate transcription in these cells.

### *LncEry*–Wdr82 regulates the transcriptional activation of *Klf1* in MEP cells

As *lncEry* was found to be associated with Wdr82 (Figure 6), we further explored the molecular mechanisms of the role of *lncEry* and Wdr82 in MEP cells by performing Cut&Tag assays on *lncEry*-deficient MEP cells using an antibody against Wdr82. The results showed that *lncEry* deletion decreased chromatin accessibility and the binding of Wdr82 at CRR around *Klf1* gene body as well as the binding of Wdr82 at whole genome region (Figure 7A and S9A). Therefore, we speculated that *lncEry* can physically interact and functionally coordinate with Wdr82 to regulate the transcription of *Klf1*. To test this, we established two pGL3-luciferase reporters containing CRRs, as shown in Figure 7A, then performed reporter assays with pGL3-luciferase reporters containing *Klf1* CRRs or mutant CRRs (without the main *LncEry* binding site) co-transfected into 293T cells together with *lncEry*, Wdr82, or both, as well as the *Renila* luciferase vector for normalization (Figure 7B and S9B). The assays showed that overexpression of *lncEry* or co-expression of *lncEry* and Wdr82 enhanced the activity of the *Klf1-CRR* reporter, but not the *Klf1*-CRRs-mt reporter (Figure 7B). These results suggest that *lncEry* coordinates with Wdr82 to regulate the transcription of *Klf1* in early erythroid differentiation.

**Figure 7.**
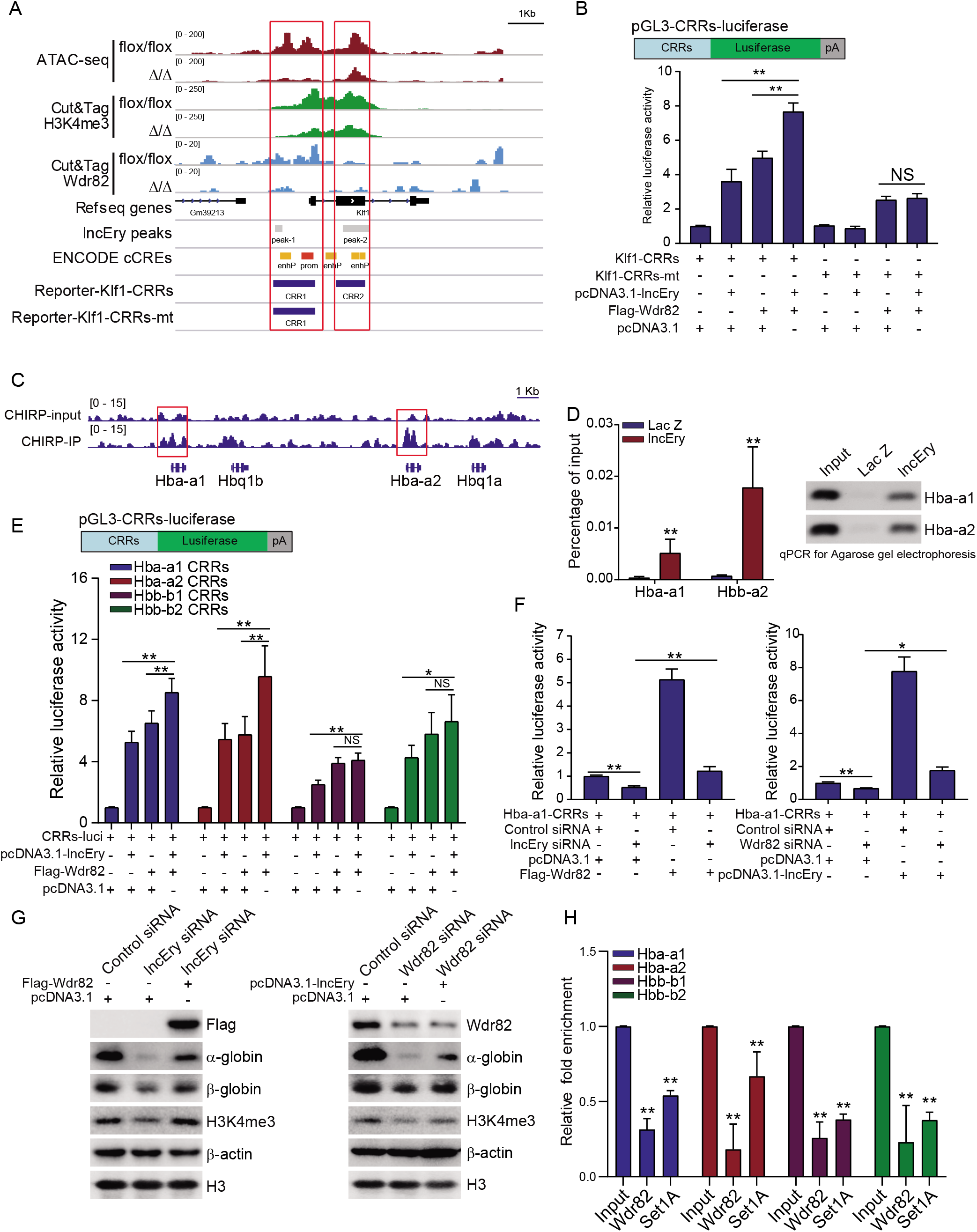
*LncEry-Wdr82* regulates transcriptional activation of Klf1 and globin genes. (A) Visualization of ATAC-seq, Cut&Tag for H3K4me3 and Wdr82, ChIRP-seq *lncEry* peaks, predict cis-regulate elements, and luciferase reporter clone CRRs in *Klf1* gene regions with IGV software. (B) Schematic diagrams of the pGL3-CRRs-luciferase reporter constructs and relative luciferase activity was determined by sequential normalization to Renilla and pGL3-vector activity. (C) ChIRP-seq trace showing *lncEry* binding in relation to the indicated gene regions. Red boxes indicate the CRRs of globin genes (D) qPCR and agarose gel electrophoresis analysis of isolated chromatin sequences of indicated gene promoter regions. (E) Schematic diagrams of pGL3-CRRs-luciferase reporter constructs (upper panel). For reporter assays, MEL cells were co-transfected with pcDNA3.1-lncEry or Wdr82 or pcDNA3.1-lncEry and Wdr82 together with Renilla and globin gene CRRs luciferase. The relative luciferase activity was determined by sequential normalization to Renilla and pGL3-vector activity. (F) MEL cells were co-transfected with the indicated siRNAs or plasmids together with Renilla and the globin gene CRRs luciferase. The relative luciferase activity was determined by sequential normalization to Renilla and pGL3-vector activity. (G) MEL cells were co-transfected with the indicated siRNAs or plasmids and the cellular extracts were analyzed by western blotting with antibodies against the indicated proteins. (H) MEL cells were transfected with control or *lncEry* siRNAs; the soluble chromatin was immunoprecipitated with antibodies against Wdr82 or Set1A and analyzed by qPCR with the indicated primers. The relative fold enrichment was determined by sequential normalization with the cycle threshold values of the input and control siRNA samples. Each bar represents the mean ± S.D. for biological triplicate experiments. *P < 0.05, **P < 0.01.

### *LncEry*–Wdr82 regulates the transcriptional activation of globin genes

In the late stage of erythropoiesis, *lncEry* depletion decreased the transcription of globin genes. We found that *lncEry* was located at the CRRs of *Hba-a1* and *Hba-a2* (Figure 7C) but not the genomic regions of *Hbb-b1* and *Hbb-b2* (Figure S9C). Furthermore, using qPCR, we detected *lncEry* binding on the CRRs of globin genes (Figure 7D and Figure S9C). However, we were unable to detect *lncEry* binding on *Hbb-b1* and *Hbb-b2* CRRs, possibly because it has lower affinity for these promoters or because cofactors are required for *lncEry* binding.

Because *lncEry* associated with Wdr82 and directly bound to the CRRs of globin genes in MEL cells (Figure 6C and 6D), we speculated that *lncEry* physically interacts and functionally coordinates with Wdr82 to regulate globin gene transcription in late erythropoiesis. To test this, MEL cells were co-transfected pGL3-luciferase reporters containing *Hba-a1, Hba-a2, Hbb-b1*, or *Hbb-b2* promoters (Figure 7E) along with *lncEry*, Wdr82, or both, or the *Renila* luciferase vector. Reporter assays showed that overexpression of either *lncEry* or *Wdr82* resulted in a significant increase in *Hba-a1, Hba-a2, Hbb-b1*, and *Hbb-b2* reporter activity. Co-expression of *lncEry* and *Wdr82* enhanced *Hba-a1* and *Hba-a2* (but not *Hbb-b1* or *Hbb-b2*) reporter activity further, which was perhaps because reporter activity was saturated (Figure 7E). Flag-Wdr82 expression was confirmed by western blotting (Figure S9D).

To investigate whether the effects of *lncEry* on the transcriptional activation of globin genes were associated with Wdr82, we transfected Flag-*Wdr82* plasmids into *lncEry*-deficient MEL cells. Subsequent luciferase reporter assays showed that *lncEry* depletion inhibited *Hba-a1, Hba-a2, Hbb-b1*, and *Hbb-b2* reporter activities, but this inhibition was rescued, at least in part, upon Flag-*Wdr82* overexpression (Figure 7F and Figure S9E-H). Similarly, Wdr82 depletion inhibited the four globin gene reporter activities, and this inhibition was rescued by *lncEry* overexpression (Figure 7F and Figure S9E-H).

We confirmed the effects of *lncEry* and Wdr82 on globin gene transcription at the protein level, and again, *lncEry* depletion decreased the protein expression of α-globin and β-globin, and Wdr82 overexpression partially rescued the effect (Figure 7G, left panel) and *vice versa* (Figure 7G, right panel). Interestingly, *lncEry* depletion decreased the level of H3K4me3, which could be rescued by overexpression of Wdr82, and *vice versa* (Figure 7G and Figure S9I). These results led us to propose that *lncEry* might participate in the Set1A/Wdr82 complex to affect H3K4me3 levels and, as a result, regulate the transcriptional activation of globin genes.

To strengthen the above hypothesis, our final analyses examined the effects of *lncEry* on Wdr82 and Set1A recruitment to globin gene CRRs using qChIP assay. To this end, we immunoprecipitated soluble chromatin from control or *lncEry*-depleted MEL cells using antibodies against Wdr82 or Set1A, and then performed qPCR analysis to identify the precipitated DNA. *lncEry* depletion specifically decreased Wdr82 and Set1A enrichment on the CRRs of globin genes, but not the two other components of the Set1A/Wdr82 complex Ash2l and RbBP5^54^ (Figure 7H and S9J). These findings might be due to the low affinity of Ash2l and RbBp5 antibodies for ChIP samples (Figure 7H and Figure S9J). Collectively, these results support the hypothesis that *lncEry* recruits Wdr82 and stabilizes the Set1A/Wdr82 complex located on the CRRs of globin genes (Figure 8).

**Figure 8.**
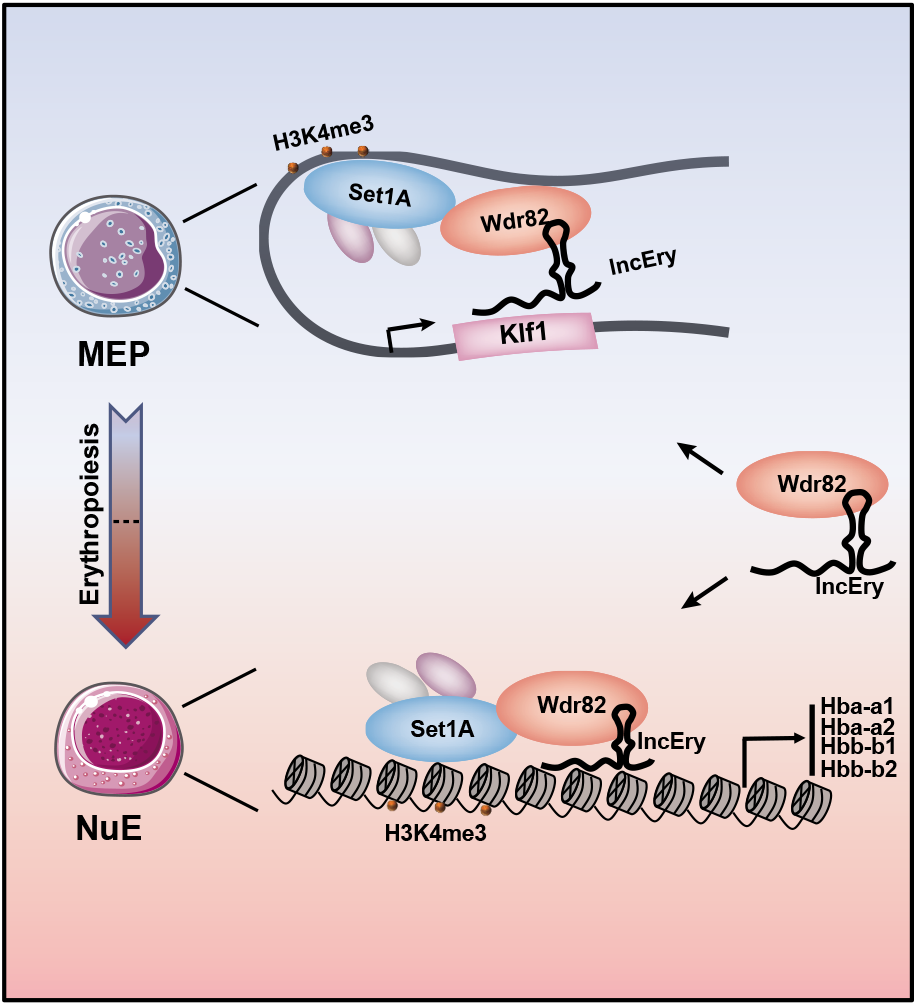
Mechanistic model of how *lncEry* combines with Wdr82 to participate in erythropoiesis regulation. Generally, the novel annotated lncRNA, *lncEry*, interacts with Wdr82 to control the location of the Set1A/Wdr82 complex and facilitate H3K4me3 binding to CRRs of target genes. In MEP cells, *lncEry* bind to the CRRs of *Klf1* to recruit Wdr82 and elevate H3K4me3 levels at CRRs to promote *Klf1* expression and coordinate early erythroid differentiation. In NuE, *lncEry* binds to CRRs of globin genes and combines with Wdr82 to promote transcription of *Hba-a1, Hba-a2, Hbb-b1*, and *Hbb-b2*, which promotes terminal erythropoiesis. In brief, *lncEry* combines with Wdr82 to promote *Klf1* and globin gene expression to regulate the early and late stages of erythropoiesis, respectively.

## DISCUSSION

Because a number of lncRNAs have been shown to be important for hematopoiesis, in this study, we aimed to investigate the importance of these regulatory factors in erythropoiesis, about which much less is known. We found a novel functional lncRNA, *Gm15915 (lncEry)*, that is specifically and highly expressed in MEPs and NuEs, and we annotated a new and highly expressed *lncEry* isoform localized in the nucleus. *LncEry* depletion decreased *Klf1* and globin gene expression in erythroid progenitors and mature cells, respectively, and therefore impaired murine erythroid cell differentiation and maturation. Mechanistically, *lncEry* together with Wdr82 facilitates the binding of the histone H3K4me3 to the CRRs of *Klf1* and globin genes, which in turn, regulates the early and late stages of erythropoiesis (Figure 8).

Erythroid differentiation is regulated at multiple levels to ensure the proper generation of mature cells under multiple physiological conditions^27^. Numerous functional lncRNAs that regulate cellular processes, such as cell development, differentiation, division, survival, and death, have been identified in recent years^55,56^. The nuclear localization of *lncEry* suggests that it might participate in regulating gene expression by modulating certain nuclear events such as epigenetic modifications, transcription, or mRNA splicing^57^. Wdr82 is a unique subunit of the Set1A (KMT2F) histone H3-Lys4 methyltransferase complex and can recruit the Set1A complex to the transcription start sites of target genes and bind to Ser 5 phosphorylated RNA polymerase II to promote transcriptional activation^58–60^. Recruitment of the KMT2 complex to specific chromatin regions by lncRNAs has been reported^61^, which supports our hypothesis that *lncEry* interacts with Wdr82 and participates in transcriptional activation at CRRs. Furthermore, Wdr82 depletion results in dysfunction of the Set1A/Set1B complex, affecting H3K4me3 status and inhibiting the transcriptional activation of Pou5f1, thereby preventing early embryonic development^62^. Whether *lncEry* also participates in embryonic development with Wdr82 needs further exploration.

For transcriptional regulation, recruitment of the mixed lineage leukemia (MLL) complex to specific chromatin regions not only depends on the presence of plant homeodomains but also lncRNAs; for example a long intergenic non-coding RNA, HOTTIP, guides WDR5 to chromatin to recruit the MLL complex^63,64^. As a member of the Set1A complex, Wdr82 has affinity for the *lncEry* C terminus, which is a common region of all *lncEry* isoforms. *lncEry* can generally target CRRs of *Klf1* and globin genes, thus we speculated that binding of the Wdr82/Set1A complex to the genome is likely to partially depend on lncEry. According to previous reports, Wdr82 promotes Set1A-dependent H3K4 trimethylation^54^ to participate in transcriptional activation. Even so, lncEry depletion decreased enrichment of Wdr82 and the level of H3K4me3 in the CRRs of *Klf1* and globin genes, which supports our hypothesis that *lncEry* directs the Set1A/Wdr82 complex to the CRRs of *Klf1* and globin genes in MEP and MEL cells and stabilizes its location. We thus believe we have identified a new and important regulator associated with the Wdr82/Set1A complex that promotes the transcriptional activation of *Klf1* and globin genes in early- and late-stage erythropoiesis, respectively.

Most lncRNAs show little conservation between species and exhibit a rapid evolutionary turnover. Indeed, we did not find *lncEry* in human cells, yet there are two annotated lncRNAs (ENST00000552603.1 and ENST0000055074.1) similar to *lncEry* located anti-sense of *Ntn4*. However, we did not manage to show these lncRNAs to have the same function or expression pattern as *lncEry* (data not shown). Despite this, we cannot exclude the possibility that other functional lncRNAs bind to the Wdr82/Set1A complex to participate in related biological processes.

In conclusion, we have identified a potential component of the Wdr82/Set1A complex that participates in the transcriptional regulation of *Klf1* and globin genes and, by extension, the early and late stages of erythroid differentiation to regulate erythropoiesis. Whether lncRNAs together with Wdr82 are viable targets for manipulating other processes and their clinical application need to be addressed in additional research.

## METHODS

### Mice

C57BL/6J and B6.SJL mice were purchased from the animal facility of the State Key Laboratory of Experimental Hematology (SKLEH, Tianjin, China). *lncEry^flox/+^* mice were generated by Beijing Biocytogen Co.,Ltd. All animal procedures were performed in compliance with the animal care guidelines approved by the Institutional Animal Care and Use Committees of the SKLEH and the Institute of Hematology.

### Antibodies and reagents

The following antibodies were used in this study: FLAG (F1804, 1:2000 for Western blot), β-actin (A1978, 1:10,000 for WB) from Sigma; histone H3 (ab1791, 1:10,000 for WB), RNA polymerase II (phospho S5) (ab5408, 1:200 for ChIP), H3K4me3 (ab8580, 1:200 for ChIP and Cut&Tag), and H3K27ac (ab6002, 1:200 for Cut&Tag) from Abcam; Wdr82 (99715, 1:1000 for WB, 1:100 for Cut&Tag), DDX5 (9877T, 1:1,000 for WB), and RBBP5 (13171S, 1:200 for ChIP) from Cell Signaling Technology; PCBP2 (NBP2-19715, 1:1,000 for WB) from Novusbio; PTBP1 (32-4800, 1:1,000 for WB) from Invitrogen; α-globin (14537-1-AP, 1:2,000 for WB) from Proteintech; EKLF (OM184222, 1:1,000 for WB) and β-globin (OM256195, 1:1,000 for WB) from OmnimAbs; ASH2L (A11278, 1:200 for ChIP) from Abclonal. Anti-FLAG M2 affinity gel (A2220), 3 × FLAG peptide (F4799), doxycycline (D9891), and 2’,7’-dichlorofluorescin diacetate (DCF-DA) (35845) were purchased from Sigma. Guinea pig anti-rabbit IgG (Heavy & Light Chain) antibody (ABIN101961, 1:100 for Cut&Tag) was from Antibodies Online. Tetramethylrhodamine methyl ester (TMRE) (I34361) and 2-NDBG [2-(N-(7-Nitrobenz-2-oxa-1,3-diazol-4-yl)Amino)-2-Deoxyglucose] (N13195) were purchased from Invitrogen. Recombinant murine SCF (250-03), murine IL-3 (213-13) and Recombinant Human EPO (100-64) were obtained from PeproTech.

### ROS levels, mitochondrial membrane potential and glucose uptake analysis

MEP cells from flox/flox or Δ/Δ mice were labeled with surface markers, and after 30 min of incubation at 4°C, the cells were washed with 1 mL staining buffer. Then, the cells were stained with DCFH-DA, TMRE or 2-NDBG for 20 minutes at 37°C with shaking, and washed with 2 mL cold staining buffer. Finally, the cells were immediately analyzed by flow cytometry.

### Plasmids and viral production

The FLAG-tagged *lncEry* isoforms or truncation mutants were expressed using a pcDNA3.1 vector. FLAG-tagged *Wdr82* was inserted into a pLenti-Tight-Puro vector. *Klf1, Hba-a1, Hba-a2, Hbb-b1*, and *Hbb-b2* CRRs and luciferase were ligated into pGL3-luciferase vectors. The *lncEry, Wdr82*, and *Ddx5* LV-shRNA-GFP lentiviruses were produced by GeneChem (Shanghai, China). For lentiviral production, the target plasmid was transfected together with *pSPAX2* and *pMD2G* into 293T cell lines using Lipofectamine 2000. The supernatant was harvested after 48 and 72 h of culture and concentrated using an Amicon filter (100K NMWL, Millipore).

### Cell culture

MEL and HEK293T cells were purchased from the American Type Culture Collection (Manassas, VA) and cultured in RPMI1640 (A10491-01, Gibco) or DMEM (SH30243.01, Hyclone) medium with 10% fetal bovine serum (FBS) respectively. BM cells were obtained from C57BL/6 mice and cultured in SFEM (09650) medium (STEM CELL). Cells that allow protein expression under doxycycline treatment were created using two steps. First, cells were infected with a lentivirus carrying rtTA and selected using neomycin. The established rtTA cells were subsequently infected with a virus expressing pLenti-Tight-Puro vector that encodes Wdr82 and selected using puromycin. All of the cells integrated with rtTA were cultured in Tet-Approved FBS and RPMI1640. All cells were authenticated by examination of their morphology and growth characteristics and were confirmed to be mycoplasma-free.

### Western blotting

Western blotting was performed as described previously^65^. In brief, cellular extracts were harvested from cells and resuspended in 5 × SDS-PAGE loading buffer. The boiled protein samples were then subjected to SDS-PAGE followed by immunoblotting with the appropriate primary antibodies and secondary antibodies.

### RACE assays

5’ and 3’ RACE reactions were performed to isolate full-length *lncEry* from the total RNA of MEP cells using the 5’- and 3’-Full RACE Kits (TaKaRa, Dalian, China) in accordance with the manufacturer’s instructions. Primers used for RACE are listed in Supplementary Table 6.

### Flow cytometry

For cell-sorting experiments using mouse MEP cells, cKit^+^ cells were enriched before flow cytometry using cKit magnetic beads (Miltenyi Biotec). The cells were subsequently stained with a lineage cocktail and cKit (eBioscience, 17-1171-82, 1:200), Sca-1 (eBioscience, 25-5981-82, 1:200), CD34 (eBioscience, 13-0341-82, 1:100), and CD16/CD32 (eBioscience, 45-0161-82, 1:200) antibodies. The lineage cocktail included Gr-1 (Biolegend, 108424, 1:400), Mac-1 (Biolegend, 101226, 1:400), B220 (Biolegend, 103224, 1:400), CD4 (Biolegend, 100414, 1:400), CD8 (Biolegend, 100714, 1:400), CD3 (Biolegend, 100330, 1:400), and Ter-119 (Biolegend, 116223, 1:400). DAPI (D9542, 1 mg/mL, Sigma-Aldrich) was used to exclude dead cells. For the HSPC analysis, nucleated BM cells were stained with lineage-specific antibodies against Sca-1, cKit, CD34, CD16/CD32, and CD127 (Biolegend, 135040, 1:400). For PreMegE and MPP analyses, nucleated BM cells were stained with lineage cocktail, Sca-1, cKit, CD16/CD32, CD41 (eBioscience, 46-0411-82, 1:400), CD150 (Biolegend, 115904, 1:400), CD48 (Biolegend, 103432, 1:400), and CD105 (Biolegend, 120409, 1:400). The lineage cocktail included CD3, CD4, CD8, B220, Gr-1, Ter119, and Mac-1. For Pro-E analysis, nucleated BM cells were stained with CD3, B220, Mac-1, Ter119, and CD44 (eBioscience, 45-0441-82, 1:200). A modified LSR II flow cytometer with four lasers (355 nm, 488 nm, 561 nm, and 633 nm) was used for the analysis, and an Aria III flow cytometer with four lasers (375, 488, 561, and 633 nm) was used for sorting. The analyses were performed using FACSDiVa and FlowJo (Tree Star) software.

### Colony-forming assays

Murine BM cells were cultured in M3434 complete methylcellulose-based medium (03434, StemCell Technologies) for 10-14 days to generate BFU-E, CFU-G, CFU-M, CFU-GM, and CFU-GEMM colonies. Murine BM or sorted MEP cells were cultured in M3436 medium (03436, StemCell Technologies) for 48 h to generate CFU-E colonies. Murine-sorted MEP cells were cultured in M3334 medium (03334, StemCell Technologies) for 10-14 days to generate BFU-E colonies.

### Transplantation

For lncEry knockdown assay, GFP-fused control or *lncEry* shRNA lentiviruses were transduced into B6.SJL (CD45.1^+^) mouse cKit^+^ HSPCs, and 4 x 10^5^ transduced cells (CD45.1^+^GFP^+^) were transplanted together with 1.5 x 10^5^ CD45.2^+^ BM cells into lethally irradiated C57BL/6J (CD45.2) recipients and repopulation was measured monthly. For competitive BM transplantation, 5 x10^5^ BM cells of *flox/flox* or Δ/Δ mice, were transplanted into lethally irradiated (9.5 Gy) B6.SJL recipient mice in competition with 2.5 x 10^5^ CD45.1^+^ BM cells and repopulation was measured.

### RNA pulldown and mass spectrometry analyses

Substrate RNAs were synthesized by *in vitro* transcription using a T7 RNA production system (Promega, P1300) in accordance with the manufacturer’s instructions. The 3’-end desthiobiotin-labeled RNA probes used in the RNA pulldown were generated using an RNA 3’ End Desthiobiotinylation Kit (20163, Thermo Scientific) following the manufacturer’s instructions. RNA pulldown was performed using a Magnetic RNA-Protein Pull-Down Kit (20164, Thermo Scientific). In brief, 3’-end desthiobiotin-labeled RNA probes were captured by streptavidin magnetic beads and mixed with MEP or MEL cell extracts (containing 1 mg of protein) in IP lysis buffer (87787, Thermo Scientific) in protein-RNA-binding buffer and incubated for 30-60 min at 4°C with agitation or rotation. After general washing, the retrieved proteins were detected by western blotting or subjected to silver staining and LC-MS/MS analysis (Supplementary Tables 4 and 5).

### Nano-HPLC-MS/MS analysis of *lncEry*-binding proteins

To identify proteins associated with *lncEry*, LC-MS/MS analysis was performed on an Orbitrap Q Exactive mass spectrometer. Tryptic peptides were separated by reverse-phase liquid chromatography on an easy-nLC 1000 system (Thermo Fisher Scientific) and directly sprayed into a Q Exactive Plus mass spectrometer (Thermo Fisher Scientific). Mass spectrometry analysis was carried out in data-dependent mode with an automatic switch between a full MS and an MS/MS scan on the Orbitrap. For the full MS scan, the automatic gain control target was 1e6 and the scan ranged from 300 to 1800 with a resolution of 70,000. The 10 most intense peaks with a charge state ≥2 were selected for fragmentation by high-energy collision dissociation with a normalized collision energy of 27%. The MS2 spectra were acquired with 17,500 resolutions. All MS/MS spectra were searched against the Uniport-Human protein sequence database using Mascot 2.2. Peptide sequences were searched using trypsin specificity while allowing for a maximum of two missed cleavages. Cys carbamidomethylation was specified as a fixed modification, and oxidation of methionine was fixed as a variable modification. The peptide mass tolerance was set at ±20 ppm, and the fragment mass tolerance was set at ±0.1 Da.

### RNA immunoprecipitation (RIP)

RIP assays were performed using a Magna Nuclear RIP Kit from Merck Millipore (17-10520, Millipore) with 5 μg IgG or anti-Wdr82 antibody in accordance with the manufacturer’s instructions. The eluted RNAs were purified using TRIzol LS regent (Life Technology). We then performed qPCR analysis with the indicated primers (Supplementary Table 6).

### Quantitative PCR (qPCR)

Total cellular RNA was isolated with TRIzol reagent (Invitrogen) and used for first-strand cDNA synthesis via the Reverse Transcription System (Roche). Quantitation of all gene transcripts was made by qPCR using a Power SYBR Green PCR Master Mix (Roche) and Thermo Quant Studio 5 sequence detection system (Thermo) with the expression of ACTB as the internal control. The primers used are listed in Supplementary Table 6.

### RNAscope and Immunofluorescence assays

50,000 cells were centrifuged onto slides on 800 g for 5 min. Cells were fixed with 4% paraformaldehyde for 15 min and dehydrated in 50%, 70%, and 100% ethanol for 2 min each. After incubating with hydrogen peroxide and proteinase IV, the samples were treated using the RNAscope multiplex fluorescent V2 assay kit (323100, ACD) and incubated with the specific probes (Mm-Gm15915-O2: 555551; Negative Control DapB: 310043 and Positive Control Mm-Ppib: 313911 from ACD) and Cy3 dye (designed by ACD) in accordance with the manufacturer’s instructions. Before staining with DAPI, the samples were incubated with the appropriate primary and secondary antibodies coupled to AlexaFluor 488 (Invitrogen) following the instructions provided by ACD. Confocal images were captured using a Spinning Disk Confocal Microscope system (UltraVIEW VOX) with a ×100 oil objective. To avoid bleed-through effects in the double-staining experiment, each dye was scanned independently in multi-tracking mode.

### Recombinant protein purification

Lysates from MEL cells stably expressing FLAG-Wdr82 were prepared by incubating the cells in lysis buffer [300 mM NaCl, 1% NP-40, 0.1% SDS, 0.5% sodium-deoxycholate, and 50 mM Tris-HCl (pH 8.0)] containing a protease inhibitor cocktail (Roche). Anti-FLAG immunoaffinity columns were prepared using anti-FLAG M2 affinity gel (Sigma) following the manufacturer’s instructions. Cell lysates were applied to an equilibrated FLAG column of 200-μl bed volume to allow for protein adsorption to the column resin. After binding, the column was extensively washed with high-salt solution for 5 min five times each [300 mM NaCl, 150 mM KCl, 1% NP-40, 0.1% SDS, 0.5% sodium-deoxycholate, and 50 mM Tris·HCl (pH 8.0)]. A FLAG peptide (Sigma) was applied to the column to elute the FLAG protein complex, as recommended by the manufacturer. The eluents were collected and visualized by SDS-PAGE followed by Coomassie blue staining. The purified protein was used in subsequent molecular interaction assays.

### Surface plasmon resonance (SPR)

Sensorgrams of *lncEry*-P5 binding to Wdr82 were measured by SPR technology using a Biacore 3000 instrument (GE Healthcare). The flowing-phase RNA *lncEry-P5* was synthesized by *in vitro* transcription using a T7 RNA production system (Promega, P1300) in accordance with the manufacturer’s instructions. The substrate-phase Wdr82 proteins were purified from MEL cells and immobilized on a CM5 sensor chip (BIAcore) with the following procedure. All experiments were carried out using HBS-EP (10 mM HEPES pH 7.4, 150 mM NaCl, 3.4mM EDTA, and 0.005% surfactant P20) as the running buffer at a constant flow rate of 30 μl/min at 25°C. The Wdr82 protein was diluted in 10 mM sodium acetate buffer (pH 4.0) to a final concentration of 2.5 μM and covalently immobilized on the hydrophilic carboxymethylated dextran matrix of the CM5 sensor chip using a standard primary amine coupling procedure. LncEry-P5 was dissolved in the running buffer to concentrations ranging from 36.7 nM to 294 nM. All data were analyzed by BIA evaluation software, and the sensorgrams were processed by automatic correction for nonspecific bulk refractive index effects. Kinetic analyses of lncEry-P5/Wdr82 binding were performed based on the 1:1 Langmuir binding fit model in accordance with the procedures in the software manual.

### RNA interference

All siRNA transfections were performed using Lipofectamine RNAiMAX (Invitrogen) following the manufacturer’s recommendations. The final concentration of the siRNA molecules was 10 nM, and the cells were harvested 72 h or 96 h after transfection, depending on the experiment. Control, *lncEry*, and *Wdr82* siRNAs were chemically synthesized by Sigma (Shanghai, China). The shRNAs against *lncEry, Wdr82*, or *Ddx5* in lentivirus U6-MSX-IRES vectors were purchased from GeneChem. The siRNA and shRNA sequences are provided in Supplementary Table 6.

### RNA-sequencing and analysis

RNA-Seq included three parts of experiment samples. First, a total of 16 hematopoietic cell populations were sorted from the BM of C57BL/6 mice. Second, MEP cells were sorted from the BM of *flox/flox* or Δ/Δ mice. Third, MEL cells were transfected with control or *lncEry* siRNAs. Total RNA was isolated using the RNeasy (Qiagen) purification kit. Libraries were prepared using an Illumina RNA library preparation TruSeq PE kit. High-throughput RNA-Seq was performed on an Illumina Xten sequencer (paired-end 150 bp sequencing). Clean reads were filtered by removing reads including adapters, reads including poly-N and low-quality reads from raw data. All the following analyses were based on clean data. Clean data were firstly aligned to the mouse genome (GRCm38) with GENCODE M16 gene annotation using HISAT2 (v2.1.0)^66^. DEG analyses were conducted using Cufflinks and Cuffdiff (v2.2.1) software^67^. Unsupervised hierarchical clustering was conducted using pheatmap R package. Genes with the q-value less than 0.05 in the Cuffdiff results were considered to be significantly different genes.

### Assay for Transposase-Accessible Chromatin sequencing

ATAC-seq was performed as described previously^68^, In brief, 50,000 MEP cells sorted from BM of *flox/flox* or Δ/Δ mouse cells were lysed in lysis buffer (10 mM Tris-HCl, PH 7.4, 10 mM NaCl, 3 mM MgCl_2_, 0.1% (v/v) IGPAL CA-630) for 10 min on ice. After centrifuged on 500 g for 5 min, the obtained nuclei were added to 50 μL transposition reaction buffer (offered by Vazyme TD501-01) followed with incubation at 37°C for 30 min. After tagmentation, VAHTS DNA Clean Beads were used to stop the reaction, and DNA was purified for final library construction (TruePrep DNA Library Prep Kit V2 for Illumina) before paired-end high-throughput sequencing using HiSeq XTe. Clean reads were aligned to the mouse genome (GRCm38) using BWA package, and peaks were called using MACS2 package (v2.2.5)^69^ with a false discovery rate (FDR) cutoff of 0.05.

### Cleavage under targets and tagmentation

Cut&Tag experiments were performed as described previously^50^ together with the Hyperactive TM In-Situ ChIP Library Prep Kit for Illumina from Vazyme (TD902-01). MEP cells sorted from the BM of *flox/flox* or Δ/Δ mice cells were captured by ConA beads and incubated with primary and secondary antibodies in Antibody buffer and Dig-wash Buffer, respectively, for the time indicated in the manufacturer’s instructions. The cells were then incubated with Hyperactive pA-Tn5 Transposon and fragmented in Tagmentation Buffer at 37°C for 1 h. The extracted DNA fragments were analyzed by high-throughput sequencing. Clean reads were aligned to the mouse genome (GRCm38) using the Bowtie2 package^70^, and Peaks were detected using MACS2 callpeak (v2.2.5)^69^ with an FDR cutoff of 0.05.

### Chromatin immunoprecipitation sequencing

ChIP-seq was performed as described previously^71^. In brief, ~10 million cells were crosslinked with 1% formaldehyde for 10 min at room temperature and quenched by adding glycine to a final concentration of 125 mM for 5 min. The fixed cells were resuspended in SDS lysis buffer [1% SDS, 5 mM EDTA, and 50 mM Tris-HCl (pH 8.1)] in the presence of protease inhibitors and subjected to sonication (Bioruptor, Diagenode) to generate chromatin fragments of ~300 bp in length. For immunoprecipitation, after dilution, the chromatin was incubated with control or specific antibodies (3 μg) overnight at 4°C with constant rotation; 50 μL of 50% (vol/vol) protein G magnetic beads was then added, and the incubation was continued for an additional 2 h. The beads were washed with the following buffers: TSE I, TSE II, buffer III, and Tris-EDTA buffer as described previously^71^. Then the cross-links were removed from the pulled-down chromatin complex together with the input sample by incubation at 65°C for 2 h in elution buffer. The eluted DNA was purified using a PCR purification kit (Qiagen) and analyzed by qPCR using the primers detailed in Supplementary Table 6 or by high-throughput sequencing. Clean reads were aligned to the mouse genome (GRCm38) using the Bowtie2 package^70^. ChIP-seq peaks were detected using MACS2 callpeak (v2.2.5)^69^ with a minimum FDR cutoff of 0.05.,

### Chromatin Isolation by RNA purification sequencing

ChIRP-seq was performed as described previously^48^. Eluted DNA was purified using a PCR purification kit (Qiagen) and analyzed by qPCR using the qChIP primers described in Supplementary Table 6 or by high-throughput sequencing. The processing of sequencing data was the same as that for the ChIP-seq data described above. Peaks were annotated using the ChIPseeker R package (v1.24.0)^72^.

### Luciferase reporter assay

The modulation of globin gene promoter activity by *lncEry* was analyzed by luciferase assay. A pGL3 construct carrying the globin gene CRRs sequence was obtained by PCR enrichment using the primers (Supplementary Table 6), and the *Klf1* CRRs sequence was synthesized. Luciferase reporter activity was measured using the Dual Luciferase Assay System (E1910, ProMega). Relative luciferase activity was normalized to Renila luciferase and control vector luciferase reporter activity.

### Statistical analyses

Data from biological triplicate experiments are presented, and the data represent the means ± S.D. A two-tailed unpaired Student’s t-test was used to compare two groups of data. Analysis of variance (ANOVA) with Bonferroni’s correction was used to compare multiple groups of data. A *P* value <0.05 was considered statistically significant. All statistical analyses were performed in SPSS 20.0 software, and graphs were generated by GraphPad Prism 8.0 software.

## Data availability

RNA-seq datasets of 12 hematopoietic cell populations were deposited in the NCBI GEO (GSE142216). Other sequencing data have been deposited in the NCBI SRA (PRJNA647682).

## Acknowledgments

The authors would like to thank all lab members for their assistance with the experiments. They thank Dr. Yiran Ren for providing help on SPR assays. This work was supported by grants from the Ministry of Science and Technology of China (2017YFA0103400, 2020YFE0203000, 2016YFA0100600), the National Natural Science Foundation of China (81922002, 81730006, 81861148029, 81890990, 81870086, 81900113, 81890993, 81900117, 81920108004), the CAMS Initiative for Innovative Medicine (2017-I2M-3-009, 2016-I2M-1-017), the CAMS Fundamental Research Funds for Central Research Institutes (2019RC310003), Distinguished Young Scholars of Tianjin (19JCJQJC63400).

## Author contributions

SD.Y., GH.S. and C.C., designed and performed the experiments, analyzed the data, and wrote the paper. P. W. performed bioinformatic analyses. YJ.K., ZF.Z., YC.H., Q.G., FJ.W., FL.G., ZN.Y., X.L., XN.Z, and SH.L. helped with mouse experiments. T.L., CY.Z., and L.Y. helped with bioinformatic analyses. YP.L. helped with experimental design. YN.M., J.Y., SR.Y., F.D., LH.S., and LG.N. assisted with the manuscript. J.L., H.C., and T.C. proposed the study, designed the experiments, interpreted the results, wrote the paper, and oversaw the research project.

