## Supplementary Figures and Tables for "WDR82-binding long non-coding RNA *lncEry* controls mouse erythroid differentiation and maturation": Supplemental Files.pdf

### Figure S1

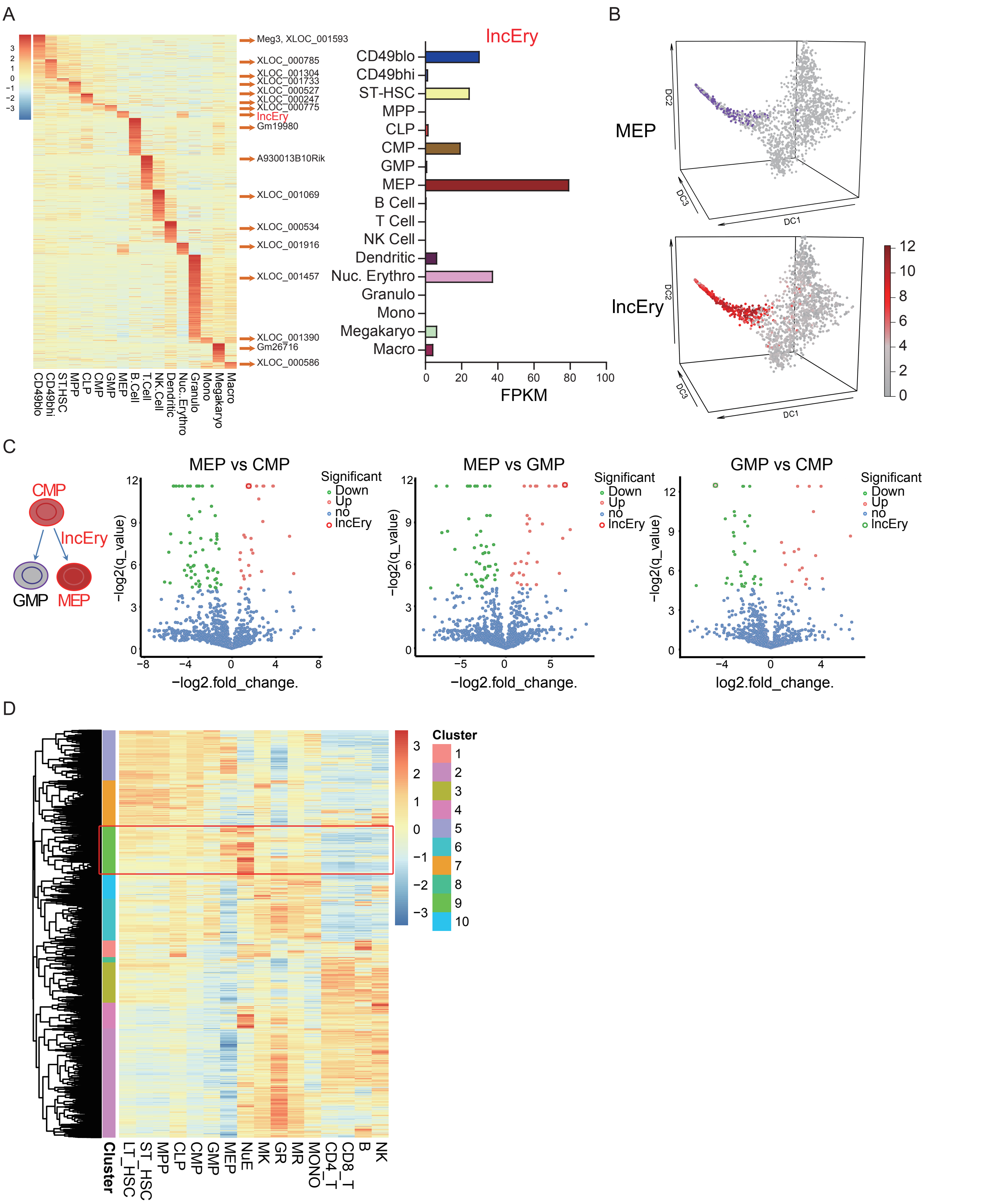
A
Meg3, XLOC_001593
XLOC_000785
XLOC_001304
XLOC_001733
XLOC_000527
XLOC_000247
XLOC_000775
lncEry
Gm19980
A930013B10Rik
XLOC_001069
XLOC_000534
XLOC_001916
XLOC_001457
XLOC_001390
Gm26716
XLOC_000586
CD49blo
CD49bhi
ST.HSC
MPP
CLP
CMP
GMP
MEP
B.Cell
T.Cell
NK.Cell
Dendritic
Nuc..Erythro
Granulo
Mono
Megakaryo
Macro
lncEry
CD49blo
CD49bhi
ST-HSC
MPP
CLP
CMP
GMP
MEP
B Cell
T Cell
NK Cell
Dendritic
Nuc. Erythro
Granulo
Mono
Megakaryo
Macro
FPKM
B
MEP
lncEry
DC1
DC2
DC3
C
CMP
lncEry
GMP
MEP
MEP vs CMP
MEP vs GMP
GMP vs CMP
Significant
Down
Up
no
lncEry
−log2(q_value)
−log2.fold_change.
log2.fold_change.
D
Cluster
LT_HSC
ST_HSC
MPP
CLP
CMP
GMP
MEP
NuE
MK
GR
MR
MONO
CD4_T
CD8_T
B
NK

### Figure S2

A

NONMMUG004428.1 (lncEry isoform-1) sequence: 908 bp

AGGGGACACGCTGGGTCCTGCGGATATGGATATCCTGAAAACACAGGCCCAGCTCACTGGGAAGTGAGCTGTCCACGGGAGACCACAGGAGACCACAG
GCAGCATTGTGGGGATAGACTGAAAACATCTTGGGAAATAGTCGCAGTGTCACAAGAACCACCCTGGCGCTTTGTTTAATGCATCTTTTGAGCCCCTGCG
GAGACCCCGTGCTGAGTTCCACAGTGCAGTATCAAAAGAGAGGAGACACCTTAATCTGATACTCTGGGAAGGGGATCTAGCCTGTATCAGGAAGCAGAG
AGGGAGCCAATCCAGATGCTTCAAAAGTCCTGGCAGGTGGCCCTTGGGTCCCCACCCAGAAGAAGCCTGTGTGTCACGACTGGAGAGAGGAGAGGCAC
TGTGGAAGCCTTCCATTTCCCAGAAGCCTCTTCTTGTTTTGTCCCCAGGAAGAGGGTATTCCTGAGCTTTCTCCTGAGATGCAATCAAGCTACGTGGCACA
AGCTGGGTCCTTTTGAGGCTTACTTTTAAGTTTTGTTAGGTTCTGAACTCAGAGACTCCATCAGGCTCCATCTATGCTCCCCCTTCCTGCATCAGCATCCGG
TCACTTTCTCCAGACAGTCAGAGAGGTGATCTCAGGGCCTACCCCCAAGCATCATTGTCCTGCCCCGCCTGATGTCCAAAGGCCTTGAAACTGTTGTTAGC
AATCAGAGGTCTTCACCACCTGCCCCAGGGACCCAGCAACAGCAGGATACATTATCACAGCTGCCAAGATCATCACGAGTAGCAGATCCTCACTACATCTT
CAGATAAACAGCCAGTCCCCCCTCTCTTTGCTCCCTTTTTCCCTCCACCCTCGTTGAGAGGCAGCCTCTGTATACCCTTTCCCCAATAAACTTCCTATGTGA
GATCTG

NONMMUG004428.2 (lncEry isoform-2) sequence: 961 bp

ATTTGGATAGCCAGGAGAGAAGAGCAGCCTGGAAGACCATCTGCAACTGTCTGCAGGGGACACGCTGGGTCCTGCGGATATGGATATCCTGAAAACACA
GGCCCAGCTCACTGGGAAGTGAGCTGTCCACGGGAGACCACAGGAGACCACAGGCAGCATTGTGGGGATAGACTGAAAACATCTTGGGAAATAGTCGC
AGTGTCACAAGAACCACCCTGGCGCTTTGTTTAATGCATCTTTTGAGCCCCTGCGGAGACCCCGTGCTGAGTTCCACAGTGCAGTATCAAAAGAGAGGAG
ACACCTTAATCTGATACTCTGGGAAGGGGATCTAGCCTGTATCAGGAAGCAGAGAGGGAGCCAATCCAGATGCTTCAAAAGTCCTGGCAGGTGGCCCTTG
GGTCCCCACCCAGAAGAAGCCTGTGTGTCACGACTGGAGAGAGGAGAGGCACTGTGGAAGCCTTCCATTTCCCAGAAGCCTCTTCTTGTTTTGTCCCCAG
GAAGAGGGTATTCCTGAGCTTTCTCCTGAGATGCAATCAAGCTACGTGGCACAAGCTGGGTCCTTTTGAGGCTTACTTTTAAGTTTTGTTAGGTTCTGAACT
CAGAGACTCCATCAGGCTCCATCTATGCTCCCCCTTCCTGCATCAGCATCCGGTCACTTTCTCCAGACAGTCAGAGAGGTGATCTCAGGGCCTACCCCCAA
GCATCATTGTCCTGCCCCGCCTGATGTCCAAAGGCCTTGAAACTGTTGTTAGCAATCAGAGGTCTTCACCACCTGCCCCAGGGACCCAGCAACAGCAGGA
TACATTATCACAGCTGCCAAGATCATCACGAGTAGCAGATCCTCACTACATCTTCAGATAAACAGCCAGTCCCCCCTCTCTTTGCTCCCTTTTTCCCTCCAC
CCTCGTTGAGAGGCAGCCTCTGTATACCCTTTCCCCAATAAACTTCCTATGTGAGATCTG

lncEry isoform-3 sequence: 908 bp

AAAGAACATCCCTCCATGGCCTCTGCATCAGCTCCTGCTTCCTGACCTGCTTGAGTTCCAGTCCTGACTTCCTTGGTGATGAACAACAGTATGGAAATGCT
TGCCGGTGTATGAAATTAAAGAATTGAGTGCATTCAGGATTTGGATAGCCAGGAGAGAAGAGCAGCCTGGAAGACCATCTGCAACTGTCTGGGATAGACT
GAAAACATCTTGGGAAATAGTCGCAGTGTCACAAGAACCACCCTGGCGCTTTGTTTAATGCATCTTTTGAGCCCCTGCGGAGACCCCGTGCTGAGTTCCA
CAGTGCAGTATCAAAAGAGAGGAGACACCTTAATCTGATACTCTGGGAAGGGGATCTAGCCTGTATCAGGAAGCAGAGAGGGAGCCAATCCAGATGCTTC
AAAAGTCCTGGCAGGTGGCCCTTGGGTCCCCACCCAGAAGAAGCCTGTGTGTCACGACTGGAGAGAGGAGAGGCACTGTGGAAGCCTTCCATTTCCCAG
AAGCCTCTTCTTGTTTTGTCCCCAGGAAGAGGGTTCTGAACTCAGAGACTCCATCAGGCTCCATCTATGCTCCCCCTTCCTGCATCAGCATCCGGTCACTTT
CTCCAGACAGTCAGAGAGGTGATCTCAGGGCCTACCCCCAAGCATCATTGTCCTGCCCCGCCTGATGTCCAAAGGCCTTGAAACTGTTGTTAGCAATCAGA
GGTCTTCACCACCTGCCCCAGGGACCCAGCAACAGCAGGATACATTATCACAGCTGCCAAGATCATCACGAGTAGCAGATCCTCACTACATCTTCAGATAA
ACAGCCAGTCCCCCCTCTCTTTGCTCCCTTTTTCCCTCCACCCTCGTTGAGAGGCAGCCTCTGTATACCCTTTCCCCAATAAACTTCCTATGTGAGATCTGT
CA

B

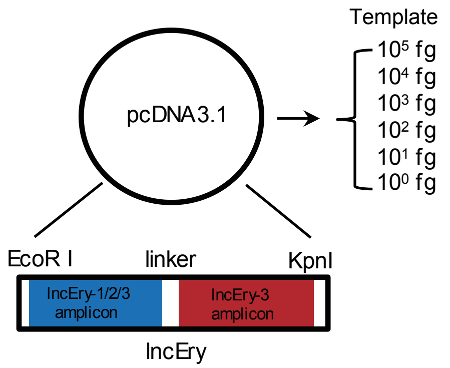

C

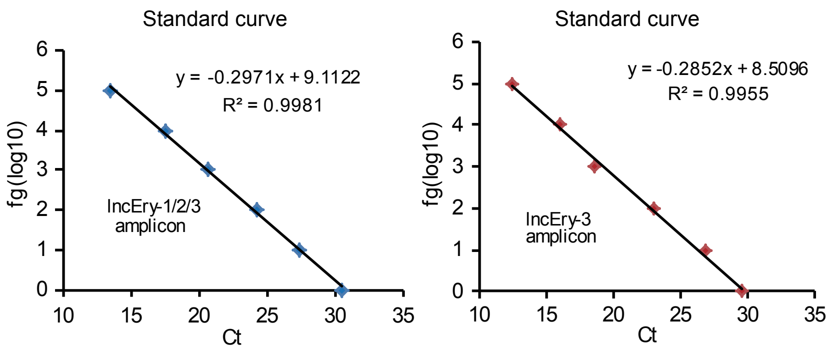

D

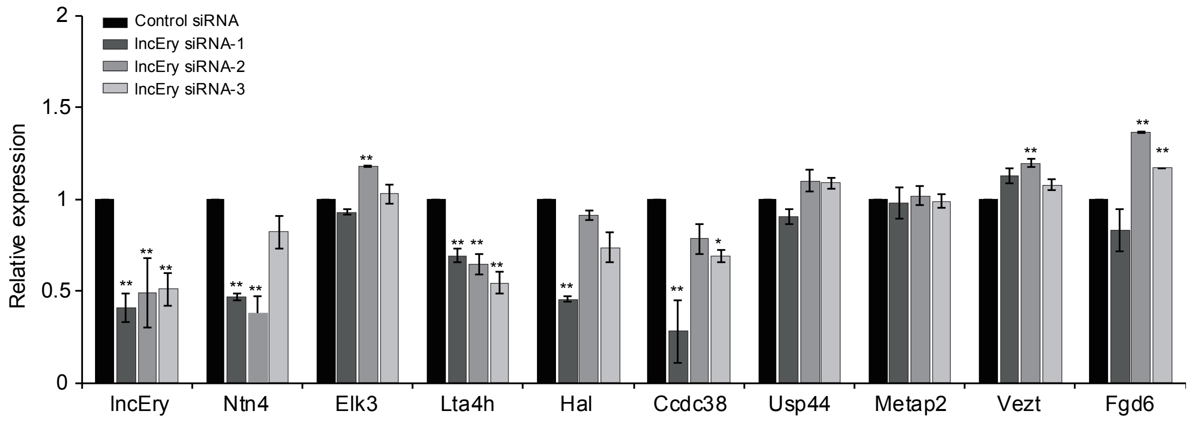

Figure S3

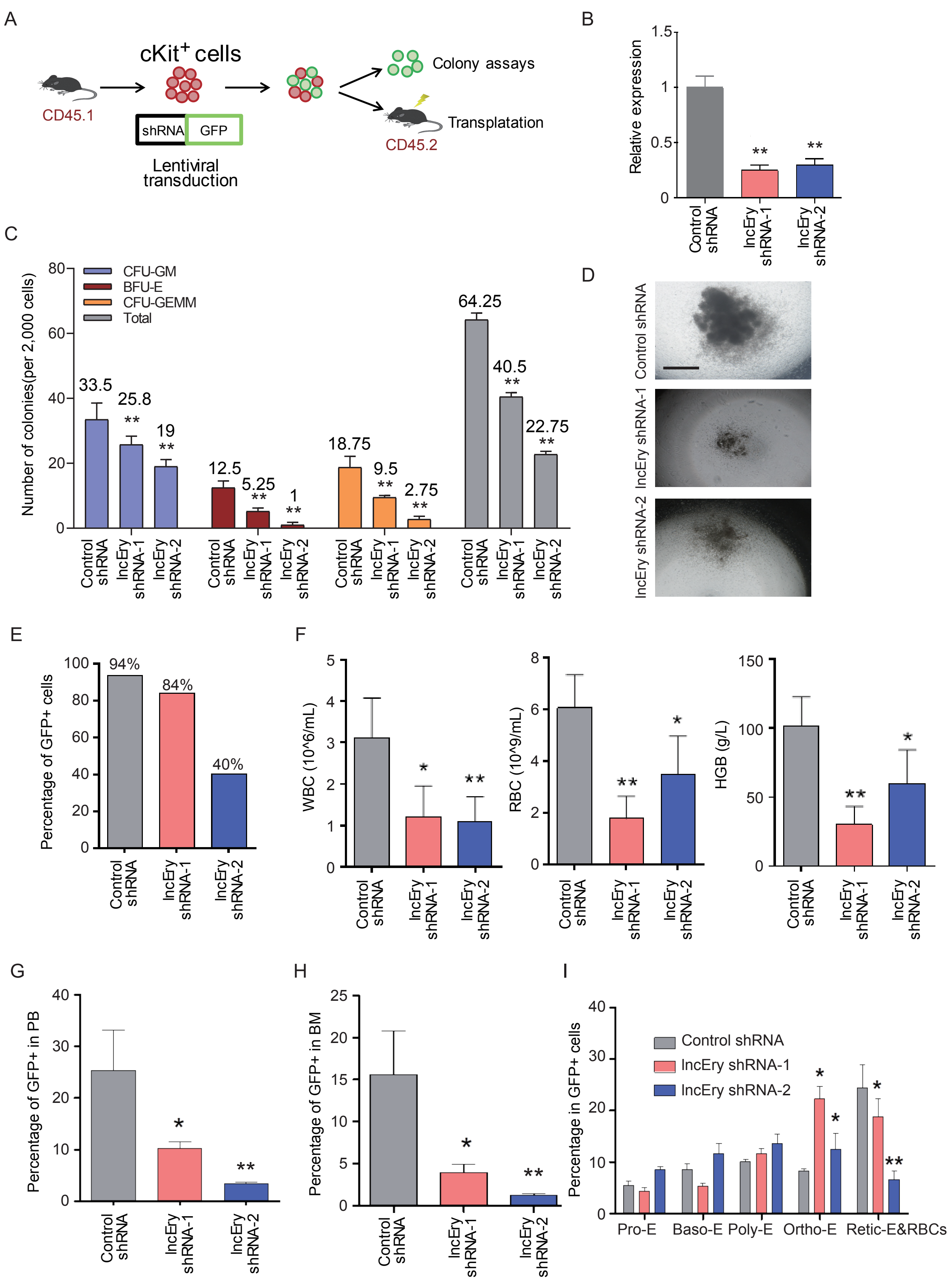
A
cKit+ cells
CD45.1
shRNA
GFP
Lentiviral transduction
Colony assays
Transplatation
CD45.2
B
Relative expression
Control shRNA
lncEry shRNA-1
lncEry shRNA-2
C
Number of colonies(per 2,000 cells)
CFU-GM
BFU-E
CFU-GEMM
Total
33.5
25.8
19
12.5
5.25
1
18.75
9.5
2.75
64.25
40.5
22.75
D
Control shRNA
lncEry shRNA-1
lncEry shRNA-2
E
Percentage of GFP+ cells
94%
84%
40%
F
WBC (10^6/mL)
RBC (10^9/mL)
HGB (g/L)
G
Percentage of GFP+ in PB
H
Percentage of GFP+ in BM
I
Percentage in GFP+ cells
Pro-E
Baso-E
Poly-E
Ortho-E
Retic-E&RBCs

### Figure S4

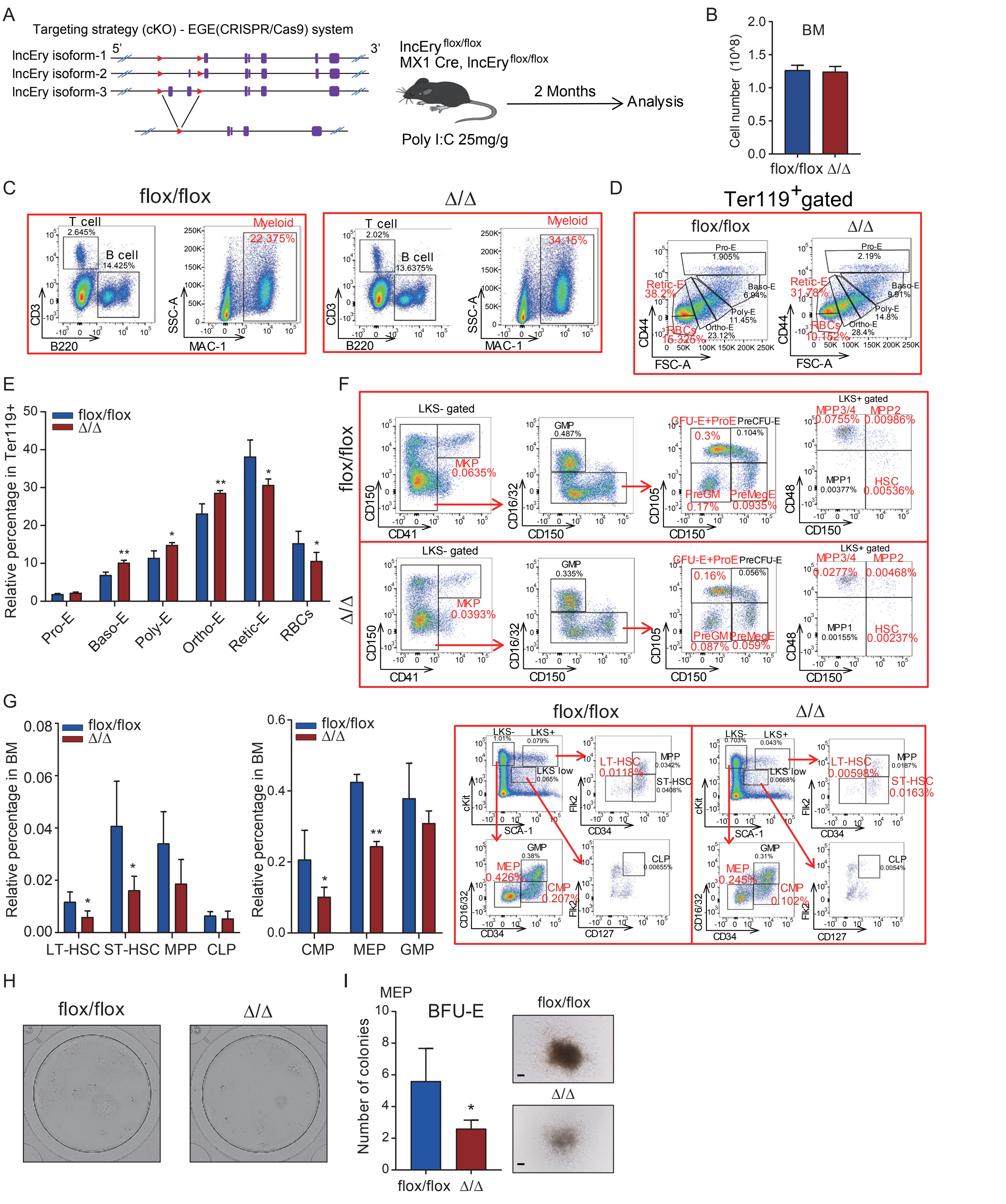
A
Targeting strategy (cKO) - EGE(CRISPR/Cas9) system
lncEry isoform-1
lncEry isoform-2
lncEry isoform-3
lncEry flox/flox
MX1 Cre, lncEry flox/flox
Poly I:C 25mg/g
2 Months
Analysis
B
BM
Cell number (10^8)
flox/flox Δ/Δ
C
flox/flox
Δ/Δ
T cell
B cell
Myeloid
B220
MAC-1
CD3
SSC-A
D
Ter119+gated
Pro-E
Baso-E
Poly-E
Ortho-E
Retic-E
RBCs
FSC-A
CD44
E
Relative percentage in Ter119+
F
LKS- gated
LKS+ gated
MKP
GMP
CFU-E+ProE
PreCFU-E
PreGM
PreMegE
MPP3/4
MPP2
MPP1
HSC
CD41
CD150
CD16/32
CD105
CD48
G
Relative percentage in BM
LT-HSC ST-HSC MPP CLP
CMP MEP GMP
LKS-
LKS+
LKS low
LT-HSC
ST-HSC
CLP
SCA-1
cKit
CD34
Flk2
CD127
H
flox/flox
Δ/Δ
I
MEP
BFU-E
Number of colonies

### Figure S5

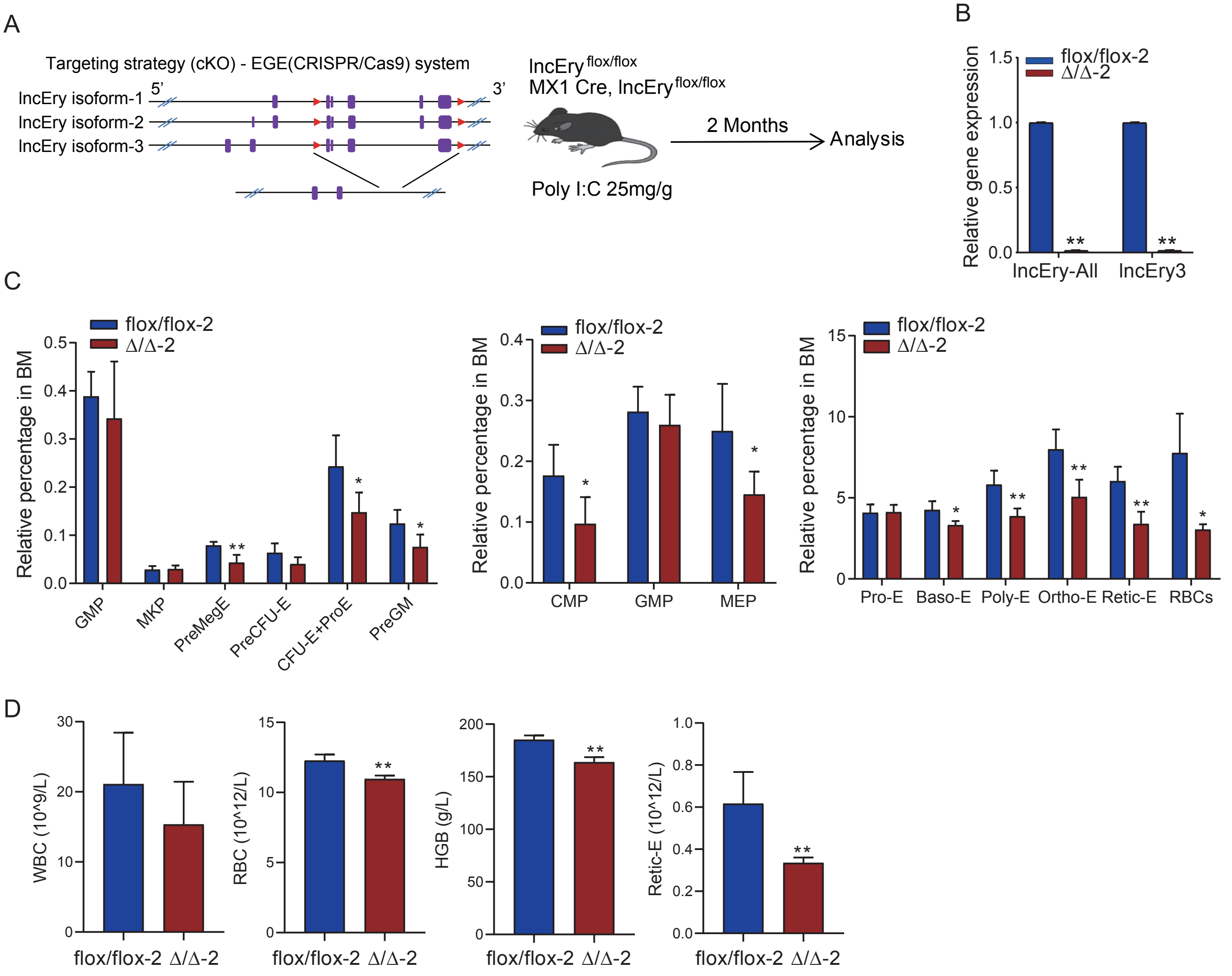
A
Targeting strategy (cKO) - EGE(CRISPR/Cas9) system
lncEry isoform-1
lncEry isoform-2
lncEry isoform-3
5'
3'
lncEry flox/flox
MX1 Cre, lncEry flox/flox
Poly I:C 25mg/g
2 Months
Analysis
B
Relative gene expression
flox/flox-2
Δ/Δ-2
lncEry-All
lncEry3
C
Relative percentage in BM
GMP
MKP
PreMegE
PreCFU-E
CFU-E+ProE
PreGM
CMP
MEP
Pro-E
Baso-E
Poly-E
Ortho-E
Retic-E
RBCs
D
WBC (10^9/L)
RBC (10^12/L)
HGB (g/L)
Retic-E (10^12/L)
flox/flox-2
Δ/Δ-2

### Figure S6

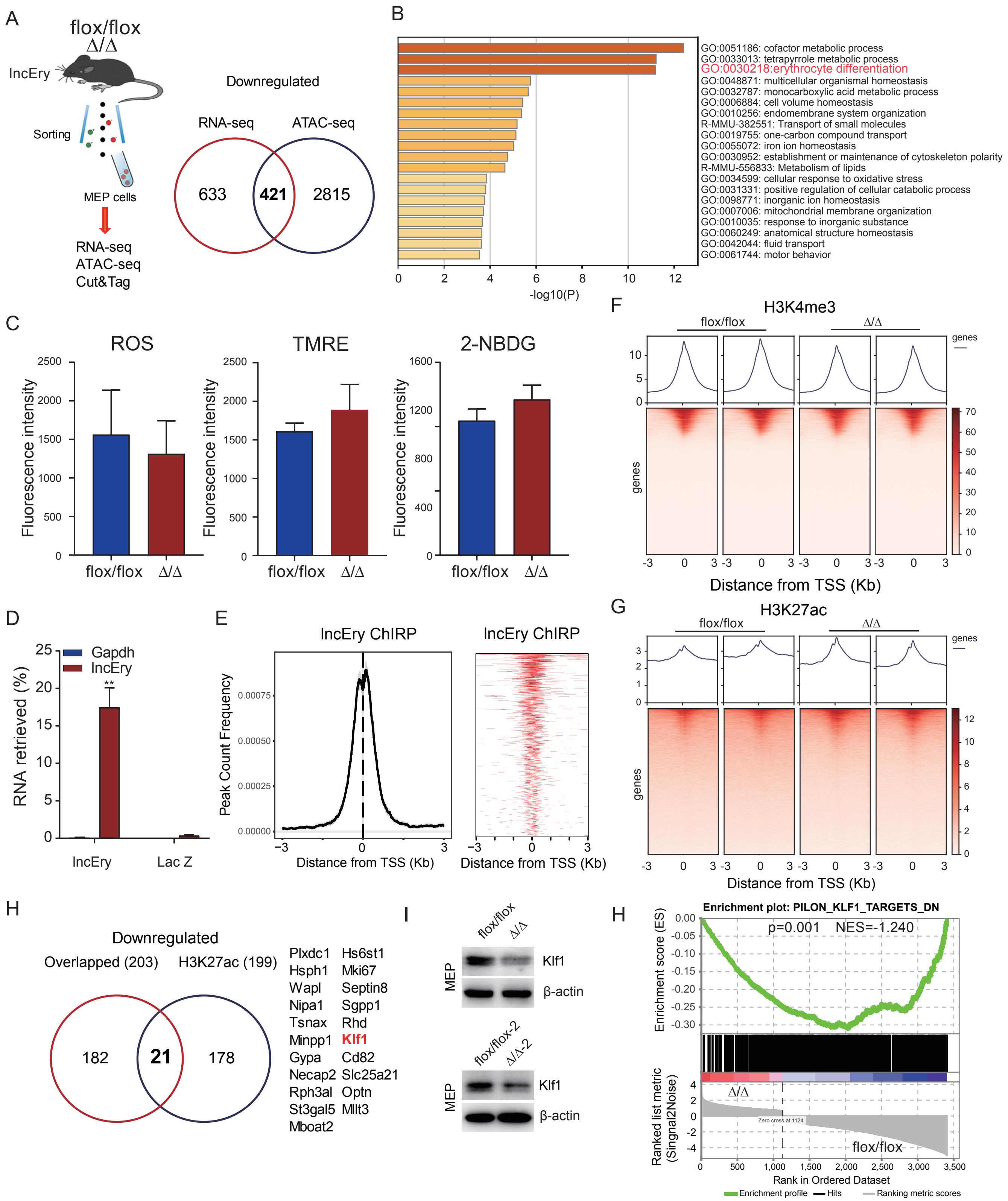
A
flox/flox
Δ/Δ
lncEry
Sorting
MEP cells
RNA-seq
ATAC-seq
Cut&Tag
Downregulated
RNA-seq
ATAC-seq
633
421
2815
B
GO:0051186: cofactor metabolic process
GO:0033013: tetrapyrrole metabolic process
GO:0030218:erythrocyte differentiation
GO:0048871: multicellular organismal homeostasis
GO:0032787: monocarboxylic acid metabolic process
GO:0006884: cell volume homeostasis
GO:0010256: endomembrane system organization
R-MMU-382551: Transport of small molecules
GO:0019755: one-carbon compound transport
GO:0055072: iron ion homeostasis
GO:0030952: establishment or maintenance of cytoskeleton polarity
R-MMU-556833: Metabolism of lipids
GO:0034599: cellular response to oxidative stress
GO:0031331: positive regulation of cellular catabolic process
GO:0098771: inorganic ion homeostasis
GO:0007006: mitochondrial membrane organization
GO:0010035: response to inorganic substance
GO:0060249: anatomical structure homeostasis
GO:0042044: fluid transport
GO:0061744: motor behavior
-log10(P)
C
ROS
TMRE
2-NBDG
Fluorescence intensity
flox/flox
Δ/Δ
F
H3K4me3
genes
Distance from TSS (Kb)
D
Gapdh
lncEry
RNA retrieved (%)
**
lncEry
Lac Z
E
lncEry ChIRP
lncEry ChIRP
Peak Count Frequency
Distance from TSS (Kb)
Distance from TSS (Kb)
G
H3K27ac
flox/flox
Δ/Δ
genes
Distance from TSS (Kb)
H
Downregulated
Overlapped (203)
H3K27ac (199)
182
21
178
Plxdc1
Hsph1
Wapl
Nipa1
Tsnax
Minpp1
Gypa
Necap2
Rph3al
St3gal5
Mboat2
Hs6st1
Mki67
Septin8
Sgpp1
Rhd
Klf1
Cd82
Slc25a21
Optn
Mllt3
I
flox/flox
Δ/Δ
MEP
Klf1
β-actin
flox/flox-2
Δ/Δ-2
MEP
Klf1
β-actin
H
Enrichment plot: PILON_KLF1_TARGETS_DN
p=0.001
NES=-1.240
Enrichment score (ES)
Δ/Δ
Zero cross at 1124
flox/flox
Ranked list metric (Singnal2Noise)
Rank in Ordered Dataset
Enrichment profile
Hits
Ranking metric scores

### Figure S7

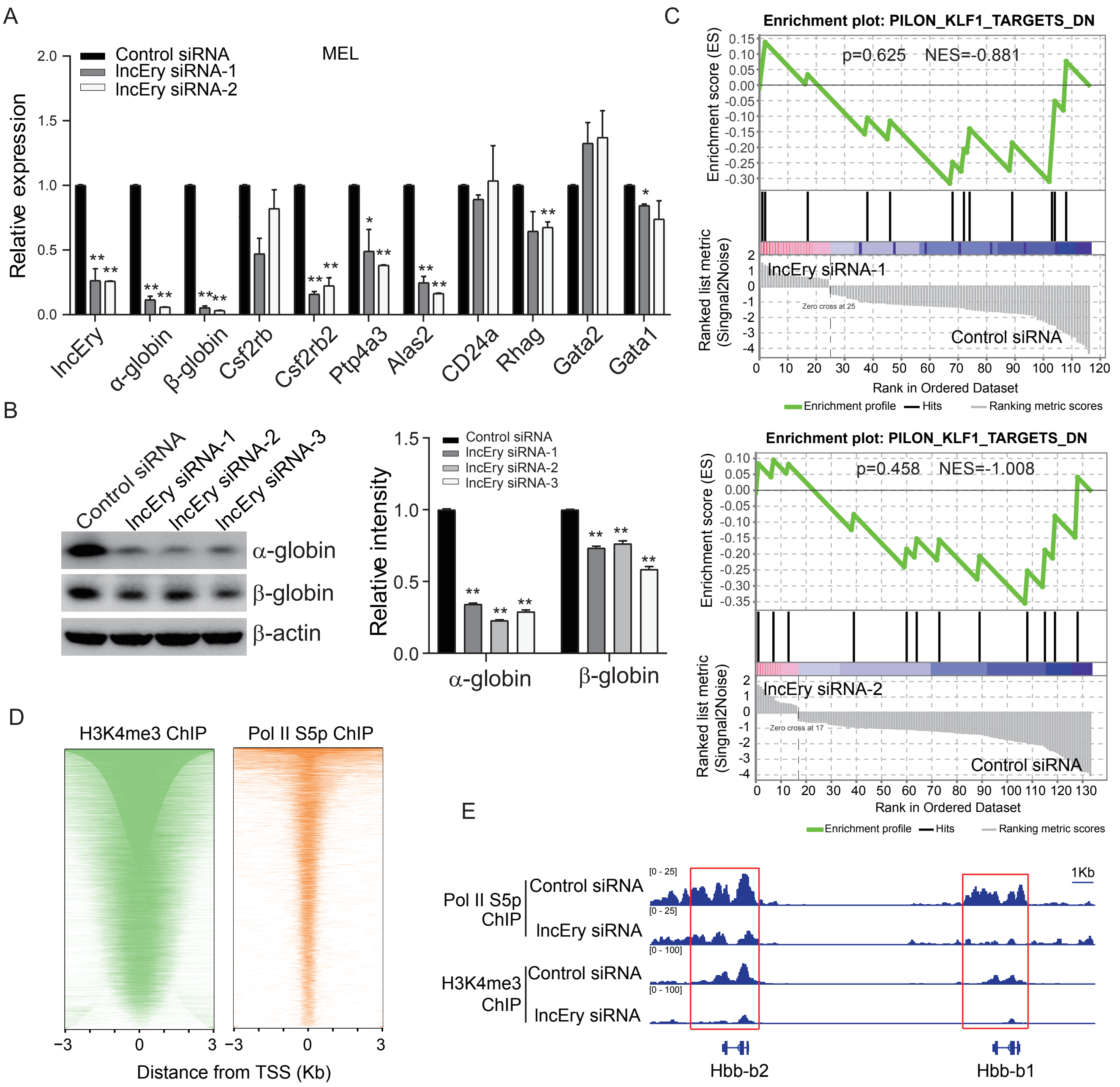
A
MEL
Control siRNA
lncEry siRNA-1
lncEry siRNA-2
Relative expression
lncEry
α-globin
β-globin
Csf2rb
Csf2rb2
Ptp4a3
Alas2
CD24a
Rhag
Gata2
Gata1
B
Control siRNA
lncEry siRNA-1
lncEry siRNA-2
lncEry siRNA-3
α-globin
β-globin
β-actin
Relative intensity
C
Enrichment plot: PILON_KLF1_TARGETS_DN
p=0.625 NES=-0.881
Enrichment score (ES)
Ranked list metric (Singnal2Noise)
lncEry siRNA-1
Zero cross at 25
Control siRNA
Rank in Ordered Dataset
Enrichment profile
Hits
Ranking metric scores
Enrichment plot: PILON_KLF1_TARGETS_DN
p=0.458 NES=-1.008
lncEry siRNA-2
Zero cross at 17
Control siRNA
Rank in Ordered Dataset
D
H3K4me3 ChIP
Pol II S5p ChIP
Distance from TSS (Kb)
E
Pol II S5p ChIP
H3K4me3 ChIP
Control siRNA
lncEry siRNA
[0 - 25]
[0 - 100]
1Kb
Hbb-b2
Hbb-b1

Figure S8

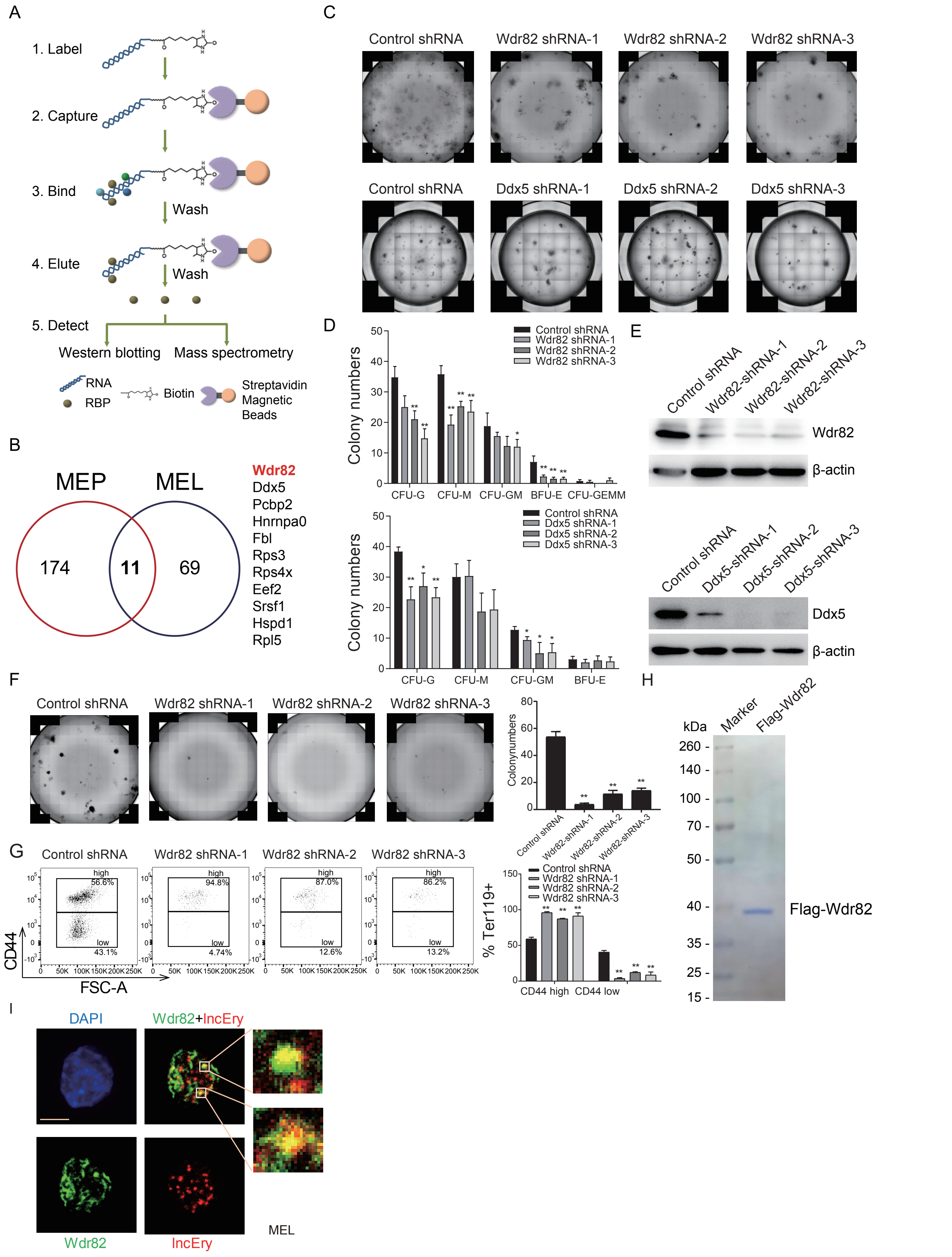
A
1. Label
2. Capture
3. Bind
Wash
4. Elute
Wash
5. Detect
Western blotting
Mass spectrometry
RNA
RBP
Biotin
Streptavidin Magnetic Beads
B
MEP
MEL
174
11
69
Wdr82
Ddx5
Pcbp2
Hnrnpa0
Fbl
Rps3
Rps4x
Eef2
Srsf1
Hspd1
Rpl5
C
Control shRNA
Wdr82 shRNA-1
Wdr82 shRNA-2
Wdr82 shRNA-3
Control shRNA
Ddx5 shRNA-1
Ddx5 shRNA-2
Ddx5 shRNA-3
D
Colony numbers
Control shRNA
Wdr82 shRNA-1
Wdr82 shRNA-2
Wdr82 shRNA-3
CFU-G
CFU-M
CFU-GM
BFU-E
CFU-GEMM
Colony numbers
Control shRNA
Ddx5 shRNA-1
Ddx5 shRNA-2
Ddx5 shRNA-3
CFU-G
CFU-M
CFU-GM
BFU-E
E
Control shRNA
Wdr82-shRNA-1
Wdr82-shRNA-2
Wdr82-shRNA-3
Wdr82
β-actin
Control shRNA
Ddx5-shRNA-1
Ddx5-shRNA-2
Ddx5-shRNA-3
Ddx5
β-actin
F
Control shRNA
Wdr82 shRNA-1
Wdr82 shRNA-2
Wdr82 shRNA-3
Colonynumbers
Control shRNA
Wdr82-shRNA-1
Wdr82-shRNA-2
Wdr82-shRNA-3
G
Control shRNA
Wdr82 shRNA-1
Wdr82 shRNA-2
Wdr82 shRNA-3
high
56.6%
low
43.1%
high
94.8%
low
4.74%
high
87.0%
low
12.6%
high
86.2%
low
13.2%
CD44
FSC-A
% Ter119+
Control shRNA
Wdr82 shRNA-1
Wdr82 shRNA-2
Wdr82 shRNA-3
CD44 high
CD44 low
H
kDa
Marker
Flag-Wdr82
260 -
140 -
100 -
70 -
50 -
40 -
35 -
25 -
15 -
Flag-Wdr82
I
DAPI
Wdr82+lncEry
Wdr82
lncEry
MEL

### Figure S9

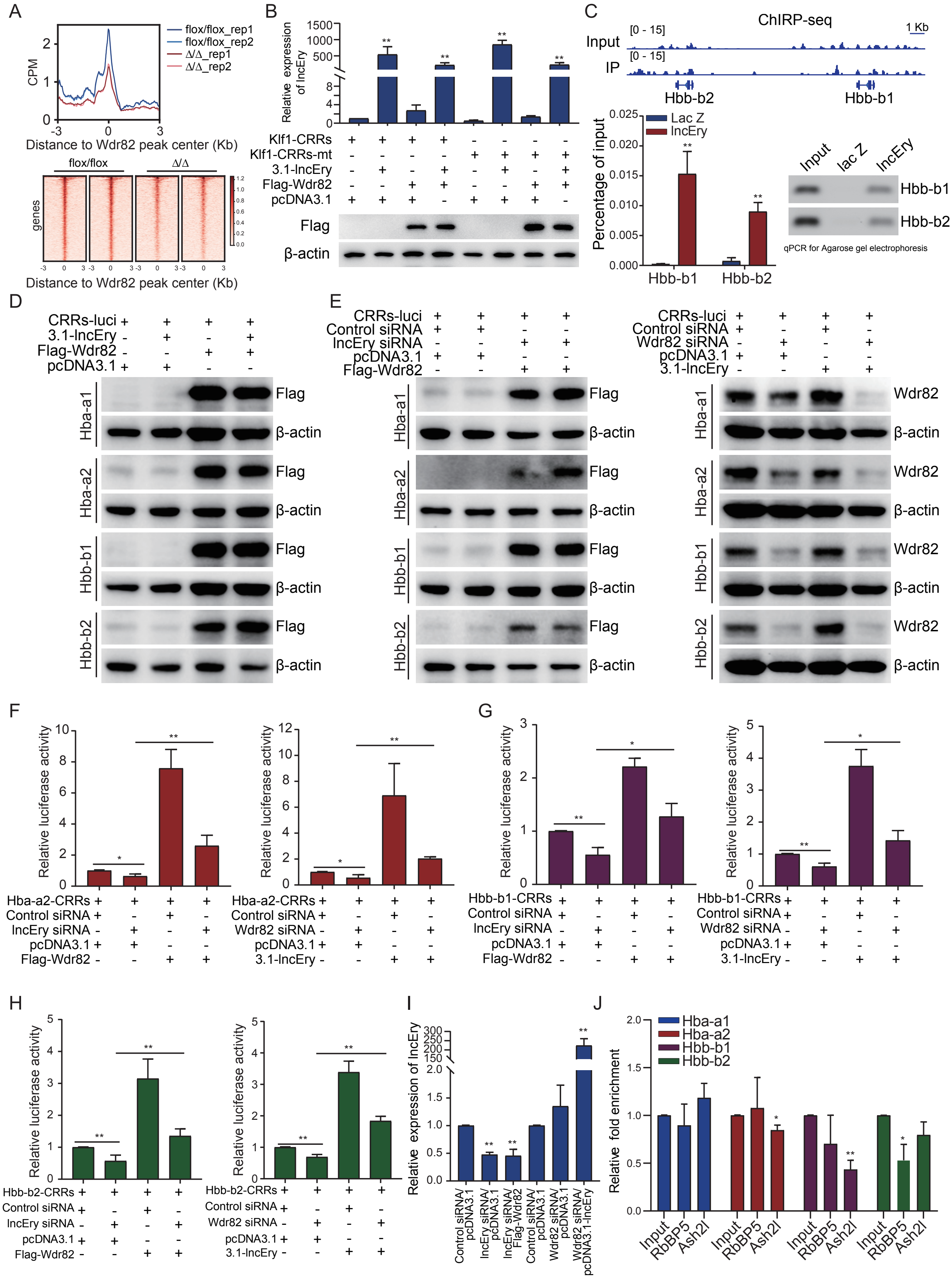
A
flox/flox_rep1
flox/flox_rep2
Δ/Δ_rep1
Δ/Δ_rep2
CPM
Distance to Wdr82 peak center (Kb)
flox/flox
Δ/Δ
genes
Distance to Wdr82 peak center (Kb)
B
Relative expression of lncEry
Klf1-CRRs
Klf1-CRRs-mt
3.1-lncEry
Flag-Wdr82
pcDNA3.1
Flag
β-actin
C
ChIRP-seq
Input
IP
1 Kb
Hbb-b2
Hbb-b1
Lac Z
lncEry
Percentage of input
Input
lac Z
lncEry
qPCR for Agarose gel electrophoresis
D
CRRs-luci
3.1-lncEry
Flag-Wdr82
pcDNA3.1
Hba-a1
Hba-a2
Hbb-b1
Hbb-b2
E
Control siRNA
lncEry siRNA
Wdr82 siRNA
Wdr82
F
Relative luciferase activity
Hba-a2-CRRs
G
Hbb-b1-CRRs
H
Hbb-b2-CRRs
I
Relative expression of lncEry
Control siRNA/pcDNA3.1
lncEry siRNA/pcDNA3.1
lncEry siRNA/Flag-Wdr82
Control siRNA/pcDNA3.1
Wdr82 siRNA/pcDNA3.1
Wdr82 siRNA/pcDNA3.1-lncEry
J
Hba-a1
Hba-a2
Hbb-b1
Hbb-b2
Relative fold enrichment
Input
RbBP5
Ash2l

**Supplementary Figure Legends**

**Figure S1. Gm15915 is highly expressed in erythroid lineage**

(A) Heat map of differentially expressed lncRNAs in 17 hematopoietic cell populations and representative lncRNAs in different cell types. Histogram shows the expression of *lncEry* in each cell population. (B) Diffusion map of all cells colored according to the expression of *lncEry* genes. Diffusion of all cells and MEP cells is shown in purple. The color corresponds to a log2 scale of expression ranging from 0 to the maximum value for the gene. Diffusion components 1, 2, and 3 are shown. (C) Volcano plots comparing the differentially expressed genes (dots) between MEPs and CMPs, MEPs and GMPs, or GMPs and CMPs. Red or green dots with black circles show representative upregulated and downregulated genes, respectively. Gm15915 is indicated by larger dots. (D) Heat map and unsupervised hierarchical clustering of differentially expressed genes in 16 hematopoietic cell populations. Ten cluster types were grouped according to the transcriptome profiles.

**Figure S2. LncEry is a *bona fide* long non-coding RNA**

(A) Nucleotide sequences of *lncEry* isoforms. (B) Schematic of approach used to quantify the relative expression of *lncEry* isoform-1/2/3 and isoform-3. Both *lncEry* isoform-1/2/3 and isoform-3 amplicons were cloned in tandem into the same plasmid. Five different dilutions were made for the qPCR to generate the standard curve. The amount of *lncEry* isoform-1/2/3 and isoform-3 was calculated by fitting the Ct value to the respective standard curve. The amount of lncEry isoform-1/2 was calculated by subtracting the value of *lncEry* isoform-3 from the value of lncEry isoform-1/2/3. (C) Representative standard curves for *lncEry* isoform-1/2/3 and isoform-3. (D) MEL cells were transfected with control or *lncEry* siRNA, and the expression levels of indicated genes were analyzed by qPCR. Each error bar represents the mean ± S.D. for triplicate experiments. **P < 0.01.

**Figure S3. Erythroid differentiation is impaired by *lncEry* knockdown in HSPCs**

(A) Schematic of the experimental procedure of gene knockdown. Briefly, GFP-fused control or *lncEry* shRNA lentiviruses were transduced into donor ($CD45.1^+$) murine $cKit^+$

HSPCs, which were injected into lethally irradiated (9.5Gy) recipient ($CD45.2^+$) mice with the indicated $GFP^+$ percentage, or $GFP^+$ cells were sorted and analyzed by colony assay. (B) $cKit^+$ cells stably expressing different sets of *lncEry* shRNAs were collected for qPCR analysis (n = 4). (C) $GFP^+$ cells were cultured for 10-14 days for CFU assays in complete methylcellulose-based medium, and colonies were counted (n = 4). (D) Colony assays of $GFP^+$ cells transfected with control or *lncEry* shRNAs. Representative images from triplicate experiments are shown. Scale bar, 200 µm. (E) Percentage of $GFP^+$ cells before transplantation into recipient mice (n = 5-7 mice per group). (F) Absolute numbers or concentrations of indicated items in peripheral blood (PB) 21 days after transplantation (n = 3-7 mice per group). (G-H) Percentage of $GFP^+$ cells in recipient mice PB or bone marrow (BM) 21 days after transplantation (n = 3-7 mice per group). (I) Percentage of proerythroblast (Pro-E), basophilic erythroblasts (Baso-E), polychromatic erythroblasts (Poly-E), orthochromatic erythroblasts (Ortho-E), reticulocytes (Retic-E) or red blood cells (RBCs) in $GFP^+$ cells of recipient mice BM (n = 3-7 mice per group). Each error bar represents the mean ± S.D. *$P < 0.05$, **$P < 0.01$.

**Figure S4. Erythroid differentiation is impaired in *lncEry* Δ/Δ mouse**

(A) Schematic of the *loxP* sequence integration into exons 1-2 of *lncEry* isoform-3 using CRISPR/Cas9 technology (left). Schematic of conditional knockout (cKO) mice induced with MX1-Cre for 2 months before flow cytometry (right). (B) Cell numbers in *flox/flox* and Δ/Δ mouse bone marrow (BM). (C) Gating strategies for FACS analysis of T, B, and myeloid cells. (D) Plot of CD44 versus FSC of the Ter119-positive cells with gating for proerythroblast (Pro-E), basophilic erythroblast (Baso-E), polychromatic erythroblast (Poly-E), orthochromatic erythroblast (Ortho-E), reticulocyte (Retic-E), or red blood cell (RBC) populations. (E) Percentage of indicated populations in $Ter119^+$ cells from control and *lncEry* cKO mice. (F) Gating strategies for FACS of the indicated cells. (G) Percentage of indicated populations in BM cells from control and *lncEry* cKO mice and related gating strategies. (H) Representative scan images from triplicate experiments in Figure 4G. (I) BFU-E colony assays of 500 control or *lncEry* cKO BM cells cultured in methylcellulose-based medium with EPO cytokine stimulation for 10-14 days. Scale bar,

100 μm. Each error bar represents the mean ± S.D. for triplicate experiments. *P < 0.05, **P < 0.01.

**Figure S5. *LncEry* deletion impairs erythroid differentiation**

(A) Schematic of *loxP* sequences integrated into the lncEry locus (left). Schematic of conditional knockout (cKO) mice induced with MX1-Cre for 2 months before flow cytometry analysis. (B) qPCR analysis of *lncEry* isoform expression in *flox/flox-2* or Δ/Δ-2 BM cells. (C) Percentage of indicated populations in the BM of *flox/flox-2* or Δ/Δ-2 mice. (D) Absolute numbers or concentrations of indicated items in the peripheral blood of *flox/flox-2* or Δ/Δ-2 mice.

**Figure S6. *LncEry* deletion decrease *Klf1* expression in MEP cells**

(A) Experimental flow-chart for sorting of MEP cells from bone marrow (BM) of *flox/flox* or Δ/Δ mice and RNA-seq, ATAC-seq, and Cut&Tag (left). The number of overlapping downregulated differentially expressed genes (DEGs) and downregulated peak genes in *lncEry*-depleted MEP cells according to RNA-seq and ATAC-seq, respectively (right). (B) Gene enrichment analysis showing −log10 of the uncorrected P-value on the x-axis; darker shading corresponds to a greater number of enriched genes in each term. (C) Determination of low cytometric analysis of ROS levels by DCF-DA (left), mitochondrial membrane potential by TMRE (middle), and glucose uptake 2-NBDG (left) in MEP cells from BM of *flox/flox* or Δ/Δ mice. (D) qPCR confirming that ChIRP retrieved 17% of cellular *lncEry* RNA and 0.1% *Gapdh* RNA. Lac Z probes were used as negative controls. (E-G) TSS profile and heat map showing binding of *lncEry* from ChIRP-seq, H3K4me3 and H3K27ac from Cut&Tag in relation to promoter regions. (H) Number of overlapping genes from Figure 4D and downregulated peak genes from H3K27ac Cut&Tag sequencing of MEP cells. (I) Western blot analysis of the expression of indicated proteins in MEP cells sorted from BM of *flox/flox* or Δ/Δ mice. (H) GSEA enrichment plot of *Klf1* target gene set for DEGs between *flox/flox* and Δ/Δ MEP cells. Error bars represent the mean ± S.D. of triplicate experiments. *P < 0.05, **P < 0.01.

**Figure S7. *LncEry* regulates late stage of erythropoiesis by promoting expression of globin genes**

(A) qPCR analysis of indicated genes in *lncEry*-depleted MEL cells. (B) MEL cells were transfected with control or *lncEry* siRNA, and cellular extracts were prepared and analyzed by western blotting (left). The intensity of each band was quantified by densitometry with Image J software normalized to β-actin (right). (C) Heat map showing binding of H3K4me3 and RNA polymerase II S5p in relation to promoter regions. (D) ChIP-seq trace showing RNA polymerase II S5p and H3K4me3 binding of control or *lncEry*-knockout cells in relation to the indicated gene promoter. (E) GSEA enrichment plot of *Klf1* target gene set for DEGs between control and *lncEry* siRNAs in MEL cells. The data were visualized using IGV software. Each error bar represents the mean ± S.D. for triplicate experiments. *P < 0.05, **P < 0.01.

**Figure S8. *LncEry* is physically associated with Wdr82**

(A) Experimental design for identifying *lncEry*-interacting proteins. *In vitro* transcribed *lncEry*-3 was used as the bait, and anti-*lncEry*-3 was used as the control. (B) Overlapping *lncEry* interaction partners from MEP and MEL cells, as analyzed by mass spectrometry. (C-D) CFU colony assays of 2,000 $cKit^+$ cells transfected with control, Wdr82, or Ddx5 shRNAs cultured 10-14 days in complete methylcellulose-based medium. The colony numbers are provided in D. Representative images from triplicate experiments are shown in C. (E) Cellular extracts of $cKit^+$ cells transfected with control, Wdr82, or Ddx5 shRNAs were prepared and analyzed by western blotting with indicated antibodies. (F) BFU-E colony assays of 2,000 $cKit^+$ cells transfected with control or Wdr82 shRNAs cultured 10-14 days in complete methylcellulose-based medium with EPO cytokine stimulation. The colony numbers were counted (right). Representative images from triplicate experiments are shown (left). (G) Flow analysis percentage of $CD44^+$ in $Ter119^+$ $cKit^+$ cells transfected with control or Wdr82 shRNAs cultured in serum-free expansion medium with growth factors SCF, IL-3, and EPO for 7-10 days. Flow analysis of CD44 expression in $Ter119^+$ cells. (H) MEL cells with doxycycline-inducible expression of stably integrated Flag-Wdr82 were collected. Cellular extracts were prepared and subjected to affinity purification using an anti-FLAG affinity column. After extensive washing in high salt solution, the purified Wdr82 protein was stained with Coomassie blue. (I) RNAscope and immunostaining assays of MEL cells were performed using *lncEry* probes and Wdr82 antibodies, respectively, followed by confocal microscopy. Scale bar, 10 μm. Each error bar represents the mean ± S.D. for triplicate experiments. *P < 0.05, **P < 0.01.

**Figure S9. *LncEry*-Wdr82 regulates transcriptional activation of *Klf1* and globin genes**

(A) Peak center profile and heat map showing binding of Wdr82 at whole genome of MEP cells. (B) qPCR of *lncEry* expression and western blotting of the expression of indicated proteins in the reporter assays shown in Figure 7B. (C) ChIRP-seq trace showing *lncEry* binding in relation to the indicated gene regions (upper panel). qPCR and agarose gel electrophoresis of chromatin-isolated sequence of the indicated gene regions. (D-E) Western blot of the expression of indicated proteins in the reporter assays shown in Figure 7 E-F and Supplementary Figure 8 E-G. (F-H) MEL cells were co-transfected with the indicated siRNAs or plasmids together with Renilla and the indicated promoter luciferase constructs. Relative luciferase activity was determined by sequential normalization to Renilla and pGL3-vector activity. (I) qPCR of *lncEry* expression in each sample shown in Figure 7G. (J) MEL cells were transfected with control or *lncEry* siRNAs, and the soluble chromatin was immunoprecipitated with antibodies against RbBP5 or Ash2l, followed by qPCR. Relative fold enrichment was determined by sequential normalization to the input and control siRNA samples. Each error bar represents the mean ± S.D. for triplicate experiments. *P < 0.05, **P < 0.01.

**Supplementary Table 1. Top 10 highly expressed lncRNAs in each population**

| rownames.data. | maxJSspe | sampleID | maxValu | LT_HSC | ST_HSC |
|---|---|---|---|---|---|
| ENSMUSG00000085069.2_Gm13111 | 0.64975 | LT_HSC | 1.62821 | 1.62821 | 0.449052 |
| ENSMUSG00000104066.1_Gm37955 | 0.59292 | LT_HSC | 4.14909 | 4.14909 | 0 |
| ENSMUSG00000114076.1_AC121804.2 | 0.568551 | LT_HSC | 1.1259 | 1.1259 | 0.186416 |
| ENSMUSG00000108004.1_Gm44080 | 0.565781 | LT_HSC | 2.25681 | 2.25681 | 0.141609 |
| ENSMUSG00000021268.17_Meg3 | 0.525013 | LT_HSC | 2.04609 | 2.04609 | 0.698963 |
| ENSMUSG00000103870.1_Gm38362 | 0.471166 | LT_HSC | 1.09704 | 1.09704 | 0.707631 |
| ENSMUSG00000000031.16_H19 | 0.429549 | LT_HSC | 1.02003 | 1.02003 | 0.440054 |
| ENSMUSG00000102575.1_Gm37241 | 0.42948 | LT_HSC | 1.20796 | 1.20796 | 0.724824 |
| ENSMUSG00000104486.1_Gm38066 | 0.424116 | LT_HSC | 2.31487 | 2.31487 | 1.76369 |
| ENSMUSG00000102460.1_Gm38197 | 0.412521 | LT_HSC | 3.77752 | 3.77752 | 2.75045 |
| ENSMUSG00000085125.7_Gm16070 | 0.437328 | ST_HSC | 1.46272 | 1.29951 | 1.46272 |
| ENSMUSG00000085645.1_Hoxb5os | 0.420536 | ST_HSC | 1.69558 | 1.26342 | 1.69558 |
| ENSMUSG00000111952.1_Gm4489 | 0.414052 | ST_HSC | 2.18582 | 0.889516 | 2.18582 |
| ENSMUSG00000096971.2_4930556M19Rik | 0.403306 | ST_HSC | 1.78021 | 1.60743 | 1.78021 |
| ENSMUSG00000086534.2_Gm16758 | 0.388685 | ST_HSC | 5.84713 | 2.72297 | 5.84713 |
| ENSMUSG00000102205.1_9430092D12Rik | 0.375443 | ST_HSC | 1.15606 | 0.932777 | 1.15606 |
| ENSMUSG00000105940.1_Gm42635 | 0.368934 | ST_HSC | 1.14942 | 0.883954 | 1.14942 |
| ENSMUSG00000086884.1_Gm16225 | 0.366482 | ST_HSC | 1.4109 | 0.86052 | 1.4109 |
| ENSMUSG00000096870.2_Gm21816 | 0.360337 | ST_HSC | 1.45298 | 1.11185 | 1.45298 |
| ENSMUSG00000085409.1_Gm16155 | 0.355 | ST_HSC | 1.25752 | 0.699758 | 1.25752 |
| ENSMUSG00000103075.1_Gm38109 | 0.552722 | MPP | 9.4372 | 0 | 0 |
| ENSMUSG00000097228.1_Gm26597 | 0.42016 | MPP | 1.37946 | 0.188053 | 0.420969 |
| ENSMUSG00000105296.1_Gm19708 | 0.419666 | MPP | 4.8342 | 0.729595 | 1.54733 |
| ENSMUSG00000087228.1_Gm12827 | 0.405645 | MPP | 1.42774 | 0.430446 | 0.623324 |
| ENSMUSG00000106560.1_Gm43119 | 0.394113 | MPP | 1.73949 | 0.858671 | 0.634088 |
| ENSMUSG00000087273.1_Gm13203 | 0.391927 | MPP | 2.71056 | 0.977897 | 1.84887 |
| ENSMUSG00000073164.11_2410018L13Rik | 0.369074 | MPP | 1.03416 | 0.191874 | 0.595403 |
| ENSMUSG00000085912.1_Trp53cor1 | 0.367225 | MPP | 1.83239 | 1.07072 | 1.39991 |
| ENSMUSG00000108079.1_Gm44210 | 0.359989 | MPP | 4.52399 | 2.71074 | 3.80362 |
| ENSMUSG00000109594.1_1700047O18Rik | 0.344829 | MPP | 1.78167 | 0.937144 | 1.65299 |
| ENSMUSG00000101009.6_1700108F19Rik | 0.837446 | CLP | 1.86679 | 0.053563 | 0 |
| ENSMUSG00000102332.1_Gm19331 | 0.543405 | CLP | 2.5627 | 0 | 0.013979 |
| ENSMUSG00000087038.9_2900079G21Rik | 0.51781 | CLP | 1.40787 | 0.181336 | 0 |
| ENSMUSG00000086150.1_Bach2os | 0.452715 | CLP | 1.20601 | 0.035668 | 0.022067 |
| ENSMUSG00000047462.9_A530099J19Rik | 0.404419 | CLP | 2.1391 | 0.135863 | 0.009009 |
| ENSMUSG00000110958.1_AC153891.1 | 0.392749 | CLP | 3.3706 | 0.508691 | 0.926451 |
| ENSMUSG00000114996.1_AC131586.1 | 0.391074 | CLP | 1.09418 | 0.695741 | 0.242429 |
| ENSMUSG00000109245.1_Gm44860 | 0.389297 | CLP | 4.42636 | 0.202243 | 0.042779 |
| ENSMUSG00000090024.1_Gm16350 | 0.376096 | CLP | 1.18129 | 0.679691 | 0.124826 |
| ENSMUSG00000108897.1_Gm44861 | 0.363991 | CLP | 3.08446 | 0.228891 | 0.073298 |
| ENSMUSG00000099587.1_Gm28967 | 0.873968 | CMP | 10.9805 | 0 | 0 |
| ENSMUSG00000108332.1_D530033B14Rik | 0.532131 | CMP | 3.19102 | 0.043991 | 0.077214 |
| ENSMUSG00000106588.1_Gm17590 | 0.386345 | CMP | 4.49841 | 0.952038 | 0.850317 |
| ENSMUSG00000085550.1_Gm15658 | 0.311013 | CMP | 13.9983 | 6.14635 | 11.9816 |
| ENSMUSG00000043993.6_2900052L18Rik | 0.303741 | CMP | 2.04603 | 2.03435 | 1.7782 |
| ENSMUSG00000052724.7_Gm9888 | 0.284684 | CMP | 1.97406 | 1.11854 | 1.11596 |
| ENSMUSG00000087547.1_Platr27 | 0.277918 | CMP | 1.46932 | 0.875124 | 0.598959 |
| ENSMUSG00000072893.12_4933439C10Rik | 0.259329 | CMP | 3.71047 | 2.67948 | 3.48989 |
| ENSMUSG00000100680.1_1810044D09Rik | 0.255094 | CMP | 6.98564 | 6.57618 | 6.06392 |
| ENSMUSG00000097772.7_5430416N02Rik | 0.254453 | CMP | 41.8308 | 29.4948 | 28.3625 |
| ENSMUSG00000101378.1_Gm28086 | 0.596971 | GMP | 1.37494 | 0.342609 | 0 |
| ENSMUSG00000109770.1_Gm30085 | 0.458174 | GMP | 5.38078 | 0.200877 | 0.102435 |
| ENSMUSG00000100094.1_1810008I18Rik | 0.414616 | GMP | 1.95416 | 0.38611 | 0.191008 |

| | | | | | |
|---|---|---|---|---|---|
| ENSMUSG00000045928.2_4933440M02Rik | 0.407807 | GMP | 4.16104 | 0.488857 | 0.268983 |
| ENSMUSG00000026729.9_4930562F07Rik | 0.372984 | GMP | 1.02913 | 0.062425 | 0.115057 |
| ENSMUSG00000109959.1_Gm45355 | 0.351132 | GMP | 1.40804 | 0.709115 | 0.391696 |
| ENSMUSG00000078891.4_Gm11008 | 0.33245 | GMP | 10.4755 | 0 | 0 |
| ENSMUSG00000097157.2_Gm26512 | 0.316534 | GMP | 2.47303 | 1.50236 | 2.10622 |
| ENSMUSG00000097440.2_Gm6277 | 0.313872 | GMP | 3.95394 | 2.14314 | 2.32225 |
| ENSMUSG00000096986.2_4930509E16Rik | 0.311837 | GMP | 1.57897 | 0.608061 | 0.582164 |
| ENSMUSG00000085245.1_Gm11713 | 0.470545 | MEP | 1.61134 | 0.08252 | 0.072732 |
| ENSMUSG00000085634.1_Gm15290 | 0.466542 | MEP | 6.24858 | 0.754863 | 0 |
| ENSMUSG00000074388.3_Gm5544 | 0.413004 | MEP | 1.11494 | 0.202581 | 0.118794 |
| ENSMUSG00000100037.1_Gm29103 | 0.396827 | MEP | 2.24451 | 0.079178 | 0.288061 |
| ENSMUSG00000086968.8_4933431E20Rik | 0.38756 | MEP | 12.7575 | 2.56932 | 1.55694 |
| ENSMUSG00000086712.2_AI427809 | 0.378333 | MEP | 4.74844 | 1.1538 | 0.13892 |
| ENSMUSG00000085723.2_Gm15915 | 0.372103 | MEP | 45.5968 | 6.91776 | 1.05407 |
| ENSMUSG00000086587.7_Gm11837 | 0.366789 | MEP | 11.887 | 4.22196 | 2.55381 |
| ENSMUSG00000034764.15_1700006J14Rik | 0.335417 | MEP | 30.4967 | 4.25246 | 1.12848 |
| ENSMUSG00000031736.11_Crnde | 0.332892 | MEP | 4.74775 | 1.20563 | 0.839334 |
| ENSMUSG00000085447.1_9530026F06Rik | 1 | NuE | 3.22008 | 0 | 0 |
| ENSMUSG00000091351.1_Gm17135 | 1 | NuE | 1.08577 | 0 | 0 |
| ENSMUSG00000102095.1_C730036E19Rik | 1 | NuE | 4.06459 | 0 | 0 |
| ENSMUSG00000103647.1_Gm37707 | 1 | NuE | 1.04472 | 0 | 0 |
| ENSMUSG00000087330.1_Bloodlinc | 0.911623 | NuE | 37.9171 | 0.061293 | 0 |
| ENSMUSG00000115207.1_AC159006.2 | 0.902806 | NuE | 2.95773 | 0.003458 | 0 |
| ENSMUSG00000114652.1_AC107663.2 | 0.896733 | NuE | 3.15743 | 0 | 0 |
| ENSMUSG00000114235.1_AC117769.1 | 0.895844 | NuE | 4.42 | 0.029659 | 0 |
| ENSMUSG00000085118.2_Gm15774 | 0.889455 | NuE | 9.49099 | 0 | 0.011668 |
| ENSMUSG00000112569.1_Gm33111 | 0.886461 | NuE | 7.20203 | 0.010898 | 0 |
| ENSMUSG00000110017.1_Gm45237 | 0.63022 | MK | 1.27999 | 0.139874 | 0.071111 |
| ENSMUSG00000107552.1_Gm44096 | 0.622757 | MK | 4.14038 | 0 | 0 |
| ENSMUSG00000112261.1_AC167222.1 | 0.575095 | MK | 4.90576 | 0.030766 | 0 |
| ENSMUSG00000087382.7_Ctcflos | 0.57051 | MK | 7.75436 | 0.014827 | 0.025266 |
| ENSMUSG00000102715.1_Gm6209 | 0.542806 | MK | 1.11399 | 0.066815 | 0.013152 |
| ENSMUSG00000114422.1_AC165281.1 | 0.534483 | MK | 1.07887 | 0.05274 | 0.031542 |
| ENSMUSG00000090081.1_Gm16587 | 0.533688 | MK | 5.62272 | 0.019854 | 0.017354 |
| ENSMUSG00000085558.7_4930412C18Rik | 0.510349 | MK | 5.45696 | 0.474515 | 0.229676 |
| ENSMUSG00000097466.8_D430036J16Rik | 0.506873 | MK | 2.13323 | 0.014439 | 0.008191 |
| ENSMUSG00000104068.1_Gm37199 | 0.506831 | MK | 4.16297 | 0.09958 | 0.063361 |
| ENSMUSG00000099599.1_Gm28548 | 0.764305 | GR | 1.37031 | 0 | 0 |
| ENSMUSG00000113960.1_4933412O06Rik | 0.736601 | GR | 1.86073 | 0.013403 | 0 |
| ENSMUSG00000090307.7_1700071M16Rik | 0.712948 | GR | 2.76817 | 0.005543 | 0 |
| ENSMUSG00000112599.1_AC160863.3 | 0.656838 | GR | 2.88606 | 0 | 0 |
| ENSMUSG00000113309.1_AC173210.1 | 0.649602 | GR | 1.92715 | 0.016484 | 0 |
| ENSMUSG00000114980.1_AC102815.1 | 0.644631 | GR | 1.92489 | 0.014716 | 0 |
| ENSMUSG00000112097.1_Gm32515 | 0.635264 | GR | 7.94109 | 0 | 0 |
| ENSMUSG00000084807.1_Gm13073 | 0.627194 | GR | 4.86369 | 0 | 0.04613 |
| ENSMUSG00000095369.1_Gm21859 | 0.62354 | GR | 1.20733 | 0.015446 | 0 |
| ENSMUSG00000097358.1_Gm26773 | 0.608085 | GR | 4.6552 | 0.035198 | 0.004864 |
| ENSMUSG00000087196.1_Gm13373 | 0.555406 | MR | 1.05128 | 0 | 0 |
| ENSMUSG00000085261.1_Gm13814 | 0.545887 | MR | 1.215 | 0.005615 | 0.004844 |
| ENSMUSG00000091985.1_Gm17354 | 0.509354 | MR | 1.68985 | 0.088704 | 0.064397 |
| ENSMUSG00000112360.1_AC153568.1 | 0.473128 | MR | 1.43206 | 0 | 0 |
| ENSMUSG00000093622.1_Gm20703 | 0.468765 | MR | 1.79961 | 0 | 0.09643 |
| ENSMUSG00000111034.1_AC153955.1 | 0.43716 | MR | 1.54326 | 0 | 0 |
| ENSMUSG00000110235.1_Gm5086 | 0.430872 | MR | 2.72389 | 0.01308 | 0 |
| ENSMUSG00000085135.7_Gm13713 | 0.407462 | MR | 2.09062 | 0 | 0.357566 |
| ENSMUSG00000102750.1_Gm37058 | 0.40267 | MR | 2.7901 | 0.050484 | 0.007618 |

| | | | | | |
|---|---|---|---|---|---|
| ENSMUSG00000111535.1_Gm35154 | 0.393749 | MR | 3.66254 | 0.047822 | 0.03024 |
| ENSMUSG00000114884.1_AC168059.2 | 0.493753 | MONO | 2.1703 | 0 | 0.118464 |
| ENSMUSG00000112095.1_A130077B15Rik | 0.389587 | MONO | 1.78458 | 0.063703 | 0.045037 |
| ENSMUSG00000103451.1_Gm33973 | 0.386255 | MONO | 1.43335 | 0.054888 | 0 |
| ENSMUSG00000053749.12_Gm9920 | 0.363518 | MONO | 2.55875 | 0.183645 | 0.698794 |
| ENSMUSG00000110593.1_Gm45822 | 0.360566 | MONO | 1.37072 | 0 | 0 |
| ENSMUSG00000105632.1_Gm43272 | 0.357471 | MONO | 1.27846 | 0.194976 | 0.190794 |
| ENSMUSG00000107355.1_AI839979 | 0.342071 | MONO | 59.4612 | 1.22478 | 0.329361 |
| ENSMUSG00000085360.1_Arhgap27os2 | 0.342044 | MONO | 2.86496 | 0.41093 | 0.359529 |
| ENSMUSG00000097832.1_Gm26912 | 0.309396 | MONO | 1.18828 | 0.555819 | 0.367139 |
| ENSMUSG00000052248.15_Zeb2os | 0.293745 | MONO | 23.146 | 2.39573 | 2.74877 |
| ENSMUSG00000086517.1_Gm12534 | 1 | CD4_T | 2.08539 | 0 | 0 |
| ENSMUSG00000084860.1_2610204G07Rik | 0.875504 | CD4_T | 1.94105 | 0 | 0 |
| ENSMUSG00000086255.1_Gm11534 | 0.84947 | CD4_T | 2.56993 | 0 | 0 |
| ENSMUSG00000085808.1_Gm14718 | 0.790587 | CD4_T | 2.27331 | 0 | 0 |
| ENSMUSG00000111546.1_AC117245.3 | 0.745465 | CD4_T | 3.75442 | 0.006746 | 0 |
| ENSMUSG00000084993.1_Gm12305 | 0.740844 | CD4_T | 2.49981 | 0.16952 | 0 |
| ENSMUSG00000110766.1_AC154295.1 | 0.732597 | CD4_T | 1.18924 | 0 | 0 |
| ENSMUSG00000109827.1_Gm6213 | 0.68294 | CD4_T | 1.123 | 0 | 0 |
| ENSMUSG00000106275.1_Gm42495 | 0.670406 | CD4_T | 1.0139 | 0.082601 | 0 |
| ENSMUSG00000110803.1_Gm20275 | 0.641758 | CD4_T | 2.6793 | 0 | 0 |
| ENSMUSG00000114390.1_AC124680.1 | 0.873339 | CD8_T | 6.77646 | 0 | 0 |
| ENSMUSG00000107266.1_Gm43698 | 0.834298 | CD8_T | 42.1512 | 0 | 0 |
| ENSMUSG00000050179.3_A930002I21Rik | 0.707359 | CD8_T | 10.5809 | 0.01054 | 0.010111 |
| ENSMUSG00000098108.7_Gm27008 | 0.663133 | CD8_T | 2.04468 | 0 | 0 |
| ENSMUSG00000104417.1_Gm37068 | 0.634878 | CD8_T | 1.06953 | 0 | 0 |
| ENSMUSG00000112417.1_A430028G04Rik | 0.633711 | CD8_T | 2.44731 | 0.00914 | 0 |
| ENSMUSG00000110656.1_Gm20735 | 0.633439 | CD8_T | 2.59022 | 0 | 0 |
| ENSMUSG00000108132.1_Gm44175 | 0.624126 | CD8_T | 1.16318 | 0.013438 | 0.007623 |
| ENSMUSG00000102280.1_Gm36999 | 0.573073 | CD8_T | 5.82217 | 0 | 0 |
| ENSMUSG00000112230.1_Ifngas1 | 0.555213 | CD8_T | 4.92686 | 0 | 0 |
| ENSMUSG00000108232.1_Gm43916 | 0.801785 | B | 1.31523 | 0.024846 | 0.023339 |
| ENSMUSG00000087089.1_Gm12160 | 0.735372 | B | 1.81794 | 0 | 0 |
| ENSMUSG00000102785.5_Gm2447 | 0.695574 | B | 1.19324 | 0 | 0 |
| ENSMUSG00000114306.1_AC144938.1 | 0.691087 | B | 1.30193 | 0 | 0 |
| ENSMUSG00000104927.1_Gm43388 | 0.687518 | B | 11.6595 | 0.082856 | 0 |
| ENSMUSG00000102882.1_Gm2065 | 0.684498 | B | 1.12657 | 0.029212 | 0 |
| ENSMUSG00000085844.1_Gm11690 | 0.682975 | B | 3.4988 | 0 | 0 |
| ENSMUSG00000092591.1_Gm20429 | 0.678756 | B | 5.12298 | 0 | 0.038529 |
| ENSMUSG00000084773.1_Gm12159 | 0.663414 | B | 1.29486 | 0.011543 | 0 |
| ENSMUSG00000111839.1_Gm39383 | 0.654847 | B | 5.5244 | 0.108092 | 0.016814 |
| ENSMUSG00000104077.1_Gm37027 | 1 | NK | 1.50321 | 0 | 0 |
| ENSMUSG00000101264.1_Gm28347 | 0.829187 | NK | 3.50844 | 0.143359 | 0 |
| ENSMUSG00000105247.1_Gm42519 | 0.768068 | NK | 2.40062 | 0.048001 | 0.04666 |
| ENSMUSG00000104781.1_Gm43303 | 0.730094 | NK | 1.68304 | 0 | 0.039999 |
| ENSMUSG00000089755.1_0610012D04Rik | 0.713558 | NK | 1.37401 | 0 | 0.04146 |
| ENSMUSG00000087132.7_A930001C03Rik | 0.704067 | NK | 1.22434 | 0 | 0 |
| ENSMUSG00000105963.1_Gm43647 | 0.690537 | NK | 1.62252 | 0 | 0 |
| ENSMUSG00000098292.1_Gm27194 | 0.655956 | NK | 12.5518 | 0.07738 | 0.023464 |
| ENSMUSG00000099343.1_Gm28836 | 0.649958 | NK | 2.9958 | 0 | 0 |
| ENSMUSG00000104340.1_Gm10522 | 0.647861 | NK | 10.3623 | 0.018277 | 0 |
| ENSMUSG00000101264.1_Gm28347 | 0.829187 | NK | 3.50844 | 0.143359 | 0 |
| ENSMUSG00000105247.1_Gm42519 | 0.768068 | NK | 2.40062 | 0.048001 | 0.04666 |
| ENSMUSG00000089755.1_0610012D04Rik | 0.713558 | NK | 1.37401 | 0 | 0.04146 |
| ENSMUSG00000104781.1_Gm43303 | 0.712246 | NK | 1.68304 | 0 | 0.039999 |
| ENSMUSG00000087132.7_A930001C03Rik | 0.700003 | NK | 1.22434 | 0 | 0 |

| | | | | | |
|---|---|---|---|---|---|
| ENSMUSG00000105963.1_Gm43647 | 0.690537 | NK | 1.62252 | 0 | 0 |
| ENSMUSG00000098292.1_Gm27194 | 0.648659 | NK | 12.5518 | 0.07738 | 0.023464 |
| ENSMUSG00000099343.1_Gm28836 | 0.640606 | NK | 2.9958 | 0 | 0 |
| ENSMUSG00000104340.1_Gm10522 | 0.640601 | NK | 10.3623 | 0.018277 | 0 |

| MPP | CLP | CMP | GMP | MEP | NuE | MK | GR | MR |
|---|---|---|---|---|---|---|---|---|
| 0.033276 | 0 | 0 | 0 | 0 | 0.005874 | 0.008608 | 0 | 0.02065 |
| 0 | 2.03299 | 0 | 0 | 0 | 0 | 0 | 0 | 0 |
| 0.540502 | 0 | 0 | 0 | 0 | 0 | 0 | 0 | 0 |
| 0.124163 | 0.550327 | 0.064549 | 0 | 0 | 0.0146 | 0.035659 | 0.087086 | 0.026717 |
| 0.420022 | 0.060573 | 0.001611 | 0.002613 | 0 | 0.001482 | 0.040569 | 0.026139 | 0.079889 |
| 0.462978 | 0.074732 | 0.029828 | 0.037753 | 0 | 0 | 0.058744 | 0 | 0.040494 |
| 0.909328 | 0.128875 | 0.212566 | 0 | 0.014206 | 0.023685 | 0.035422 | 0.024859 | 0.019825 |
| 0.594981 | 0.043387 | 0.338151 | 0.144792 | 0 | 0 | 0.053683 | 0 | 0.135907 |
| 1.39577 | 0.141865 | 0.192225 | 0.050544 | 0 | 0.019599 | 0.19054 | 0 | 0.188987 |
| 2.32794 | 0.235311 | 0.414009 | 0.15286 | 0.00702 | 0.015106 | 0.159916 | 0.011387 | 0.348545 |
| 0.880142 | 0.030076 | 0.084766 | 0.004491 | 0 | 0.001859 | 0.060175 | 0 | 0.154737 |
| 0.965609 | 0.023715 | 0.433179 | 0.081524 | 0 | 0 | 0.027587 | 0 | 0.234737 |
| 0.67035 | 0.076401 | 1.04405 | 0 | 0 | 0 | 0.532125 | 0 | 0.231466 |
| 0.811958 | 0.076814 | 0.485024 | 0.165344 | 0.029608 | 0 | 0.219996 | 0 | 0.139706 |
| 2.27356 | 0.171249 | 2.17983 | 0.33091 | 0.034637 | 0.0472 | 0.755359 | 0.04768 | 1.07034 |
| 1.01648 | 0.221184 | 0.199331 | 0.077334 | 0 | 0.066964 | 0.323267 | 0 | 0.213562 |
| 0.139662 | 0.096707 | 0.791731 | 0.094011 | 0.351504 | 0.014815 | 0.327961 | 0.133759 | 0.250177 |
| 1.08265 | 0.534954 | 0.526076 | 0.240116 | 0 | 0 | 0.260619 | 0 | 0 |
| 1.34289 | 0.425706 | 0.249757 | 0.129794 | 0 | 0.042551 | 0.560369 | 0.031112 | 0.302256 |
| 1.15274 | 0.219773 | 0.774436 | 0.12433 | 0.15275 | 0 | 0.392702 | 0 | 0.356781 |
| 9.4372 | 0 | 0 | 0 | 0 | 0 | 7.06541 | 0 | 0 |
| 1.37946 | 1.11196 | 0.333099 | 0.060277 | 0.009514 | 0.036111 | 0.077201 | 0.020397 | 0.118234 |
| 4.8342 | 1.89021 | 0.63654 | 0.379291 | 0 | 0.072193 | 0.115015 | 0 | 0.280639 |
| 1.42774 | 0.37523 | 0.581287 | 0.075082 | 0.020209 | 0 | 0.127379 | 0 | 0.158635 |
| 1.73949 | 1.57263 | 0.357083 | 0.018585 | 0.067877 | 0.043888 | 0.033445 | 0.014034 | 0.061602 |
| 2.71056 | 0.426511 | 0.428364 | 0.31754 | 0 | 0.02836 | 0.217758 | 0 | 0.159391 |
| 1.03416 | 0.436115 | 0.16147 | 0.070643 | 0.029898 | 0.016292 | 0.129177 | 0 | 0.124856 |
| 1.83239 | 0.753868 | 0.426718 | 0.074266 | 0 | 0.059052 | 0.198822 | 0.091784 | 0.346923 |
| 4.52399 | 1.54692 | 1.17494 | 0.667727 | 0.031839 | 0.255935 | 0.748742 | 0 | 0.338367 |
| 1.78167 | 1.17099 | 0.440796 | 0.2421 | 0.014897 | 0.117781 | 0.223806 | 0.028331 | 0.044942 |
| 0.032768 | 1.86679 | 0 | 0 | 0 | 0 | 0 | 0 | 0 |
| 1.24884 | 2.5627 | 0.056114 | 0 | 0 | 0 | 0.072322 | 0.166134 | 0 |
| 0.397183 | 1.40787 | 0.071288 | 0 | 0 | 0.01306 | 0.270462 | 0.032828 | 0.021745 |
| 0.023497 | 1.20601 | 0.006566 | 0.00711 | 0 | 0.961486 | 0.249445 | 0.025407 | 0.060937 |
| 0.177956 | 2.1391 | 0.00923 | 0.08632 | 0.005261 | 0.095789 | 0.091866 | 0.767769 | 1.09512 |
| 2.27825 | 3.3706 | 0.832635 | 0.222694 | 0.022479 | 0.031586 | 0.355273 | 0.081668 | 0.468373 |
| 0.301564 | 1.09418 | 0.214315 | 0.025856 | 0.040727 | 0.074894 | 0.290283 | 0.013879 | 0.027334 |
| 0.061673 | 4.42636 | 0.016129 | 0.021428 | 0.006047 | 0.874704 | 2.16771 | 0.512017 | 2.47071 |
| 0.12822 | 1.18129 | 0 | 0.199043 | 0 | 0.089646 | 0.177618 | 0 | 0 |
| 0.124028 | 3.08446 | 0.055621 | 0.038101 | 0 | 1.1131 | 1.71838 | 0.876414 | 2.21487 |
| 0 | 0 | 10.9805 | 0 | 0 | 0 | 0 | 0.124428 | 0 |
| 0.039294 | 0.061196 | 3.19102 | 0.03031 | 0 | 0 | 0.144484 | 0.153129 | 0.144698 |
| 0.128132 | 0.807714 | 4.49841 | 0.731027 | 0.313508 | 0.165288 | 1.17378 | 0.136501 | 1.99378 |
| 7.55787 | 2.17167 | 13.9983 | 6.31512 | 3.11855 | 0.310951 | 3.97416 | 0.246084 | 4.60455 |
| 1.68162 | 0.862755 | 2.04603 | 0.235489 | 0.612333 | 0.2469 | 1.08382 | 0.641613 | 0.581527 |
| 1.45734 | 0.694332 | 1.97406 | 0.911947 | 1.18795 | 0.228987 | 1.01204 | 0.407216 | 0.738625 |
| 1.03218 | 0.90606 | 1.46932 | 1.00468 | 0.857364 | 0.144023 | 0.528504 | 0.104766 | 0.410838 |
| 3.45351 | 2.64191 | 3.71047 | 3.04166 | 2.19179 | 2.06463 | 2.5769 | 0.702486 | 1.55458 |
| 6.7626 | 4.59139 | 6.98564 | 5.60366 | 4.47185 | 2.64982 | 4.64874 | 3.84035 | 4.83832 |
| 29.9394 | 34.2125 | 41.8308 | 33.8375 | 20.8971 | 9.05598 | 14.7631 | 3.77099 | 11.2287 |
| 0 | 0 | 0 | 1.37494 | 0 | 0 | 0 | 0 | 0 |
| 0.032867 | 0.143737 | 3.14686 | 5.38078 | 0.120705 | 0.044406 | 0.407509 | 0.42964 | 0.437002 |
| 0 | 0.553178 | 0.021189 | 1.95416 | 0 | 0.139559 | 0.723056 | 0.597956 | 0.301131 |

| | | | | | | | | |
|---|---|---|---|---|---|---|---|---|
| 0.035236 | 1.10893 | 0.042295 | 4.16104 | 0.014394 | 0.163097 | 0.584024 | 0.280465 | 0.169831 |
| 0.172148 | 0.335235 | 0.363991 | 1.02913 | 0 | 0.089276 | 0.318295 | 0.157715 | 0.209588 |
| 0.398635 | 0.30976 | 0 | 1.40804 | 0.159579 | 0.590224 | 0.571148 | 0.454321 | 0.755924 |
| 0 | 5.80486 | 6.90785 | 10.4755 | 5.26273 | 0 | 2.81414 | 0 | 0 |
| 1.46808 | 0.73665 | 0.893848 | 2.47303 | 0.077678 | 0.271659 | 1.34324 | 0.603013 | 0.684935 |
| 2.07303 | 2.1328 | 0.934508 | 3.95394 | 0.259813 | 0.299104 | 1.92366 | 0.336331 | 1.09884 |
| 0.509152 | 0.524149 | 0.75945 | 1.57897 | 0.931214 | 0.385672 | 0.745446 | 0.329539 | 0.442853 |
| 0.039756 | 0.033102 | 0.711192 | 0.027371 | 1.61134 | 0.074546 | 0.037511 | 0.078244 | 0.25621 |
| 0.041888 | 0.28057 | 0.034433 | 0.033419 | 6.24858 | 6.02372 | 0.17869 | 0 | 0.160568 |
| 0.061712 | 0.099775 | 0.580448 | 0.053073 | 1.11494 | 0.045651 | 0.100338 | 0.091219 | 0.143523 |
| 0.047382 | 0.520062 | 2.06075 | 0 | 2.24451 | 0.171763 | 0.541774 | 0.070067 | 0.488066 |
| 0.515594 | 0.74717 | 2.65298 | 0.317998 | 12.7575 | 2.79282 | 0.701026 | 0.225762 | 0.909724 |
| 0.138443 | 0.787463 | 0.859491 | 0.107767 | 4.74844 | 2.45191 | 0.547909 | 0.141622 | 0.360759 |
| 0.637153 | 1.28226 | 15.7932 | 0.595862 | 45.5968 | 6.24064 | 1.70716 | 0.502636 | 2.74511 |
| 1.38531 | 1.32641 | 0.856156 | 0.162575 | 11.887 | 7.00864 | 0.330282 | 0.335933 | 0.595925 |
| 0.284458 | 3.15306 | 9.7165 | 1.17526 | 30.4967 | 4.46373 | 5.34313 | 4.28064 | 5.33525 |
| 1.15903 | 1.2029 | 1.64417 | 1.48783 | 4.74775 | 0.952839 | 0.661039 | 0.385432 | 0.963433 |
| 0 | 0 | 0 | 0 | 0 | 3.22008 | 0 | 0 | 0 |
| 0 | 0 | 0 | 0 | 0 | 1.08577 | 0 | 0 | 0 |
| 0 | 0 | 0 | 0 | 0 | 4.06459 | 0 | 0 | 0 |
| 0 | 0 | 0 | 0 | 0 | 1.04472 | 0 | 0 | 0 |
| 0 | 0 | 0 | 0 | 0.024625 | 37.9171 | 0 | 0 | 0 |
| 0 | 0.01425 | 0.002462 | 0.007614 | 0 | 2.95773 | 0 | 0 | 0.003485 |
| 0 | 0 | 0.007969 | 0 | 0 | 3.15743 | 0.011037 | 0.014281 | 0.01177 |
| 0 | 0 | 0.011523 | 0 | 0 | 4.42 | 0 | 0 | 0 |
| 0 | 0 | 0 | 0 | 0.005943 | 9.49099 | 0.015973 | 0.044931 | 0.007637 |
| 0 | 0.009165 | 0 | 0.002678 | 0.012075 | 7.20203 | 0.034462 | 0 | 0.004311 |
| 0.009517 | 0.008452 | 0.038713 | 0 | 0 | 0.047068 | 1.27999 | 0.075707 | 0.006354 |
| 0 | 0.044811 | 0 | 0.043707 | 0.099577 | 0.27131 | 4.14038 | 0.411433 | 0.14443 |
| 0 | 0.450122 | 0 | 0.020739 | 0 | 0.125276 | 4.90576 | 0.228026 | 0.13314 |
| 0 | 0.031123 | 0 | 0.00361 | 0 | 1.61028 | 7.75436 | 0.583284 | 0.137519 |
| 0.007991 | 0.03268 | 0.004917 | 0 | 0 | 0.057761 | 1.11399 | 0.439522 | 0.13749 |
| 0 | 0.156305 | 0.01455 | 0 | 0.015919 | 0.110834 | 1.07887 | 0 | 0.021014 |
| 0 | 0.033179 | 0 | 0.047458 | 0 | 0.252579 | 5.62272 | 1.4289 | 0.860208 |
| 0.102032 | 0.12928 | 0.061403 | 0.107604 | 0.108078 | 0.244432 | 5.45696 | 0.434409 | 0.255595 |
| 0.041707 | 0.864091 | 0.009782 | 0 | 0 | 0.163998 | 2.13323 | 0.09354 | 0.210675 |
| 0.026178 | 0.032887 | 0.003416 | 0.009885 | 0.003856 | 0.245432 | 4.16297 | 1.99739 | 0.15317 |
| 0 | 0 | 0 | 0 | 0 | 0.060095 | 0 | 1.37031 | 0.100295 |
| 0 | 0 | 0 | 0 | 0 | 0.092551 | 0.033957 | 1.86073 | 0.097829 |
| 0 | 0 | 0 | 0 | 0 | 0.124718 | 0.090781 | 2.76817 | 0.146669 |
| 0 | 0 | 0 | 0 | 0 | 0.067933 | 0.328765 | 2.88606 | 0.217698 |
| 0 | 0 | 0 | 0 | 0 | 0.164692 | 0.175509 | 1.92715 | 0.170471 |
| 0 | 0 | 0 | 0 | 0 | 0.128334 | 0.269266 | 1.92489 | 0.071464 |
| 0 | 0 | 0 | 0 | 0 | 0.845573 | 0.129635 | 7.94109 | 0.190185 |
| 0 | 0.085693 | 0.041311 | 0 | 0 | 0.068291 | 0.582306 | 4.86369 | 0.133365 |
| 0 | 0.013759 | 0 | 0.013647 | 0 | 0.046223 | 0.14916 | 1.20733 | 0.089506 |
| 0 | 0.007642 | 0 | 0.016612 | 0 | 0.282998 | 0.311744 | 4.6552 | 0.502958 |
| 0 | 0.027212 | 0 | 0 | 0 | 0.275926 | 0.028416 | 0.088868 | 1.05128 |
| 0 | 0.053954 | 0 | 0.029965 | 0 | 0.127922 | 0.105024 | 0.276704 | 1.215 |
| 0.032859 | 0.056008 | 0.003966 | 0 | 0 | 0.302983 | 0.140427 | 0.213953 | 1.68985 |
| 0 | 0.105054 | 0 | 0 | 0.118748 | 0.702028 | 0.193796 | 0 | 1.43206 |
| 0 | 0.084281 | 0 | 0.068676 | 0 | 0.300762 | 0.085688 | 0.164086 | 1.79961 |
| 0 | 0.146682 | 0 | 0 | 0 | 0.375374 | 0.354357 | 0.134175 | 1.54326 |
| 0.44265 | 1.21812 | 0.02434 | 0 | 0 | 0.849107 | 0.466 | 0.078999 | 2.72389 |
| 0 | 0 | 0 | 0 | 0 | 1.05625 | 0 | 0 | 2.09062 |
| 0.017225 | 0.567837 | 0.02348 | 0.008094 | 0 | 0.496595 | 0.773324 | 0.604505 | 2.7901 |

| | | | | | | | | |
|---|---|---|---|---|---|---|---|---|
| 0 | 0.776789 | 0.012983 | 0.031067 | 0.014913 | 2.05048 | 0.622614 | 0.570856 | 3.66254 |
| 0 | 0.497412 | 0 | 0.192122 | 0 | 0.188193 | 0.549337 | 0.260574 | 0 |
| 0.029788 | 0.610525 | 0.093094 | 0.323771 | 0 | 0.485202 | 0.388513 | 0.336836 | 1.03879 |
| 0.187393 | 0.642295 | 0.040645 | 0.038673 | 0 | 0.594092 | 0.502253 | 0.092307 | 1.18048 |
| 0.408163 | 0.382391 | 0.464307 | 0.481753 | 0.257093 | 0.620645 | 0.398604 | 0.236111 | 1.0808 |
| 0 | 0.393108 | 0.394191 | 0.333677 | 0 | 0.876489 | 0 | 0 | 0 |
| 0.426581 | 0.959587 | 0.159398 | 0.344694 | 0.009653 | 0.254705 | 0.307368 | 0.13768 | 0.669389 |
| 0.57954 | 3.45243 | 0.813545 | 2.47861 | 0.049886 | 11.9682 | 6.19715 | 40.9826 | 42.2716 |
| 0.402633 | 0.589939 | 0.205975 | 0.164205 | 0 | 0.79829 | 0.674966 | 0.386052 | 0.649497 |
| 0.401747 | 0.503303 | 0.334577 | 0.898485 | 0.260257 | 0.482712 | 0.605205 | 0.593966 | 0.400366 |
| 3.11327 | 5.21202 | 5.51419 | 6.74801 | 2.84237 | 5.55235 | 5.8132 | 5.62916 | 10.6558 |
| 0 | 0 | 0 | 0 | 0 | 0 | 0 | 0 | 0 |
| 0 | 0 | 0 | 0 | 0 | 0.028908 | 0 | 0 | 0 |
| 0.045411 | 0 | 0 | 0 | 0.043354 | 0 | 0 | 0 | 0 |
| 0 | 0 | 0 | 0 | 0 | 0 | 0 | 0 | 0 |
| 0 | 0.092345 | 0 | 0 | 0.005849 | 0.004852 | 0.008607 | 0 | 0.018087 |
| 0.067991 | 0.055681 | 0 | 0 | 0 | 0 | 0 | 0 | 0 |
| 0 | 0 | 0 | 0 | 0 | 0.048776 | 0 | 0 | 0 |
| 0 | 0.06267 | 0 | 0 | 0 | 0 | 0 | 0 | 0 |
| 0 | 0.064137 | 0 | 0 | 0 | 0 | 0 | 0 | 0 |
| 0 | 0 | 0 | 0 | 0 | 0 | 0 | 0 | 0 |
| 0 | 0 | 0 | 0 | 0 | 0 | 0 | 0 | 0 |
| 0 | 0.036316 | 0 | 0 | 0 | 0.069319 | 0 | 0 | 0 |
| 0 | 0.015293 | 0 | 0.007627 | 0 | 0.033027 | 0 | 0 | 0.011887 |
| 0 | 0.099652 | 0 | 0 | 0 | 0 | 0.01279 | 0 | 0 |
| 0 | 0 | 0 | 0 | 0 | 0 | 0 | 0 | 0 |
| 0 | 0.013866 | 0 | 0 | 0 | 0.069252 | 0.069121 | 0 | 0.009924 |
| 0 | 0.012091 | 0 | 0 | 0 | 0 | 0 | 0 | 0 |
| 0.005598 | 0.012547 | 0.007303 | 0.007568 | 0.016515 | 0.133231 | 0.026304 | 0.051515 | 0.014964 |
| 0 | 0 | 0 | 0 | 0 | 0.011051 | 0.019364 | 0 | 0 |
| 0 | 0.010768 | 0 | 0 | 0 | 0.04523 | 0 | 0 | 0 |
| 0 | 0 | 0 | 0 | 0 | 0 | 0.020564 | 0 | 0 |
| 0 | 0 | 0 | 0 | 0 | 0.27104 | 0 | 0 | 0 |
| 0 | 0 | 0 | 0 | 0 | 0.005422 | 0.111973 | 0 | 0 |
| 0.00668 | 0 | 0 | 0 | 0 | 0.114712 | 0.104694 | 0 | 0.008369 |
| 0 | 0.061296 | 0 | 0 | 0 | 0.188411 | 0.189303 | 0 | 0 |
| 0.003794 | 0.035448 | 0 | 0 | 0 | 0.018358 | 0.016563 | 0 | 0 |
| 0.00561 | 0.017563 | 0 | 0 | 0 | 0.184889 | 0.105428 | 0 | 0.008448 |
| 0.021527 | 0.043471 | 0 | 0.02543 | 0 | 0.071299 | 0.159399 | 0 | 0.090566 |
| 0.010147 | 0.008439 | 0 | 0 | 0 | 0.140431 | 0.060777 | 0.037698 | 0.013105 |
| 0 | 0.031368 | 0 | 0 | 0 | 0.121159 | 0.117936 | 0 | 0 |
| 0 | 0 | 0 | 0 | 0 | 0 | 0 | 0 | 0 |
| 0 | 0 | 0 | 0 | 0 | 0 | 0 | 0 | 0 |
| 0.007632 | 0.012389 | 0.013206 | 0.006336 | 0.007405 | 0 | 0.008375 | 0 | 0.009179 |
| 0.018661 | 0 | 0.017246 | 0.015908 | 0 | 0.043174 | 0 | 0 | 0.055672 |
| 0 | 0 | 0.040717 | 0 | 0 | 0 | 0.093726 | 0 | 0 |
| 0.03483 | 0.033426 | 0.009953 | 0 | 0 | 0.007389 | 0 | 0 | 0.028074 |
| 0 | 0 | 0 | 0 | 0 | 0 | 0 | 0 | 0.074129 |
| 0.049482 | 0.031123 | 0.035089 | 0.031315 | 0.009857 | 0.063774 | 0.052865 | 0 | 0.050501 |
| 0 | 0 | 0 | 0 | 0 | 0.043151 | 0.17947 | 0.037614 | 0 |
| 0.015608 | 0.038675 | 0 | 0.026537 | 0 | 0.071205 | 0.094094 | 0.030504 | 0.074954 |
| 0 | 0 | 0 | 0 | 0 | 0 | 0 | 0 | 0 |
| 0.007632 | 0.012389 | 0.013206 | 0.007405 | 0.006336 | 0.008375 | 0 | 0 | 0 |
| 0 | 0 | 0.040717 | 0 | 0 | 0.093726 | 0 | 0 | 0 |
| 0.018661 | 0 | 0.017246 | 0 | 0.015908 | 0 | 0 | 0.083924 | 0.0207 |
| 0.03483 | 0.033426 | 0.009953 | 0 | 0 | 0 | 0.015379 | 0 | 0 |

| | | | | | | | | |
|---|---|---|---|---|---|---|---|---|
| 0 | 0 | 0 | 0 | 0 | 0 | 0 | 0 | 0 |
| 0.049482 | 0.031123 | 0.035089 | 0.009857 | 0.031315 | 0.052865 | 0.131615 | 0 | 0.019513 |
| 0 | 0 | 0 | 0 | 0 | 0.17947 | 0 | 0.089952 | 0 |
| 0.015608 | 0.038675 | 0 | 0 | 0.026537 | 0.094094 | 0.020539 | 0.114772 | 0.029351 |

| MONO | CD4_T | CD8_T | B | NK | lnc | geneid |
|---|---|---|---|---|---|---|
| 0 | 0 | 0 | 0 | 0 | 1 | ENSMUSG00000085069.2 |
| 0 | 0 | 0 | 0 | 0 | 1 | ENSMUSG00000104066.1 |
| 0 | 0 | 0 | 0 | 0 | 1 | ENSMUSG00000114076.1 |
| 0.006355 | 0 | 0 | 0 | 0.047153 | 1 | ENSMUSG00000108004.1 |
| 0 | 0.094508 | 0.017106 | 0 | 0.001361 | 1 | ENSMUSG00000021268.17 |
| 0 | 0 | 0 | 0 | 0 | 1 | ENSMUSG00000103870.1 |
| 0 | 0 | 0.013791 | 0 | 0.018154 | 1 | ENSMUSG00000000031.16 |
| 0 | 0 | 0 | 0 | 0 | 1 | ENSMUSG00000102575.1 |
| 0 | 0 | 0 | 0 | 0 | 1 | ENSMUSG00000104486.1 |
| 0.006896 | 0 | 0 | 0 | 0.013453 | 1 | ENSMUSG00000102460.1 |
| 0 | 0 | 0 | 0 | 0 | 1 | ENSMUSG00000085125.7 |
| 0.030568 | 0 | 0 | 0 | 0 | 1 | ENSMUSG00000085645.1 |
| 0 | 0 | 0 | 0 | 0.167477 | 1 | ENSMUSG00000111952.1 |
| 0.073619 | 0 | 0 | 0 | 0.009844 | 1 | ENSMUSG00000096971.2 |
| 0 | 0 | 0.005531 | 0 | 0 | 1 | ENSMUSG00000086534.2 |
| 0.109719 | 0 | 0 | 0 | 0 | 1 | ENSMUSG00000102205.1 |
| 0.012834 | 0 | 0.025307 | 0 | 0.074662 | 1 | ENSMUSG00000105940.1 |
| 0 | 0 | 0.470754 | 0 | 0 | 1 | ENSMUSG00000086884.1 |
| 0.198539 | 0 | 0 | 0.006787 | 0 | 1 | ENSMUSG00000096870.2 |
| 0.153977 | 0 | 0 | 0 | 0 | 1 | ENSMUSG00000085409.1 |
| 0 | 0 | 0 | 0 | 0 | 1 | ENSMUSG00000103075.1 |
| 0.039914 | 0 | 0 | 0.041868 | 0 | 1 | ENSMUSG00000097228.1 |
| 0.215284 | 0 | 0 | 0 | 0 | 1 | ENSMUSG00000105296.1 |
| 0.282023 | 0 | 0 | 0 | 0 | 1 | ENSMUSG00000087228.1 |
| 0.006876 | 0.049614 | 0.073971 | 0 | 0.050472 | 1 | ENSMUSG00000106560.1 |
| 0 | 0 | 0 | 0.131886 | 0.575785 | 1 | ENSMUSG00000087273.1 |
| 0.007817 | 0.103183 | 0.599655 | 0.099908 | 0.246238 | 1 | ENSMUSG00000073164.11 |
| 0.077285 | 0.036029 | 0 | 0.007193 | 0.269803 | 1 | ENSMUSG00000085912.1 |
| 0.029922 | 0 | 0 | 0 | 0 | 1 | ENSMUSG00000108079.1 |
| 0.302355 | 0.070345 | 0.263815 | 0.268932 | 0.088662 | 1 | ENSMUSG00000109594.1 |
| 0 | 0 | 0 | 0 | 0 | 1 | ENSMUSG00000101009.6 |
| 0.013462 | 0.069546 | 0.021932 | 0 | 0 | 1 | ENSMUSG00000102332.1 |
| 0 | 0.028385 | 0.058591 | 0.030367 | 0 | 1 | ENSMUSG00000087038.9 |
| 0.014771 | 0.152993 | 0.09215 | 0.059809 | 0.023944 | 1 | ENSMUSG00000086150.1 |
| 0.034669 | 0.200873 | 0.44389 | 0.328064 | 0.157518 | 1 | ENSMUSG00000047462.9 |
| 0 | 0.038162 | 0 | 0 | 0.198087 | 1 | ENSMUSG00000110958.1 |
| 0.033176 | 0.026362 | 0.070395 | 0.060131 | 0.298169 | 1 | ENSMUSG00000114996.1 |
| 0.064633 | 0.344039 | 0.135985 | 0.259414 | 0.416091 | 1 | ENSMUSG00000109245.1 |
| 0 | 0.856137 | 0.201592 | 0.263146 | 0.295919 | 1 | ENSMUSG00000090024.1 |
| 0.12983 | 0.251465 | 0.190273 | 0.238251 | 0.22138 | 1 | ENSMUSG00000108897.1 |
| 0 | 0 | 0 | 0 | 0 | 1 | ENSMUSG00000099587.1 |
| 0.078125 | 0.343165 | 0.24059 | 0.163097 | 0.07458 | 1 | ENSMUSG00000108332.1 |
| 0 | 0 | 0 | 0 | 0.015422 | 1 | ENSMUSG00000106588.1 |
| 1.01713 | 0.053057 | 0 | 0.053573 | 0 | 1 | ENSMUSG00000085550.1 |
| 0.179401 | 0.051832 | 0 | 0 | 0.666069 | 1 | ENSMUSG00000043993.6 |
| 1.43689 | 0.43541 | 0.567552 | 0.48945 | 0.493896 | 1 | ENSMUSG00000052724.7 |
| 0.939939 | 0.516831 | 1.464 | 1.22166 | 0.329437 | 1 | ENSMUSG00000087547.1 |
| 1.5595 | 2.86557 | 2.0793 | 2.15353 | 1.45771 | 1 | ENSMUSG00000072893.12 |
| 4.39792 | 2.83392 | 4.45271 | 3.58205 | 1.93892 | 1 | ENSMUSG00000100680.1 |
| 20.2564 | 21.3752 | 28.9683 | 17.0533 | 40.1039 | 1 | ENSMUSG00000097772.7 |
| 0.316453 | 0 | 0 | 0 | 0 | 1 | ENSMUSG00000101378.1 |
| 0.070546 | 0 | 0 | 0 | 0 | 1 | ENSMUSG00000109770.1 |
| 0.166149 | 0 | 0 | 0 | 0.016373 | 1 | ENSMUSG00000100094.1 |

| | | | | | | |
|---|---|---|---|---|---|---|
| 0.79323 | 0.034786 | 1.00313 | 0 | 0.287432 | 1 | ENSMUSG00000045928.2 |
| 0.735969 | 0.035693 | 0.033074 | 0.068934 | 0 | 1 | ENSMUSG00000026729.9 |
| 0 | 0 | 0 | 0 | 0 | 1 | ENSMUSG00000109959.1 |
| 6.31684 | 0 | 0 | 0 | 9.33175 | 1 | ENSMUSG00000078891.4 |
| 0.665476 | 0.051013 | 0 | 0.005811 | 0.008018 | 1 | ENSMUSG00000097157.2 |
| 0.22125 | 0.060302 | 0.139809 | 0.165862 | 0.978243 | 1 | ENSMUSG00000097440.2 |
| 0.158825 | 0.25283 | 0.178113 | 0.491515 | 0.040432 | 1 | ENSMUSG00000096986.2 |
| 0.037315 | 0 | 0.055167 | 0 | 0.23468 | 1 | ENSMUSG00000085245.1 |
| 0 | 0 | 0 | 0 | 0 | 1 | ENSMUSG00000085634.1 |
| 0.04641 | 0.224158 | 0 | 0.074004 | 0.145615 | 1 | ENSMUSG00000074388.3 |
| 0.044322 | 0.059241 | 0.079502 | 0.020097 | 0 | 1 | ENSMUSG00000100037.1 |
| 0.035473 | 0.04592 | 0.014086 | 0.013494 | 0.018289 | 1 | ENSMUSG00000086968.8 |
| 0.20336 | 0.06016 | 0.085163 | 0.902554 | 0.051718 | 1 | ENSMUSG00000086712.2 |
| 0.072371 | 0.172607 | 0.172918 | 0.052827 | 0.114317 | 1 | ENSMUSG00000085723.2 |
| 0.033299 | 0.048757 | 0.065383 | 0.095473 | 0.043802 | 1 | ENSMUSG00000086587.7 |
| 1.60868 | 0.128031 | 0.035697 | 0.054233 | 0.380396 | 1 | ENSMUSG00000034764.15 |
| 0.579161 | 0.321779 | 0.195534 | 0.162565 | 0.118014 | 1 | ENSMUSG00000031736.11 |
| 0 | 0 | 0 | 0 | 0 | 1 | ENSMUSG00000085447.1 |
| 0 | 0 | 0 | 0 | 0 | 1 | ENSMUSG00000091351.1 |
| 0 | 0 | 0 | 0 | 0 | 1 | ENSMUSG00000102095.1 |
| 0 | 0 | 0 | 0 | 0 | 1 | ENSMUSG00000103647.1 |
| 0 | 0 | 0 | 0 | 0 | 1 | ENSMUSG00000087330.1 |
| 0.00279 | 0 | 0 | 0 | 0.00437 | 1 | ENSMUSG00000115207.1 |
| 0 | 0 | 0 | 0 | 0 | 1 | ENSMUSG00000114652.1 |
| 0.013453 | 0 | 0 | 0 | 0 | 1 | ENSMUSG00000114235.1 |
| 0 | 0 | 0 | 0 | 0 | 1 | ENSMUSG00000085118.2 |
| 0.003069 | 0.004495 | 0 | 0 | 0 | 1 | ENSMUSG00000112569.1 |
| 0.004953 | 0.016307 | 0.00925 | 0 | 0.019511 | 1 | ENSMUSG00000110017.1 |
| 0 | 0 | 0 | 0 | 0 | 1 | ENSMUSG00000107552.1 |
| 0 | 0.036054 | 0 | 0 | 0.571791 | 1 | ENSMUSG00000112261.1 |
| 0.01944 | 0 | 0.073835 | 0 | 0 | 1 | ENSMUSG00000087382.7 |
| 0.01473 | 0.014643 | 0.011606 | 0 | 0 | 1 | ENSMUSG00000102715.1 |
| 0.015727 | 0.022506 | 0 | 0.02564 | 0.336635 | 1 | ENSMUSG00000114422.1 |
| 0.036584 | 0 | 0 | 0 | 0.029694 | 1 | ENSMUSG00000090081.1 |
| 0.150839 | 0 | 0 | 0 | 0.205936 | 1 | ENSMUSG00000085558.7 |
| 0 | 0.011733 | 0 | 0 | 0.280733 | 1 | ENSMUSG00000097466.8 |
| 0.057749 | 0.023814 | 0.035742 | 0.035791 | 0.166094 | 1 | ENSMUSG00000104068.1 |
| 0 | 0 | 0 | 0 | 0 | 1 | ENSMUSG00000099599.1 |
| 0.012344 | 0 | 0 | 0 | 0 | 1 | ENSMUSG00000113960.1 |
| 0.024273 | 0 | 0 | 0 | 0 | 1 | ENSMUSG00000090307.7 |
| 0.013716 | 0.024657 | 0 | 0 | 0 | 1 | ENSMUSG00000112599.1 |
| 0 | 0 | 0 | 0 | 0 | 1 | ENSMUSG00000113309.1 |
| 0 | 0.019188 | 0 | 0.021666 | 0.022496 | 1 | ENSMUSG00000114980.1 |
| 0 | 0 | 0.214842 | 0 | 0 | 1 | ENSMUSG00000112097.1 |
| 0.051255 | 0 | 0 | 0 | 0.078663 | 1 | ENSMUSG00000084807.1 |
| 0 | 0.023325 | 0 | 0 | 0.102855 | 1 | ENSMUSG00000095369.1 |
| 0.035503 | 0 | 0.008411 | 0 | 0.022216 | 1 | ENSMUSG00000097358.1 |
| 0.229018 | 0.03895 | 0 | 0 | 0 | 1 | ENSMUSG00000087196.1 |
| 0.185613 | 0.007923 | 0 | 0.006398 | 0 | 1 | ENSMUSG00000085261.1 |
| 0.281417 | 0.071536 | 0 | 0.021861 | 0.015298 | 1 | ENSMUSG00000091985.1 |
| 0.285013 | 0.209898 | 0 | 0 | 0 | 1 | ENSMUSG00000112360.1 |
| 0.296089 | 0.271672 | 0 | 0.148825 | 0.287077 | 1 | ENSMUSG00000093622.1 |
| 0.568562 | 0.110998 | 0 | 0.383215 | 0.088756 | 1 | ENSMUSG00000111034.1 |
| 0.291468 | 0.079376 | 0.08195 | 0.045305 | 0 | 1 | ENSMUSG00000110235.1 |
| 0 | 0 | 0 | 1.0501 | 1.85507 | 1 | ENSMUSG00000085135.7 |
| 1.50117 | 0.315879 | 0.105671 | 0.029262 | 0.10545 | 1 | ENSMUSG00000102750.1 |

| | | | | | | |
|---|---|---|---|---|---|---|
| 1.20237 | 0.246778 | 0.066049 | 0.363251 | 0.105884 | 1 | ENSMUSG00000111535.1 |
| 2.1703 | 0 | 0 | 0 | 0 | 1 | ENSMUSG00000114884.1 |
| 1.78458 | 0.065039 | 0.123931 | 0.018866 | 0.013363 | 1 | ENSMUSG00000112095.1 |
| 1.43335 | 0.065842 | 0 | 0 | 0 | 1 | ENSMUSG00000103451.1 |
| 2.55875 | 0.068741 | 0.1342 | 0.14771 | 0.036785 | 1 | ENSMUSG00000053749.12 |
| 1.37072 | 0.654836 | 0.633697 | 0.752746 | 0 | 1 | ENSMUSG00000110593.1 |
| 1.27846 | 0 | 0 | 0.056698 | 0.051081 | 1 | ENSMUSG00000105632.1 |
| 59.4612 | 0.404095 | 0.064123 | 0.139304 | 0.129852 | 1 | ENSMUSG00000107355.1 |
| 2.86496 | 0.503297 | 0.906617 | 1.02926 | 0.385294 | 1 | ENSMUSG00000085360.1 |
| 1.18828 | 0 | 0.073388 | 0.167857 | 0.116953 | 1 | ENSMUSG00000097832.1 |
| 23.146 | 0.502448 | 0.361997 | 3.03012 | 2.84171 | 1 | ENSMUSG00000052248.15 |
| 0 | 2.08539 | 0 | 0 | 0 | 1 | ENSMUSG00000086517.1 |
| 0 | 1.94105 | 0 | 0 | 0.021418 | 1 | ENSMUSG00000084860.1 |
| 0 | 2.56993 | 0 | 0 | 0 | 1 | ENSMUSG00000086255.1 |
| 0 | 2.27331 | 0.066187 | 0 | 0.104136 | 1 | ENSMUSG00000085808.1 |
| 0 | 3.75442 | 0.084942 | 0 | 0.123901 | 1 | ENSMUSG00000111546.1 |
| 0 | 2.49981 | 0 | 0 | 0 | 1 | ENSMUSG00000084993.1 |
| 0 | 1.18924 | 0.148169 | 0 | 0 | 1 | ENSMUSG00000110766.1 |
| 0 | 1.123 | 0 | 0 | 0.228648 | 1 | ENSMUSG00000109827.1 |
| 0 | 1.0139 | 0 | 0 | 0.14022 | 1 | ENSMUSG00000106275.1 |
| 0 | 2.6793 | 0.601988 | 0 | 0.167691 | 1 | ENSMUSG00000110803.1 |
| 0.00735 | 0.080188 | 6.77646 | 0 | 0.013546 | 1 | ENSMUSG00000114390.1 |
| 0 | 0.065024 | 42.1512 | 0 | 0.160206 | 1 | ENSMUSG00000107266.1 |
| 0 | 0.820944 | 10.5809 | 0 | 0.02992 | 1 | ENSMUSG00000050179.3 |
| 0 | 0.394843 | 2.04468 | 0 | 0.017647 | 1 | ENSMUSG00000098108.7 |
| 0 | 0.239423 | 1.06953 | 0.050104 | 0.10938 | 1 | ENSMUSG00000104417.1 |
| 0 | 0.506913 | 2.44731 | 0 | 0.055243 | 1 | ENSMUSG00000112417.1 |
| 0 | 0.431845 | 2.59022 | 0.170657 | 0.131676 | 1 | ENSMUSG00000110656.1 |
| 0.011207 | 0.060543 | 1.16318 | 0.0046 | 0.06177 | 1 | ENSMUSG00000108132.1 |
| 0 | 2.66174 | 5.82217 | 0 | 0.174543 | 1 | ENSMUSG00000102280.1 |
| 0 | 0.931996 | 4.92686 | 0 | 1.32586 | 1 | ENSMUSG00000112230.1 |
| 0 | 0.035228 | 0 | 1.31523 | 0 | 1 | ENSMUSG00000108232.1 |
| 0 | 0 | 0 | 1.81794 | 0 | 1 | ENSMUSG00000087089.1 |
| 0 | 0.061259 | 0.013569 | 1.19324 | 0.070161 | 1 | ENSMUSG00000102785.5 |
| 0 | 0.022302 | 0 | 1.30193 | 0.032716 | 1 | ENSMUSG00000114306.1 |
| 0 | 0.087658 | 0.115492 | 11.6595 | 0.209201 | 1 | ENSMUSG00000104927.1 |
| 0 | 0.070897 | 0.007808 | 1.12657 | 0.08969 | 1 | ENSMUSG00000102882.1 |
| 0.005978 | 0.050385 | 0.028636 | 3.4988 | 0.15968 | 1 | ENSMUSG00000085844.1 |
| 0 | 0.17607 | 0 | 5.12298 | 0.073179 | 1 | ENSMUSG00000092591.1 |
| 0.020806 | 0.014819 | 0 | 1.29486 | 0.035996 | 1 | ENSMUSG00000084773.1 |
| 0 | 0.186873 | 0.050012 | 5.5244 | 0.251118 | 1 | ENSMUSG00000111839.1 |
| 0 | 0 | 0 | 0 | 1.50321 | 1 | ENSMUSG00000104077.1 |
| 0 | 0 | 0 | 0 | 3.50844 | 1 | ENSMUSG00000101264.1 |
| 0 | 0 | 0 | 0.055602 | 2.40062 | 1 | ENSMUSG00000105247.1 |
| 0.0207 | 0 | 0 | 0.032168 | 1.68304 | 1 | ENSMUSG00000104781.1 |
| 0 | 0 | 0.072855 | 0 | 1.37401 | 1 | ENSMUSG00000089755.1 |
| 0 | 0.065562 | 0.016982 | 0.048875 | 1.22434 | 1 | ENSMUSG00000087132.7 |
| 0 | 0.284772 | 0 | 0 | 1.62252 | 1 | ENSMUSG00000105963.1 |
| 0.019513 | 0.193947 | 0.203516 | 0.39752 | 12.5518 | 1 | ENSMUSG00000098292.1 |
| 0 | 0.422646 | 0 | 0.031425 | 2.9958 | 1 | ENSMUSG00000099343.1 |
| 0.029351 | 0.021484 | 0.052735 | 0.931161 | 10.3623 | 1 | ENSMUSG00000104340.1 |
| 0 | 0 | 0 | 0 | 0 | 3.50844 | 1 |
| 0.009179 | 0 | 0 | 0 | 0.055602 | 2.40062 | 1 |
| 0 | 0 | 0 | 0.072855 | 0 | 1.37401 | 1 |
| 0.055672 | 0 | 0 | 0 | 0.032168 | 1.68304 | 1 |
| 0.028074 | 0 | 0.065562 | 0.016982 | 0.048875 | 1.22434 | 1 |

| | | | | | | |
|---|---|---|---|---|---|---|
| 0.074129 | 0 | 0.284772 | 0 | 0 | 1.62252 | 1 |
| 0.050501 | 0 | 0.193947 | 0.203516 | 0.39752 | 12.5518 | 1 |
| 0 | 0.037614 | 0.422646 | 0 | 0.031425 | 2.9958 | 1 |
| 0.074954 | 0.030504 | 0.021484 | 0.052735 | 0.931161 | 10.3623 | 1 |

**Supplementary Table 2. Overlapping downregulated genes in MEP cells (203)**

| Gene ID | Gene Symbol |
|---|---|
| 71699 | Slc41a3 |
| 50997 | Mpp2 |
| 272396 | Tarsl2 |
| 107182 | Btaf1 |
| 16825 | Ldb1 |
| 66629 | Golph3 |
| 14081 | Acsl1 |
| 20341 | Selenbp1 |
| 17082 | Il1rl1 |
| 55934 | Rp9 |
| 72416 | Lrpprc |
| 13136 | Cd55 |
| 26556 | Homer1 |
| 14729 | Gp5 |
| 12725 | Clcn3 |
| 67383 | Carnmt1 |
| 72324 | Plxdc1 |
| 15451 | Hpn |
| 17427 | Mns1 |
| 15505 | Hsph1 |
| 208638 | Slc25a38 |
| 14735 | Gpc4 |
| 20910 | Stxbp1 |
| 194590 | Reps2 |
| 12593 | Cdyl |
| 72103 | Aplf |
| 21928 | Tnfaip2 |
| 75767 | Rab11fip1 |
| 74196 | Ttc27 |
| 218914 | Wapl |
| 21422 | Tfcp2 |
| 240283 | Dmxl1 |
| 233489 | Picalm |
| 72057 | Phf10 |
| 217935 | Dync2i1 |
| 207798 | Gramd1c |
| 70026 | Tspo2 |
| 14151 | Fech |
| 67886 | Camsap2 |
| 233280 | Nipa1 |
| 19165 | Psen2 |
| 66333 | Aqp11 |
| 17025 | Alad |
| 76857 | Spopl |
| 11555 | Adrb2 |
| 68337 | Crip2 |

| | |
|---|---|
| 52666 | Arhgef25 |
| 73046 | Glrx5 |
| 192657 | Ell2 |
| 270685 | Mthfd1l |
| 53620 | Vamp5 |
| 18174 | Slc11a2 |
| 68166 | Spire1 |
| 18441 | P2ry1 |
| 18263 | Odc1 |
| 68861 | Dipk2a |
| 15199 | Hebp1 |
| 224648 | Uhrf1bp1 |
| 30945 | Rnf19a |
| 53424 | Tsnax |
| 15510 | Hspd1 |
| 330119 | Adamts3 |
| 228012 | Tlk1 |
| 223752 | Gramd4 |
| 17330 | Minpp1 |
| 243574 | Kbtbd8 |
| 19159 | Cyth3 |
| 70617 | Fam241a |
| 11428 | Aco1 |
| 217410 | Trib2 |
| 226419 | Dyrk3 |
| 66350 | Pla2g12a |
| 14934 | Gypa |
| 66147 | Necap2 |
| 208846 | Daam1 |
| 11512 | Adcy6 |
| 244421 | Lonrf1 |
| 53945 | Slc40a1 |
| 232670 | Tspan33 |
| 69902 | Mrto4 |
| 56791 | Ube2l6 |
| 19895 | Rpia |
| 72692 | Hnrnpll |
| 74563 | Rasgef1c |
| 224860 | Plcl2 |
| 75552 | Paqr9 |
| 67667 | Alkbh8 |
| 72147 | Zbtb46 |
| 65973 | Asph |
| 320827 | Cracd |
| 12892 | Cpox |
| 380714 | Rph3al |
| 20454 | St3gal5 |
| 56222 | Cited4 |

| | |
|---|---|
| 67216 | Mboat2 |
| 50785 | Hs6st1 |
| 432628 | Mfsd2b |
| 16848 | Lfng |
| 58234 | Shank3 |
| 23970 | Pacsin2 |
| 17345 | Mki67 |
| 20751 | Spr |
| 338367 | Myo1d |
| 230935 | Dnajc11 |
| 20362 | Septin8 |
| 217125 | Samd14 |
| 211401 | Mtss1 |
| 12447 | Ccne1 |
| 20698 | Sphk1 |
| 193742 | Abhd16a |
| 14245 | Lpin1 |
| 13669 | Eif3a |
| 66870 | Serbp1 |
| 15525 | Hspa4 |
| 269152 | Kif26b |
| 50884 | Nckap1 |
| 11607 | Agtr1a |
| 15519 | Hsp90aa1 |
| 17191 | Mbd2 |
| 28109 | D10Wsu102e |
| 16905 | Lmna |
| 69632 | Arhgef12 |
| 338368 | Pheta2 |
| 19725 | Rfx2 |
| 69568 | Vkorc1l1 |
| 225995 | D030056L22Rik |
| 77053 | Sun1 |
| 233016 | Blvrb |
| 11979 | Atp7b |
| 93966 | Hemgn |
| 381126 | Garem1 |
| 11519 | Add2 |
| 192663 | Abcg4 |
| 225608 | Sh3tc2 |
| 73095 | Slc25a42 |
| 229279 | Hnrnpa3 |
| 76890 | Memo1 |
| 60455 | Pgap6 |
| 22427 | Wrn |
| 98766 | Ubac1 |
| 81535 | Sgpp1 |
| 75415 | Arhgap12 |

| | |
|---|---|
| 105787 | Prkaa1 |
| 67900 | Mtfp1 |
| 11861 | Arl4a |
| 19746 | Rhd |
| 13852 | Stx2 |
| 216781 | Trim58 |
| 20364 | Selenow |
| 14042 | Ext1 |
| 50496 | E2f6 |
| 72401 | Slc43a1 |
| 68428 | Steap3 |
| 54673 | Sh3glb1 |
| 16596 | Klf1 |
| 12521 | Cd82 |
| 217593 | Slc25a21 |
| 15288 | Hmbs |
| 78653 | Bola3 |
| 71684 | Rbm43 |
| 213539 | Bag2 |
| 57748 | Jmy |
| 216739 | Acsl6 |
| 207304 | Hectd1 |
| 109978 | Art4 |
| 66109 | Tspan13 |
| 56030 | Tmem131 |
| 266645 | Acmsd |
| 77579 | Myh10 |
| 84652 | Fam126a |
| 11804 | Aplp2 |
| 76522 | Lsm8 |
| 13143 | Dapk2 |
| 27410 | Abca3 |
| 104175 | Sbk1 |
| 109672 | Cyb5a |
| 27388 | Ptdss2 |
| 53417 | Hif3a |
| 20496 | Slc12a2 |
| 12831 | Col5a1 |
| 13857 | Epor |
| 56454 | Aldh18a1 |
| 237761 | Sowaha |
| 18027 | Nfia |
| 56473 | Fads2 |
| 69597 | Afg3l2 |
| 14630 | Gclm |
| 18844 | Plxna1 |
| 71648 | Optn |
| 12349 | Car2 |

| | |
|---|---|
| 20379 | Sfrp4 |
| 99887 | Tlcd4 |
| 12916 | Crem |
| 70122 | Mllt3 |
| 68646 | Nadk2 |
| 54616 | Extl3 |
| 109801 | Glo1 |
| 226747 | Ahctf1 |
| 71779 | Marchf8 |
| 56018 | Stard10 |
| 216705 | Clint1 |
| 57444 | Isg20 |
| 224454 | Zdhhc14 |

**Supplementary Table 3. Overlapping downregulated genes in MEL cells (75)**

Gm14327
Slc4a1
Gm5662
Csf2rb2
Alas2
Itgb3
Hbb-bs
Hbq1b
Hba-a1
Apol11b
Slamf1
E030030I06Rik
Csf2rb
Hbb-bt
Gm8995
Gda
Mrvi1
Oasl2
Ahnak
Rab3il1
Ptp4a3
Asic4
Gpsm2
Gstt1
Kalrn
Grap2
G430049J08Rik
Slfn5
Btg2
Anxa3
Rasl11a
Mybpc3
Usp18
Oas2
Itga2b
Serpina3g
Prkcq
Xaf1
Pf4
Oas3
Rsad2
Atp2b4
P2rx1
Scn10a
Gm6093
Plekho2
Cpeb4

Dnajb2
Gypa
Irgm1
Smox
Parp14
Oas1a
Adra2a
Ddx58
Pdcd1lg2
Isg15
Pim1
Redrum
Hipk2
Stat1
Hist1h1c
Cmpk2
Fam210b
Rhag
Col5a1
Cd24a
Med13l
Rap1b
Sla
Olfm1
St3gal1
Helz2
Prkar2b
Hba-x

#### Supplementary Table 4. Mass spectrometry analysis of lncEry interacting proteins in M

| Molecular w | Accession | Gene Name | CoverPercer | UniquePepC | UniquePeptide | Charge |
|---|---|---|---|---|---|---|
| 218.5 | E9QAS5 | Chd4 | 8.5848075 | 9 | | 218.5 |
| | | | | | GPFLVSAPLSTIINWER | 2 |
| | | | | | FNAPGAQQFCFLLSTR | 2 |
| | | | | | LLEQALVIEEQLR | 2 |
| | | | | | FSWAQGTDTILADEMGLGK | 2 |
| | | | | | CCNHPYLFPVAAMEAPK | 3 |
| | | | | | SSAQLLEDWGMEDIDHVFSEI | 3 |
| | | | | | FMFNIADGGFTELHSLWQNEI | 3 |
| | | | | | GGGNQVSLLNVVMDLK | 2 |
| | | | | | NQDETEDTELQGMNEYLSSFI | 2 |
| 181.8 | Q64511 | Top2b | 9.2431762 | 6 | | 181.8 |
| | | | | | GTIQELGQNQYAVSGEIFVVD | 2 |
| | | | | | LQTTLTCNSMVLFDHMGCLK | 3 |
| | | | | | GFQQISFVNSIATTK | 2 |
| | | | | | AASNCGIVESILNWVK | 2 |
| | | | | | VSIDPESNIISIWNNGK | 2 |
| | | | | | HGFLEEFITPIVK | 2 |
| 172.7 | Q01320 | Top2a | 21.989529 | 23 | | 172.7 |
| | | | | | YSGPEDDAAISLAFSK | 2 |
| | | | | | WEVCLTMSER | 2 |
| | | | | | KYDTVLDILR | 2 |
| | | | | | KEWLLGMLGAESSK | 2 |
| | | | | | GFQQISFVNSIATSK | 2 |
| | | | | | LLGLPEDYLYGQSTSYLTYNI | 2 |
| | | | | | IYVPALIFGQLLTSSNYDDDEI | 3 |
| | | | | | SYVDLYLK | 2 |
| | | | | | TLAVSGLGVVGR | 2 |
| | | | | | EWLLGMLGAESSK | 2 |
| | | | | | MELSPLQPVNENMLMNK | 2 |
| | | | | | QEIAFYSLPEFEEWK | 2 |
| | | | | | LLDGEEPLPMLPSYK | 2 |
| | | | | | NKQEIAFYSLPEFEEWK | 3 |
| | | | | | YIFTMLSPLAR | 2 |
| | | | | | FLEEFITPIVK | 2 |
| | | | | | VTIDPENNVISIWNNGK | 2 |
| | | | | | MQSLDKDIVALMVR | 2 |
| | | | | | AAIGCGIVESILNWVK | 2 |
| | | | | | IYVPALIFGQLLTSSNYDDDEI | 3 |
| | | | | | EDLAVFIEELEVVEAK | 2 |
| | | | | | SIPSMVDGLKPGQR | 2 |
| | | | | | TPSLITDYR | 2 |
| 105.9 | Q8K019 | Bclaf1 | 1.9586507 | 1 | | 105.9 |
| | | | | | STSESFIQHIVSLVHHVK | 3 |
| 95.3 | P58252 | Eef2 | 12.470862 | 6 | | 95.3 |
| | | | | | VFDAIMNFR | 2 |
| | | | | | IWCFGPDGTGPNILTDITK | 2 |
| | | | | | TFCQLILDPIFK | 2 |
| | | | | | AYLPVNESFGFTADLR | 2 |
| | | | | | LMEPIYLVEIQCPEQVVGGIY( | 3 |
| | | | | | YVEPIEDVPCGNIVGLVGVDQ | 3 |
| 87.9 | Q8VEK3 | Hnrnpu | 33.75 | 16 | | 87.9 |
| | | | | | MCLFAGFQR | 2 |
| | | | | | SSGPTSLFAVTVAPPGAR | 2 |
| | | | | | GNFTLPEVAECFDEITYVELQ | 3 |
| | | | | | YNILGTNTIMDK | 2 |
| | | | | | NGQDLGVAFK | 2 |
| | | | | | LSASSLTMESFAFLWAGGR | 2 |
| | | | | | LSASSLTMESFAFLWAGGR | 2 |
| | | | | | GYFEYIEENKYSR | 2 |
| | | | | | SPQPPVEEEDEHFDDTVVCLD | 4 |
| | | | | | DCEVVMMIGLPGAGK | 2 |
| | | | | | FDENDVITCFANFETDEVELS | 2 |
| | | | | | EVLADRPLFPHVLCHNCAVEI | 4 |

| | | | | | | |
|---|---|---|---|---|---|---|
| | | | | | NFILDQTNVSAAAQR | 2 |
| | | | | | KDCEVVMMIGLPGAGK | 3 |
| | | | | | EKPYFPIPEDCTFIQNVPLEDR | 3 |
| | | | | | GNFTLPEVAECFDEITYVELQ | 2 |
| | | | | | NQSQGYNQWQQGQFWGQKI | 4 |
| 69.2 | Q8BTS0 | Ddx5 | 32.520325 | 11 | | 69.2 |
| | | | | | QNFTEPTAIQAQGWPVALSGI | 3 |
| | | | | | GHNCPKPVLNFYEANFPANV | 4 |
| | | | | | TLSYLLPAIVHINHQPFLER | 3 |
| | | | | | WNLDELPK | 2 |
| | | | | | FVINYDYPNSSEDYIHR | 3 |
| | | | | | TGTAYTFFTPNNIK | 2 |
| | | | | | LLQLVEDR | 2 |
| | | | | | QVSDLISVLR | 2 |
| | | | | | LIDFLECGK | 2 |
| | | | | | TTYLVLDEADR | 2 |
| | | | | | DWVLNEFK | 2 |
| 66.9 | G5E924 | Hnrnpl | 31.056911 | 10 | | 66.9 |
| | | | | | QALVEFEDVLGACNAVNYAA | 3 |
| | | | | | ISRPGDSDDSR | 3 |
| | | | | | ASLNGADIYSGCCTLK | 2 |
| | | | | | NGVQAMVEFDSVQSAQR | 2 |
| | | | | | YGPQYGHPPPPPPPPDYGPHA | 4 |
| | | | | | AITHLNNNFMFGQK | 3 |
| | | | | | SSSGLLEWDSK | 2 |
| | | | | | SDALETLGFLNHYQMK | 3 |
| | | | | | VFNVFCLYGNVEK | 2 |
| | | | | | SKPGAAMVEMADGYAVDR | 3 |
| 60.9 | P63038 | Hspd1 | 6.6317627 | 2 | | 60.9 |
| | | | | | ALMLQGVDLLADAVAVTMG | 3 |
| | | | | | AAVEEGIVLGGGCALLR | 2 |
| 59.7 | G3UXA6 | Ptbp3 | 27.256318 | 6 | | 59.7 |
| | | | | | MDGVVTDLIAVGLK | 2 |
| | | | | | KIPCDVTEAEVISLGLPFGK | 3 |
| | | | | | SQAFLEMASEEAAVTMINYY | 3 |
| | | | | | ENALVQMADASQAQLAMNH | 3 |
| | | | | | AQAALQAVSAVQSGNLSLPG | 4 |
| | | | | | NNQFQALLQYADPVNAQYAI | 3 |
| 56.7 | E9Q6E5 | Srsf11 | 9.0019569 | 3 | | 56.7 |
| | | | | | ALIVVPYAEGVIPDETK | 2 |
| | | | | | TLFGFLGK | 2 |
| | | | | | FHDPDSAVVAQHLTNTVFVD | 3 |
| 38.2 | Q61990 | Pcbp2 | 44.751381 | 6 | | 38.2 |
| | | | | | LHQLAMQQSHFPMTHGNTGI | 5 |
| | | | | | GYWAGLDASAQTTSHELTIPI | 3 |
| | | | | | QVTITGSAASISLAQYLINVR | 2 |
| | | | | | IITLAGPTNAIFK | 2 |
| | | | | | AFAMIIDKLEEDISSSMTNSTA | 4 |
| | | | | | AFAMIIDKLEEDISSSMTNSTA | 4 |
| | | | | | AITIAGIPQSIIECVK | 2 |
| | | | | | ESTGAQVQVAGDMLPNSTER | 2 |
| 35.1 | Q8BFQ4 | Wdr82 | 6.3897764 | 1 | | 35.1 |
| | | | | | VVALSMSPVDDTFISGSLDK | 2 |
| 34.4 | P47962 | Rpl5 | 6.0606061 | 1 | | 34.4 |
| | | | | | VGLTNYAAAYCTGLLLAR | 2 |
| 34.3 | P35550 | Fbl | 35.474006 | 6 | | 34.3 |
| | | | | | LAAAILGGVDQIHIKPGAK | 3 |
| | | | | | VLYLGAASGTTVSHVSDIVGI | 4 |
| | | | | | MLIAMVDVIFADVAQPDQTR | 3 |
| | | | | | MQQENMKPQEQLTLEPYER | 3 |
| | | | | | DHAVVVGVYRPPPK | 3 |
| | | | | | TNIIPVLEDAR | 2 |
| 30.5 | Q9CX86 | Hnrnpa0 | 34.42623 | 6 | | 30.5 |
| | | | | | GFGFVYFQSHDAADK | 3 |
| | | | | | CFGFVTYSNVEEADAAMAAS | 3 |

| | | | | | | |
|---|---|---|---|---|---|---|
| | | | | | CFGFVTYSNVEEADAAMAAS | 4 |
| | | | | | GDVAEGDLIEHFSQFGAVEK | 3 |
| | | | | | LFIGGLNVQTSESGLR | 2 |
| | | | | | GHFEAFGTLTDCVVVVNPQT | 3 |
| 29.6 | P62702 | Rps4x | 23.954373 | 5 | | 29.6 |
| | | | | | VNDTIQIDLETGK | 2 |
| | | | | | TDITYPAGFMDVISIDK | 2 |
| | | | | | LTIAEER | 2 |
| | | | | | LSNIFVIGK | 2 |
| | | | | | FDTGNLCMVTGGANLGR | 2 |
| 26.7 | P62908 | Rps3 | 16.872428 | 3 | | 26.7 |
| | | | | | AELNEFLTR | 2 |
| | | | | | IMLPWDPSGK | 2 |
| | | | | | FVDGLMIHSGDPVNYYVDTA | 3 |

**MEP cells**

| XCorr C2 S( | MissCleavage |
|---|---|
| 19.48 | |
| 2.41 | 0 |
| 3.05 | 0 |
| 2.43 | 0 |
| 2.87 | 0 |
| 0.72 | 0 |
| 3.86 | 0 |
| 4.86 | 0 |
| 1.54 | 0 |
| 0.87 | 0 |
| 19.57 | |
| 2.03 | 0 |
| 1.06 | 0 |
| 1.02 | 0 |
| 3.61 | 0 |
| 1.8 | 0 |
| 2.31 | 0 |
| 97.96 | |
| 1.26 | 0 |
| 1.99 | 0 |
| 2.69 | 1 |
| 2.59 | 1 |
| 2.62 | 0 |
| 4.14 | 0 |
| 5.35 | 1 |
| 1.25 | 0 |
| 1.88 | 0 |
| 3.29 | 0 |
| 1.54 | 0 |
| 2.68 | 0 |
| 2.18 | 0 |
| 3.76 | 1 |
| 2.34 | 0 |
| 3.14 | 0 |
| 2.56 | 0 |
| 3.04 | 1 |
| 4.21 | 0 |
| 2.19 | 0 |
| 4.09 | 0 |
| 0.24 | 0 |
| 0.25 | 0 |
| 3.05 | |
| 3.05 | 0 |
| 7.55 | |
| 1.38 | 0 |
| 1.4 | 0 |
| 2.99 | 0 |
| 2.22 | 0 |
| 2.34 | 0 |
| 1.66 | 0 |
| 112.57 | |
| 2.08 | 0 |
| 2.46 | 0 |
| 6.43 | 1 |
| 2.92 | 0 |
| 0.83 | 0 |
| 4.33 | 0 |
| 1.71 | 0 |
| 3.01 | 1 |
| 3.67 | 0 |
| 3.01 | 0 |
| 3.86 | 0 |
| 9.13 | 0 |

| | |
|---|---|
| 3.58 | 0 |
| 2.99 | 1 |
| 6.65 | 0 |
| 4.46 | 0 |
| 7 | 0 |
| 51.19 | |
| 7.1 | 0 |
| 3.28 | 0 |
| 6.12 | 0 |
| 1.9 | 0 |
| 3.66 | 0 |
| 1.63 | 0 |
| 0.77 | 0 |
| 2.06 | 0 |
| 1.55 | 0 |
| 1.15 | 0 |
| 1.84 | 0 |
| 58.49 | |
| 8.94 | 0 |
| 0.25 | 0 |
| 0.24 | 0 |
| 3.75 | 0 |
| 8.91 | 0 |
| 2.3 | 0 |
| 2.82 | 0 |
| 5.02 | 0 |
| 3.61 | 0 |
| 0.65 | 0 |
| 2.18 | |
| 2.18 | 0 |
| 0.49 | 0 |
| 25.3 | |
| 2.88 | 0 |
| 3.07 | 1 |
| 3.21 | 0 |
| 4.7 | 0 |
| 3.5 | 0 |
| 2.41 | 0 |
| 2.32 | |
| 2.32 | 0 |
| 1.5 | 0 |
| 0.5 | 0 |
| 48.56 | |
| 1.45 | 0 |
| 6.8 | 0 |
| 6.99 | 0 |
| 2.91 | 0 |
| 3.91 | 1 |
| 6.08 | 1 |
| 4.1 | 0 |
| 0.25 | 0 |
| 2.55 | |
| 2.55 | 0 |
| 2.06 | |
| 2.06 | 0 |
| 11.33 | |
| 4.06 | 0 |
| 4.72 | 0 |
| 2.55 | 0 |
| 1.24 | 0 |
| 1.29 | 0 |
| 0.85 | 0 |
| 43 | |
| 2.58 | 0 |
| 7.91 | 0 |

| | |
|---|---|
| 7 | 1 |
| 5.7 | 0 |
| 2.55 | 0 |
| 6.03 | 0 |
| 8.2 | |
| 0.59 | 0 |
| 3.9 | 0 |
| 0.63 | 0 |
| 1.24 | 0 |
| 2.54 | 0 |
| 11.23 | |
| 1.24 | 0 |
| 2 | 0 |
| 4.83 | 0 |

**Supplementary Table 5. Mass spectrometry analysis of lncEry interacting proteins in M**

| Molecular w | Accession | Gene Name | CoverPercei | UniquePepC | UniquePeptide | Charge |
|---|---|---|---|---|---|---|
| 332 | E9Q557 | Dsp | 0.0453 | 24 | | 332908.42 |
| | | | | | R.AESGPDLR.Y | 2 |
| | | | | | R.VVIVDPETNK.E | 2 |
| | | | | | K.AISVPR.V | 2 |
| | | | | | K.DQDITR.I | 2 |
| | | | | | K.FLDQNLQK.Y | 2 |
| | | | | | K.GLQDSIR.K | 2 |
| | | | | | K.IEIER.R ! K.IEIER.L ! K.IELE | 2 |
| | | | | | K.IIEYK.R ! K.LIEYK.T | 2 |
| | | | | | K.NDLNLK.K | 2 |
| | | | | | K.QEAFSIR.M | 2 |
| | | | | | K.VIETNR.E | 2 |
| | | | | | K.VYEAR.L ! K.VYEAR.A | 2 |
| | | | | | R.AQIDNLTR.E | 2 |
| | | | | | R.ESLLVK.I | 2 |
| | | | | | R.LPVEEAYK.R | 2 |
| | | | | | R.LQAEIK.R ! K.LQAELK.I | 2 |
| | | | | | R.QVQNLVNK.S | 2 |
| | | | | | R.TQEELR.E ! R.TQEELR.R | 2 |
| | | | | | K.VYEAR.L ! K.VYEAR.A | 2 |
| | | | | | K.VYEAR.L ! K.VYEAR.A | 2 |
| | | | | | K.IEVLEEELR.L | 2 |
| | | | | | R.ETQSQLESER#.C | 2 |
| | | | | | R.GLVGIEFK.E | 2 |
| | | | | | R.YIELLTR.S | 2 |
| 144.78196 | Q9WTL4 | Insrr | 0.0054 | 2 | | 144873.13 |
| | | | | | R.SEVTELR.R | 2 |
| | | | | | R.SEVTELR.R | 2 |
| 69.246811 | Q61656 | Ddx5 | 0.0342 | 3 | | 69289.45 |
| | | | | | K.APILIATDVASR.G | 2 |
| | | | | | R.EANQAINPK.L | 2 |
| | | | | | R.TAQEVDTYR.R | 2 |
| 60.917393 | P63038 | Hspd1 | 0.0279 | 2 | | 60954.79 |
| | | | | | K.DGVTVAK.S | 2 |
| | | | | | R.VTDALNATR.A | 2 |
| 35.1 | Q8BFQ4 | Wdr82 | 0.0224 | 4 | | 35078.65 |
| | | | | | K.VAVLDGK.H | 2 |
| | | | | | K.IYDLR.G ! K.IYDLR.K ! K.L' | 2 |
| | | | | | K.YGVDLIR.Y | 2 |
| | | | | | R.LIDAFK.G | 2 |
| 34.285675 | P35550 | Fbl | 0.0245 | 1 | | 34306.38 |
| | | | | | R.GGGGGGFR.G | 2 |
| 33.3 | Q61990 | Pcbp2 | 0.0829 | 3 | | 38221.27 |
| | | | | | K.EVGSIIGK.K | 2 |
| | | | | | K.IANPVEGSTDR.Q | 2 |
| | | | | | R.INISEGNCPER.I | 2 |
| 32.25939 | P17225 | Ptbp1 | 0.038 | 4 | | 56477.46 |
| | | | | | K.FGTVLK.I | 2 |
| | | | | | R.LTSLNVK.W ! K.LTSLNVK.' | 2 |
| | | | | | R.EGQEDQGLTK.D | 2 |
| | | | | | R.SAGVPSR.V ! R.SAGVPSR.- | 2 |
| 30.511733 | Q9CX86 | Hnrnpa0 | 0.059 | 2 | | 30530.04 |
| | | | | | K.EDIHAGGGGAR.A | 3 |

| | | | | | | |
|---|---|---|---|---|---|---|
| | | | | | K.IFVGGIK.E ! K.LFVGGIK.E ! | 2 |
| 22.875049 | P97461 | Rps5 | 0.1373 | 5 | | 22889.13 |
| | | | | | K.AQCPIVER.L | 2 |
| | | | | | K.GSSNSYAIK.K | 2 |
| | | | | | R.KAQCPIVER.L | 3 |
| | | | | | R.QAVDVSPLR.R | 2 |
| | | | | | R.QAVDVSPLRR.V | 3 |
| 15.787748 | P61358 | Rpl27 | 0.125 | 2 | | 15797.53 |
| | | | | | K.VVLVLAGR.Y | 2 |
| | | | | | R.YSVDIPLDK.T | 2 |
| 14.418419 | P11031 | Sub1 | 0.2047 | 3 | | 14427.29 |
| | | | | | K.EQISDIDDAVR.K | 2 |
| | | | | | K.ILIDIR.E ! K.LIIDIR.S | 2 |
| | | | | | K.QAVPEKPVK.K | 3 |
| 26.7 | P62908 | Rps3 | 0.2345 | 5 | | 47668.1 |
| | | | | | R.ELAEDGYSGVEVR.V | 2 |
| | | | | | K.AELNEFLTR.E | 2 |
| | | | | | K.GCEVVVSGK.L | 2 |
| | | | | | R.ELAEDGYSGVEVR.V | 2 |
| | | | | | R.QGVLGIK.V | 2 |
| 29.6 | P62702 | Rps4x | 0.2471 | 9 | | 59160.73 |
| | | | | | K.DANGNSFATR.L | 2 |
| | | | | | K.IFVGTK.G | 2 |
| | | | | | K.TGENFR.L | 2 |
| | | | | | R.IGVITNR.E | 2 |
| | | | | | R.LIYDTK.G | 2 |
| | | | | | R.YPDPLIK.V | 2 |
| | | | | | R.IGVITNR.E | 2 |
| | | | | | K.YALTGDEVK.K | 2 |
| 95.3 | O08796 | Eef2 | 0.0291 | 4 | | 190595.96 |
| | | | | | R.FTDTR.K | 2 |
| | | | | | K.GEGQLSAAER.A | 2 |
| | | | | | R.CITIK.S ! R.CLTLK.H ! K.CL | 2 |
| | | | | | R.FTDTR.K | 2 |
| 21.8 | Q6PDM2 | Srsf1 | 0.1613 | 5 | | 55488.5 |
| | | | | | R.DAEDAVYGR.D | 2 |
| | | | | | R.EAGDVCYADVYR.D | 2 |
| | | | | | R.VEFPR.T ! R.VEFPR.S | 2 |
| | | | | | R.DAEDAVYGR.D | 2 |
| | | | | | R.DIDLK.N ! R.DIDLK.R ! R.DI | 2 |
| 15.3 | P63038 | Hspd1 | 0.065 | 2 | | 76225.32 |
| | | | | | R.VTDALNATR.A | 2 |
| | | | | | K.DGVTVAK.S | 2 |
| 18 | P47962 | Rpl5 | 0.064 | 3 | | 34400.26 |
| | | | | | K.VFGALK.G | 2 |
| | | | | | R.ENPVYEK.K | 2 |
| | | | | | R.LVIQDK.N | 2 |

**MEL cells**

| Score | MissCleavage |
|---|---|
| 24.09 | 0 |
| 20.15 | 0 |
| 29.72 | 0 |
| 40.37 | 0 |
| 20.57 | 0 |
| 25.86 | 0 |
| 24.43 | 0 |
| 24 | 0 |
| 26.04 | 0 |
| 30.17 | 0 |
| 24.2 | 0 |
| 25.16 | 0 |
| 21.54 | 0 |
| 22.27 | 0 |
| 26.02 | 0 |
| 27.9 | 0 |
| 22.24 | 0 |
| 24.31 | 0 |
| 25.29 | 0 |
| 23.69 | 0 |
| 24.74 | 0 |
| 31.29 | 0 |
| 26.63 | 0 |
| 27.49 | 0 |
| 21.5 | 0 |
| 23.2 | 0 |
| 45.52 | 0 |
| 26.1 | 0 |
| 40.44 | 0 |
| 21.51 | 0 |
| 33.63 | 0 |
| 42.46 | 0 |
| 20.6 | 0 |
| 24.3 | 0 |
| 30.02 | 0 |
| 40.71 | 0 |
| 27.03 | 0 |
| 37.16 | 0 |
| 46.49 | 0 |
| 27.54 | 0 |
| 20.05 | 0 |
| 33.1 | 0 |
| 32.61 | 0 |
| 44.79 | 0 |

| | |
|---|---|
| 38.71 | 0 |
| 44.68 | 0 |
| 35.17 | 0 |
| 30.03 | 1 |
| 36.23 | 0 |
| 23.94 | 1 |
| 44 | 0 |
| 31.71 | 0 |
| 61.11 | 0 |
| 24.7 | 0 |
| 27.49 | 0 |
| 28.87 | 0 |
| 25.63 | 0 |
| 40.68 | 0 |
| 51.78 | 0 |
| 20.33 | 0 |
| 27.31 | 0 |
| 28.42 | 0 |
| 46.18 | 0 |
| 49.16 | 0 |
| 30.32 | 0 |
| 22.77 | 0 |
| 42.51 | 0 |
| 20.16 | 0 |
| 23.17 | 0 |
| 28.86 | 0 |
| 22.35 | 0 |
| 31.49 | 0 |
| 56.08 | 0 |
| 28.47 | 0 |
| 24.84 | 0 |
| 55.92 | 0 |
| 22.98 | 0 |
| 33.63 | 0 |
| 21.51 | 0 |
| 22.23 | 0 |
| 25.13 | 0 |
| 22.53 | 0 |

**Supplementary Table 6.**

**5' and 3' RACE Primers**

| | |
|---|---|
| 5' RACE | GATTACGCCAAGCTT GAAAGTGACCGGATGCTGAT |
| 3'RACE | GATTACGCCAAGCTT CTGAGTTCCACAGTGCAGTATC |

Note: Red color indicates the targeting sequence against the corresponding genes.

**qPCR Primers**

| Genes | Sequences |
|---|---|
| *lncEry* | F: TATCCTGCTGTTGCTGGGTC |
| | R: CCCCCAAGCATCATTGTCCT |
| *lncEry-3* | F: AACAGTATGGAAATGCTTGC |
| | R: TTTCAGTCTATCCCAGACAG |
| *Ntn4* | F: GCAGGCTTGAATGGAGTAGC |
| | R: CCCGGAGCTTTCTTCCCAAA |
| *Gapdh* | F: TGGCCTTCCGTGTTCCTAC |
| | R: GAGTTGCTGTTGAAGTCGCA |
| *mU1* | F: CTGGCAGGGGAGATACCATG |
| | R: AGTCGAGTTTCCCGCATTTG |
| *Elk3* | F: TCCTCACGCGGTAGAGATCAG |
| | R: GTGGAGGTACTCGTTGCGG |
| *Lta4h* | F: GAGGTCGCGGATACTTGCTC |
| | R: CTCCTGTGACTGGACCGTG |
| *Hal* | F: CTGTGCGACGCTACATGAAGA |
| | R: TCATTGTCCTCTAAGGCCACC |
| *Ccdc38* | F: AACCACAGCATGAAAATCTACCA |
| | R: AGCCTGAGAATAAGAGTGGGAT |
| *Usp44* | F: ATGGATAGGTGCAAGCACGTT |
| | R: GCTCTTGGATGTACTTCCCACAG |
| *Metap2* | F: TTCGGGGGACACCTGAATG |
| | R: TGTTTTGCTACTTCATCCACCAA |

| | |
|---|---|
| *Vezt* | F: TGTAGACCCAGTGGAATCTGTAA |
| | R: ACACACATACTCAGTCCTACTGT |
| *Fgd6* | F: CTCCCCCTCCTATTGCACCTA |
| | R: GTCGGGACTTTTGGTTTTGGC |
| *Klf1* | F: GGCGAACTTTGGCACCTAAGA |
| | R: AGAAGGGACGATGTCCAGTGT |
| *Fech* | F: TGGAGAGAGATGGACTAGAGAGG |
| | R: CCACCTGTCGATTGTGCTC |
| *Ldb1* | F: CATTGGCCGGACCCTGATAC |
| | R: CGAGGGACACGAAGTTGCT |
| *Rhd* | F: GATCACAGGTTCATGGCGAG |
| | R: GTCTCCGAAAGGACGAGGAC |
| *Car2* | F: TCCCACCACTGGGGATACAG |
| | R: CTCTTGGACGCAGCTTTATCATA |
| *Ptp4a3* | F: CATCACTGTTGTGGACTGGC |
| | R: TGGATGGCGTCCTCGTACTT |
| *Rhag* | F: AGCCTGGAAGTTGCTATGATTG |
| | R: ACTTGAGGTGTTACCCTGCTG |
| *Tal1* | F: CACTAGGCAGTGGGTTCTTTG |
| | R: GGTGTGAGGACCATCAGAAATCT |
| *Alas2* | F: TGGGCTAAGAGCCATTGTCCT |
| | R: GTAGGTGTGGTCCTGTTTCTTC |
| *Csf2rb2* | F: TCCAGCCAGATCGTGACCT |
| | R: AATCCCCAAGAGATACACTCCA |
| *Csf2rb* | F: AAAAACAGCCAGTGTCCTGTG |
| | R: GATGCTGACGTTCTTGGGAAG |
| *CD24a* | F: GTTGCACCGTTTCCCGGTAA |
| | R: CCCCTCTGGTGGTAGCGTTA |
| *Gata2* | F: GCCGGGAGTGTGTCAACTG |

| | |
|---|---|
| | R: AGGTGGTGGTTGTCGTCTGA |
| *Gata1* | F: TGGGGACCTCAGAACCCTTG |
| | R: GGCTGCATTTGGGGAAGTG |
| *Wdr82* | F: GACCTCATCAGATACACCCATGC |
| | R: GTCATGCAAGGACAAGTAACGAA |
| *α-globin* | F: CACCACCAAGACCTACTTCC |
| | R: CAGTGGCTCAGGAGCTTGA |
| *β-globin* | F: TTTAACGATGGCCTGAATCACTT |
| | R: CAGCACAATCACGATCATATTGC |

**qChIP Primers**

| Genes | Sequences |
|---|---|
| *Hba-a1* | F: GTGTGTCCGGGCAACTGATA |
| | R: TGGAGCAGTTCTCATTGGCT |
| *Hba-a2* | F: GGGTAAGAAGTGTGTCCGGG |
| | R: GCCCTTGGAGCAGTTCTCAT |
| *Hbb-b1* | F: AAGGTGAACGCCGATGAAGT |
| | R: CCTGCTGGTAAGCAGACCTC |
| *Hbb-b2* | F: AGCAGGGTCAGTTGCTTCTT |
| | R: ACCAACTTCATCGGCGTTCA |

**siRNA Sequences**

| siRNAs | Sequences |
|---|---|
| Control | UUCUCCGAACGUGUCACGUdTdT |
| | ACGUGACACGUUCGGAGAAdTdT |
| lncEry-1 | GCCAAGAUCAUCACGAGUAdTdT |
| | UACUCGUGAUGAUCUUGGCdTdT |
| lncEry-2 | GGAGACACCUUAAUCUGAUdTdT |
| | AUCAGAUUAAGGUGUCUCCdTdT |
| lncEry-3 | CCUGAUGUCCAAAGGCCUUdTdT |
| | AAGGCCUUUGGACAUCAGGdTdT |
| Wdr82-1 | GCAGCCAACACAGUCGUUUdTdT |
| | AAACGACUGUGUUGGCUGCdTdT |
| Wdr82-2 | GCAAACCUGUCUGUUCCUUdTdT |
| | AAGGAACAGACAGGUUUGCdTdT |
| Wdr82-3 | GGUCAAACUUUAUGACCUUdTdT |
| | AAGGUCAUAAAGUUUGACCdTdT |

**shRNA Sequences**

| | |
|---|---|
| Control | CCGGTTCTCCGAACGTGTCACGTCTCGAGACGTGACACGTTCGGAGAATTTTTG |
| lncEry-1 | CCGGGAGGAGACACCTTAATCTGATCTCGAGATCAGATTAAGGTGTCTCCTCTTTTTG |
| lncEry-2 | CCGGCGCCTGATGTCCAAAGGCCTTCTCGAGAAGGCCTTTGGACATCAGGCGTTTTTG |
| Wdr82-1 | CCGGGAACGGAGAGAGTGGTATAAACTCGAGTTTATACCACTCTCTCCGTTCTTTTTG |
| Wdr82-2 | CCGGTTTGATCCAGAAGGGTTAATTCTCGAGAATTAACCCTTCTGGATCAAATTTTTG |

| | |
|---|---|
| Wdr82-3 | CCGGTGCAGCCAACACAGTCGTTTACTCGAGTAAACGACTGTGTTGGCTGCATTTTTG |
| Ddx5-1 | CCGGCGGGAAGCTAATCAAGCAATTCTCGAGAATTGCTTGATTAGCTTCCCGTTTTTG |
| Ddx5-2 | CCGGGCACTCAGCAATATGGAAGTACTCGAGTACTTCCATATTGCTGAGTGCTTTTTG |
| Ddx5-3 | CCGGGCGAATGTCATGGATGTGATTCTCGAGAATCACATCCATGACATTCGCTTTTTG |

Note: Red color indicates the targeting sequence against the corresponding genes.

#### ChIRP probe Sequences

| Probes | Sequences |
|---|---|
| Lac Z-1 | 5'- ccagtgaatccgtaatcatg-3' |
| Lac Z-2 | 5'-ataatttcaccgccgaaagg-3' |
| Lac Z-3 | 5'-aaacggggatactgacgaaa-3' |
| lncEry-1 | 5'-agatgttttcagtctatccc-3' |
| lncEry-2 | 5'-ccaggacttttgaagcatat-3' |
| lncEry-3 | 5'-ggactggctgtttatctgaa-3' |

#### Luciferase Reporter Primers

| Genes | Sequences |
|---|---|
| *Hba-a1* | F: CGGGGTACCGACCAACAATACCTCTTTCC |
| | R: CGACGCGTGGTTTCTTCCTGAGTCTGTC |
| *Hba-a2* | F: CGGGGTACCCTCAGACTTGTAACACAGC |
| | R: CGACGCGTCGGAGACAAAGTGGACAC |
| *Hbb-b1* | F: CGGGGTACCGTCAGGATGTAAGAACCAGAG |
| | R: CGACGCGTGATGTCTGTTTCTGAGGTTGC |
| *Hbb-b2* | F: CGGGGTACCGCAGGACCACTGACAGAC |
| | R: CGACGCGTGATGTCTGTTTCTGGGGTTG |
